# Influence of condensation domains on activity and specificity of adenylation domains

**DOI:** 10.1101/2021.08.23.457306

**Authors:** Janik Kranz, Sebastian L. Wenski, Alexander A. Dichter, Helge B. Bode, Kenan A. J. Bozhüyük

## Abstract

Many clinically used natural products are produced by non-ribosomal peptide synthetases (NRPSs), which due to their modular nature should be accessible to modification and engineering approaches. While the adenylation domain (A) plays the key role in substrate recognition and activation, the condensation domain (C) which is responsible for substrate linkage and stereochemical filtering recently became the subject of debate - with its attributed role as a “gatekeeper” being called into question. Since we have thoroughly investigated different combinations of C-A didomains in a series of *in vitro*, *in vivo*, and *in situ* experiments suggesting an important role to the C-A interface for the activity and specificity of the downstream A domain and not the C domain as such, we would like to contribute to this discussion. The role of the C-A interface, termed ‘extended gatekeeping’, due to structural features of the C domains, can also be transferred to other NRPSs by engineering, was finally investigated and characterised in an *in silico* approach on 30 wild-type and recombinant C-A interfaces. With these data, we not only would like to offer a new perspective on the specificity of C domains, but also to revise our previously established NRPS engineering and construction rules.

## Introduction

Peptide drugs like penicillins (antibiotic) (***Bills and Gloer, 2016***), cyclosporin (immunosuppressant) (***Velkov et al., 2011***), and bleomycin (anti-cancer) (***Du et al., 2000***) shaped our lives in an unprecedented way. They not only make a prodigious contribution to our public health by curing us from live threatening and formally untreatable diseases but, most of these scaffolds also share a common mode of synthesis (***Newman and Cragg, 2020***). They are complex specialised metabolites (SMs) predominantly synthesised by bacteria and fungi via biosynthetic pathways independent of the ribosome, denoted as Non-Ribosomal Peptide Synthetases (NRPSs) (***Felnagle et al., 2008***). NRPSs are large, multifunctional (mega-) enzymes in which multiple, repeating modules of enzymatic domains catalyse the incorporation and programmed functional group modifications of selected extender units into the growing peptide chain (Fig.1) (***Süssmuth and Mainz, 2017***).

**Figure 1.**
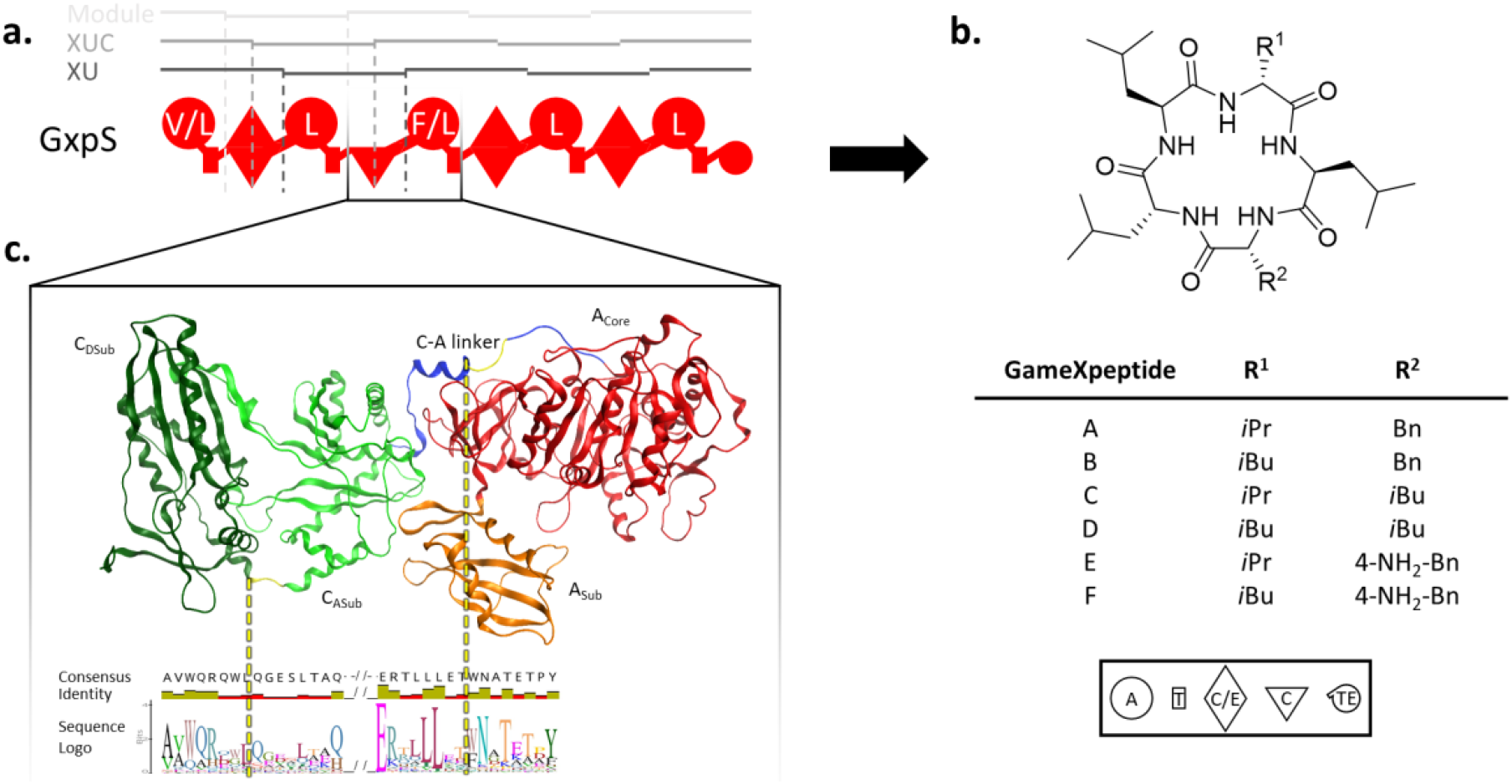
Outline of the NRPS and its declared units with the example of the GxpS. (**a**) Schematic representation of the GameXPeptide Synthetase with modules, XUs and the XUCs highlighted. The domains are illustrated by the following symbols: adenylation (A) domain, large circle; thiolation (T) domain, small rectangle; condensation (C) domain, triangle; dual condensation/epimerizazion (C/E) domain, diamond; thioesterase (TE) domain, small circle. Further editing domains like epimerization (E) domains, C- or N-methylation by methyltransferase (MT) domains or the redox state through redox-active (Ox, Red) domains are not depicted here. The standard one letter AA nomenclature is used to show the substrate acceptance. (**b**) Structure of the produced GameXPeptides (***Nollmann et al., 2015***) on the top with the varying residues R^1^ and R^2^ listed on the bottom. (**c**) Schematic representation of the C-A didomain is illustrated in ribbon representation by the SrfA-C termination module from *Bacillus subtilis* ATCC 21332 (PDB ID: 2VSQ) (***Tanovic et al., 2008***). The C domains’ *N*-terminal donor C_DSub_ (dark green) and *C*-terminal acceptor C_ASub_ (light green) site, the C-A linker (blue) and the A domain with its larger A_Core_ (red) and smaller A_Sub_ (orange) are depicted. The fusion sites of the XUC and XU marked with dashed lines (yellow) directing to their exact position in the consensus logo in an alignment below. **Supplementary Information – Table S1**

Biosynthesis of non-ribosomal peptides (NRPs) is likened to assembly-line processes (Fig.1) and dependent upon the activity and precise interplay of at least three ‘core’ domains: An Adenylation (A) domain for the selection and activation of extender units, i.e., amino acids; a Thiolation (T) domain, carrying a post translationally attached prosthetic 4’-phosphopantetheine (4ʹ-PPant) group, onto which the activated substrate is covalently attached to; and a Condensation (C) domain, covalently linking the T domain bound substrates to the growing peptidyl chain (***Sieber and Marahiel, 2005***). However, to develop novel drug entities by rationally modifying NRPSs, i.e., by altering the resulting peptides’ length and/or composition to improve drug likeness properties, bioavailability, or to overcome resistance mechanisms, understanding the inherent logic of NRP assembly is of utter importance (***Alanjary et al., 2019***).

Nowadays, crystal structure data and much of the fundamental biochemistry of all essential catalytic domains and domain complexes are available (***Süssmuth and Mainz, 2017***). For example, pioneering work on A domains not only revealed the first solved NRPS domain structure (PheA, PDB: 1AMU) (***Conti et al., 1997***), but that NRP synthesis is initiated by specific recognition and activation of the cognate substrate(s) by the A domain (***Stachelhaus et al., 1999***). After binding of the relevant dedicated amino acid from a pool of substrates by the A domain, substrate activation is achieved in a two-step chemical reaction. First, the A domain catalyses the formation of an aminoacyl adenylate intermediate using Mg^2+^–ATP consumption and release of PP_i_ (***Reimer et al., 2018***; ***Tanovic et al., 2008***). Second, the obtained amino acid – O – AMP anhydride is converted into a covalently bound thioester by a nucleophilic attack of the free thiol – 4’-PPant cofactor of the adjacent T domain (***Drake et al., 2016***; ***Gulick, 2009***). These findings, in turn, have inspired early efforts to rationally re-programme assembly-lines to produce tailor-made molecules by targeted mutagenesis of the A domains’ specificity conferring active site residues (***Eppelmann et al., 2002***; ***Schneider et al., 1998***; ***Thirlway et al., 2012***), swapping A domains (***Crüsemann et al., 2013***; ***Kries et al., 2015***), A-T or C-A di-domains (***Duerfahrt et al., 2003***; ***Stachelhaus et al., 1995***), and whole modules (C-A-T tri-domains) (***Baltz, 2014***; ***Mootz et al., 2000***)– but with limited success, indicating that further proofreading mechanisms or gatekeeping domains may be encoded within the assembly-line to ensure biosynthesis of the desired product(s).

Further gatekeeping functions are attributed to C domains (^L^C_L_, ^D^C_L_, and C/E; superscript: stereochemistry of the *C*-terminal residue of the donor substrate, subscript: stereochemistry of the acceptor substrate, C/E: dual C domain that catalyses both, epimerization and condensation) (***Belshaw et al., 1999***; ***Rausch et al., 2007***), which typically accept two T domain-bound substrates and catalyse peptide bond formation through the attack of the downstream acceptor substrate upon the thioester of the upstream donor substrate (***Finking and Marahiel, 2004***). Structural and biochemical characterizations disclosed that C domains have a pseudo-dimeric V-shaped structure with a *N*- and *C*-terminal subdomain (Fig. 1c) (***Keating et al., 2002***). Together, both subdomains are forming two opposite tunnels that lead from the donor-T and acceptor-T domain binding sites to the conserved key catalytic-residues containing active site motif HHxxxDG (***De Crécy-Lagard et al., 1995***; ***Izoré et al., 2021***; ***Keating et al., 2002***; ***Süssmuth and Mainz, 2017***). Very early on, biochemical characterizations showed that C domains exhibit a strong stereochemical selectivity for the donor-T domain bound substrate (**<u>^L^</u>**C_L_, **<u>^D^</u>**C_L_) and a significant side-chain selectivity for the acceptor-T domain bound substrate (^L^C**<u>_L_</u>**, ^D^C**<u>_L_</u>**) (***Belshaw et al., 1999***; ***Linne and Marahiel, 2000***). Nevertheless, the exact role and especially how C domains contribute to determining NRPS specificity is still unclear and subject to debate (***Baunach et al., 2021***; ***Calcott et al., 2020***; ***Izoré et al., 2021***).

Until recently, however, state-of-the-art NRPS engineering strategies assumed that the interface formed by C and A domains functions as a stable platform which should not be separated (***Tanovic et al., 2008***). Thus, the ascribed substrate specificity of the C domains could be neglected for the substrate bound to the acceptor-T domain for more than a decade (***Bozhüyük et al., 2019b***). This changed with the introduction of the eXchange Unit (XU) concept – a rule-based mix-and-match strategy to reproducibly engineer NRPSs (***Bozhüyük et al., 2018***). This concept uses A-T-C tri-domains, denoted as XUs, that can be fused within the C-A linker regions (Fig. 1c). In addition to breaking the dogma of the inseparability of the C-A interface, another important aspect of this concept is the recommendation that the substrate specificity of the corresponding C domains must be respected to obtain catalytically active chimeric-NRPSs – as was evident from literature and experimental data at the time. Thus, the XU concept has subsequently been improved even further to overcome observed substrate incompatibility issues by dividing C domains within the flexible linker that connects the *N*- (C_DSub_) and *C*-terminal (C_ASub_) subdomains (Fig. 1c), yielding the so-called eXchange Unit Condensation domain (XUC) concept (***Bozhüyük et al., 2019a***). Although both, the XU and XUC strategies allowed these assembly lines to be functionally reprogrammed with great efficiency, there is a growing body of evidence that, in particular, the attributed strong selectivity of C domains for the acceptor-T domain bound substrate is likely to be the exception rather than the rule (***Baunach et al., 2021***; ***Calcott et al., 2020***).

A number of recent insightful studies have led to results that at least question the “proof-reading” role of C domains during NRP synthesis for legitimate reasons. In a nutshell, recent studies suggest that: (I) C and A domains do not co-evolve (***Baunach et al., 2021***); (II) recombination within A domains are the main drivers of natural product diversification (***Baunach et al., 2021***; ***Booth et al., 2021***); (III) the C-A linker region contributes to A domain substrate specificity and activity (***Calcott et al., 2020***); and (IV) recent structural data found that C domains do not have a distinct pocket to select the acceptor-T domain bound side chain during peptide assembly, but that residues within the active site motif may instead serve to tune substrate selectivity (***Izoré et al., 2021***).

Herein, we sought to contribute to the controversially discussed matter of C domain specificity and whether C domain selectivity is indeed just a presumption that has unnecessarily complicated rational NRPS redesign – as most recently suggested (***Calcott et al., 2020***). For us, who introduced the XU and XUC concepts and thus contributed to the rise of the potentially false dogma, the answer to this question is of great importance. It is imperative to prevent C domain specificity from becoming a false dogma that influences future engineering efforts in the wrong way, as the NRPS community has experienced before with the falsely assumed inseparability of C-A di-domains (***Brown et al., 2018***).

With this in mind, we reviewed recombinant NRPSs created in our lab to identify functional artificial BGCs showing an unexpected behaviour, like C domains accepting noncognate substrates or altered A domain activation profiles not matching the profiles observed in the natural context (***Bozhüyük et al., 2018***; ***Bozhüyük et al., 2019a***; ***Bozhüyük et al., 2021***). Indeed, the examples identified do not support the idea that C domains generally have strict selectivity but can at least accept a range of substrates with similar physicochemical properties - supporting insights obtained from the latest solved crystal structure data (***Izoré et al., 2021***). Nevertheless, especially in the presence of promiscuous A domains, we occasionally observed changes in the substrate activation profiles or the preference of an alternative substrate over the WT substrate observed *in situ*. These observations suggest either that C domains do have some kind of gatekeeping function and thus favour certain substrates over others, or that the C domains themselves are able to tune the activity and specificity of the downstream A domains – as also reported earlier (***Meyer et al., 2016***). To shed further light on the role of C domains on NRP synthesis, we systematically analysed the effect of C domains onto A domains via a series of *in vitro*, *in vivo*, *in situ*, and *in silico* characterizations.

## Results

To quickly grasp the influence of C domains on the activity and selectivity of A-domains, we initially took advantage of the GameXPeptide (GXP) A-F producing Synthetase (GxpS) from *Photorhabdus laumondii subsp. laumondii* TT01 (***Nollmann et al., 2015***). GxpS, besides being the most widespread BGC in *Photorhabdus* and *Xenorhabdus* strains. (***Shi and Bode, 2018***), is one of our best studied, most engineered, and most promiscuous model systems (***Bian et al., 2015***; ***Bozhüyük et al., 2018***; ***Bozhüyük et al., 2019a***; ***Bozhüyük et al., 2021***) – producing a library of cyclic penta-peptides (**1-4**) (***Nollmann et al., 2015***). This library of peptide derivatives is synthesized due to the relaxed selectivity of the A domains from modules 1 (A1: leucine & valine) and 3 (A3: *p*-NH_2_-phenylalanine, phenylalanine, leucine). In addition, the latter has already been characterised *in vitro* and *in vivo* in previous work (***Bian et al., 2015***; ***Bozhüyük et al., 2019a***). In the course of these characterisations, it was even possible to determine that the GxpS_A3, in addition to a broad variety of proteinogenic amino acids (*in vitro*), recognizes and activates non-natural *para*-(*p*), *meta*- (*m*), and *ortho*- (*o*) substituted amino acids (*in vitro* and *in vivo*), such as *m*/*o*/*p*-Cl-Phe, *m*/*o*/*p*-F-Phe, *m*/*p*-Br-Phe, and *p*-O(C_3_H_3_)-Phe (***Bozhüyük et al., 2019a***). Since this promiscuity is the ideal prerequisite to analyse the influence of C domains on the activity profile of A domains, we selected GxpS_A3 as a first framework for further experiments.

### *In vitro* characterisations highlight influence of C domains on GxpS_A3’s activity and selectivity

To get first biochemical evidence of the hypothesized influence of C domains on A domain selectivity we cloned, heterologously produced (in *E. coli* BL21 (DE3) Gold), purified (via His6-Tag affinity chromatography), and *in vitro* assayed three GxpS derived proteins (P1: GxpS_A3-T3; P2: GxpS_C_ASub_-A3-T3; and P3: GxpS_C3-A3-T3) against all 20 proteinogenic amino acids in the presence or absence of an upstream domain (C or C_Asub_) (Fig. 2). For better comparability of the results, we chose two different *in vitro* assays for adenylation activity. On the one hand the ‘traditional’ γ-[^18^O_4_]-ATP pyrophosphate exchange assay (Fig. 2a) (***Phelan et al., 2009***) and on the other hand the recently introduced multiplexed hydroxamate assay (HAMA) (Fig. 2b-c) (***Stanišić et al., 2019***). Whereas the γ-[^18^O_4_]-ATP targets the first half-reaction of amino acid activation, detecting the isotopic back exchange of unlabelled PP_i_ into γ-^18^O_4_-labelled ATP and is analysed by MALDI/HRMS (***Phelan et al., 2009***), the HAMA assay targets the second half-reaction, quenching the formed aminoacyl adenylate by adding hydroxylamine and the resulting amino-hydroxamates are analysed by tandem mass spectrometry (MS/MS) analysis (***Stanišić et al., 2019***). Another major difference is the number of substrates that can be tested in one reaction. In contrast to the γ-[^18^O_4_]-ATP isotope exchange assay, which only allows testing of one substrate per reaction, the HAMA assay allows the parallel testing of dozens of competing amino acid substrates. Therefore, the HAMA assay is supposed to mimic the natural conditions in the cell much better, as all substrates are present at the same time.

**Figure 2.**
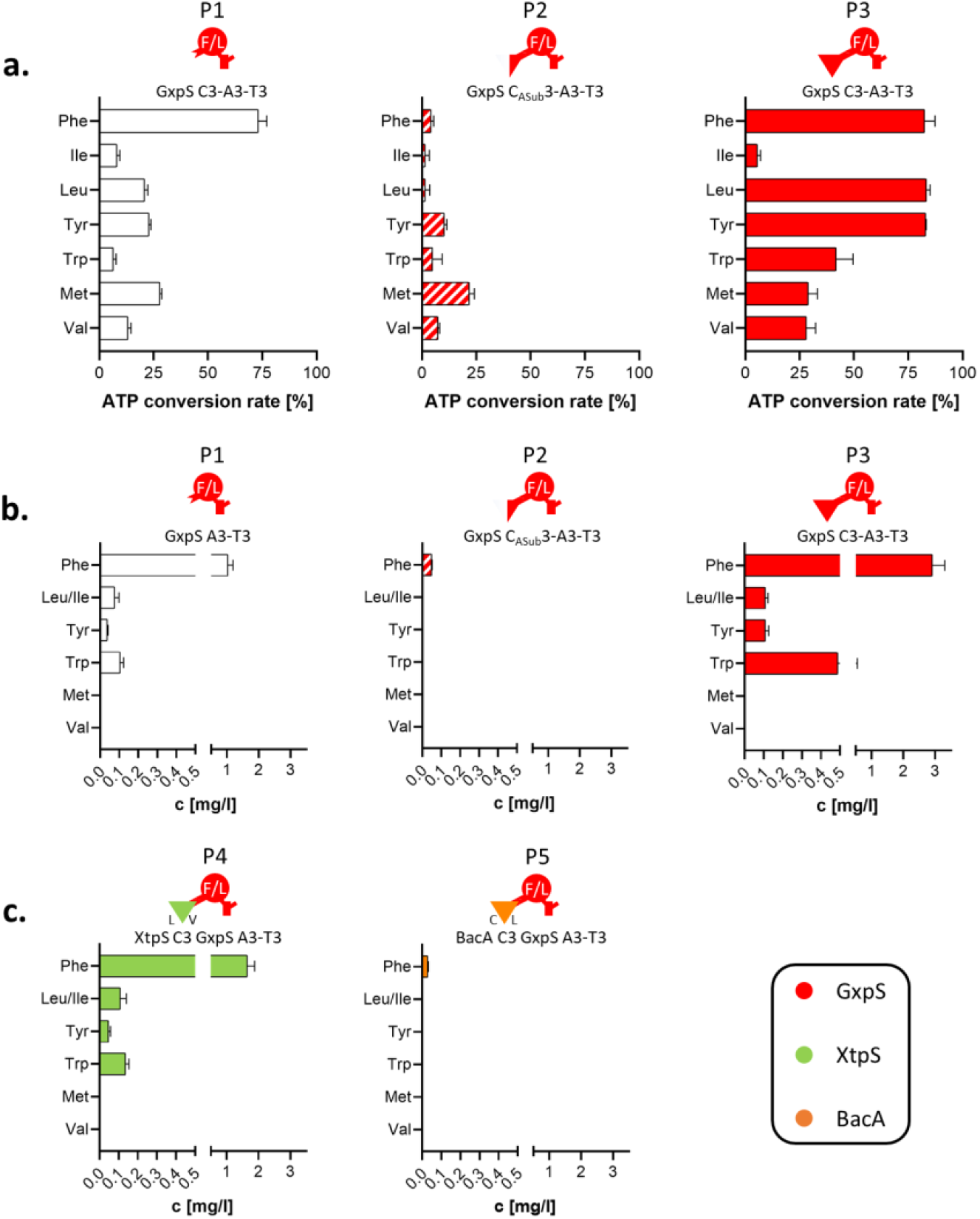
*In vitro* characterization of the GxpS A3. (**a**) GxpS A3-T3 with no, GxpS_C3_ASub_, GxpS_C3 or XtpS_C3 domain tested in a γ-[^18^O_4_]-ATP assay for ATP conversion rate measured with MALDI/HRMS; (**b**) GxpS_A3-T3 with no, GxpS_C3_ASub_, GxpS_C3 or XtpS_C3 domain tested in a HAMA for produced peptide yields measured with HPLC/HRMS; (**c**) BacA_C3 GxpS_A3-T3 tested in a HAMA for produced peptide yields measured with HPLC/HRMS. The representation of the NRPS domains by symbols is according to Fig. 1, and C_DSub_ and C_ASub_ are labelled corresponding to the preferred up- and down-stream A domain substrate in WT NRPS. **Supplementary Information – Table S1** **Supplementary Information – Table S5** **Supplementary Information – Table S6** **Supplementary Information – Table S8**

As a result of this first *in vitro* characterization of P1 - P3, it can be stated that the presence and absence of any domain (GxpS_C_Asub_3, GxpS_C3, XtpS_C3) upstream of GxpS_A3 showed a great influence on adenylation activities and substrate recognition profiles in both assays – with notable differences, though (Fig. 2). In general, P1 - P3 showed a much broader capacity to activate different substrates in the γ-[^18^O_4_]-ATP isotope exchange assay than in the HAMA assay. For instance, P1 showed adenylation activities against all 20 substrates in the γ-[^18^O_4_]-ATP isotope exchange assay with a higher preference for non-polar aromatic amino acids (Tab. S5), whereas when assayed with the HAMA assay only 5 substrates were activated (Phe, Trp, Leu/Ile, Tyr), more closely resembling the A domain’s *in vivo* behaviour (Tab. S6). As expected, however, P1 showed highest specificity for phenylalanine in both assays, and in addition good ATP conversion rates at ∼25 % for methionine, tyrosine, and leucine in the γ-[^18^O_4_]-ATP isotope exchange assay. P2, carrying the *C*- terminal subdomain of GxpS_C3 upstream of GxpS_A3, showed impaired catalytic potential to activate the offered substrates in both assays. Of note, with very different activation and specificity profiles depending on the assay chosen, i.e., P2 favoured methionine in the γ-[^18^O_4_]-ATP isotope exchange assay and phenylalanine in the HAMA assay. In contrast, for P3, carrying the full length GxpS_C3 domain, we observed an improved catalytic efficiency to activate the offered substrates. P3 revealed an almost identical activation profile as P1 in the HAMA assay, but with almost three-fold increased turnover rates, in the γ-[^18^O_4_]-ATP isotope exchange assay P3 showed highest ATP conversion rates (∼80 %) for phenylalanine, leucine and tyrosine (Fig. 2a-b).

In sum, these results are indicative for the importance of a functional C-A didomain interface for the activity and specificity of A domains. The gathered *in vitro* data of both assays along with insights from recent literature data (***Baunach et al., 2021***; ***Booth et al., 2021***; ***Calcott et al., 2020***; ***Izoré et al., 2021***; ***Stanišić et al., 2021***) provide evidence for an extended gatekeeping function for the C domains upstream of A domains rather than strict intrinsic selectivity. Similar results have also been reported previously for mono- and multi-specific modules that either strictly incorporate leucine or arginine or incorporate chemically diverse amino acids in parallel into microcystin (***Meyer et al., 2016***). Interestingly, in this study, the presence of the C domain’s *C*- terminal subdomain, including all C-A interface-forming residues, was sufficient to restore, at least in part, the specificities observed *in vivo*, whereas in our case, the presence of the *C*-terminal subdomain (Fig. 2a-b) even impaired the A domains capacity to efficiently recognise the substrates presented.

Next, and to better understand the influence of C-A interfaces on substrate recognition and activation of adjacent A domains, we targeted P1 by creating two chimeric proteins with three domains each (P4 & P5) that were analysed via the HAMA assay (Fig. 2b- c). While P4 was generated by fusion of the C3 domain of the xenotetrapeptide- producing synthetase (XtpS) from the Gram-negative *Xenorhabdus nematophila* ATCC 19061 (***Kegler et al., 2014***), which is very similar to the originally present GxpS_C3 (∼86 % sequence identity) (Tab. S8), P5 was generated by fusion of the C2 domain of the peptide antibiotic-producing bacitracin synthetase (BacA) from the unrelated Gram-positive *Bacillus licheniformis* ATCC 10716 (***Konz et al., 1997***) (∼44 % sequence identity) (Tab. S8). Both hybrid proteins – as well as all hybrid constructs described below – were created according to the splicing position established within the XU concept (***Bozhüyük et al., 2018***). As expected, P4 showed a very similar activity and amino acid recognition profile to P3, with phenylalanine being the preferred substrate (Fig. 2a-b). In contrast, P5 almost completely lost its catalytic activity, with a barely detectable signal for phenylalanine left (Fig. 2c), confirming that altered C-A interactions do have a great impact on the A domains capacity to recognise and activate respective substrates, indeed.

### Varying C domains result in altered *in vivo* product spectra

As *in vitro* experiments sometimes can lead to results not reflecting the enzymes true *in vivo* behaviour, for example, as experienced with results from biochemical characterisations of C domains (***Belshaw et al., 1999***; ***Dekimpe and Masschelein, 2021***; ***Linne and Marahiel, 2000***; ***Rausch et al., 2007***; ***Stanišić et al., 2021***), we also performed a series of *in vivo* experiments with truncated chimeric GxpS versions. GxpS has the rare potential to *in vivo* initiate peptide synthesis even after deletion of the initiation module (***Bozhüyük et al., 2021***) – as was also recently reported for the teicoplanin-producing NRPS in an *in vitro* study (***Kaniusaite et al., 2020***). However, for our experimental setup, we deleted the first two modules (A1 to C3) of GxpS, inserted either none (NRPS-1), GxpS_C_ASub_3 (NRPS-2), GxpS_C3 (NRPS-3) (Fig. 3a) or related C domains (63 to 69 % sequence identity) (Tab. S8) of various origins with different ascribed acceptor site specificities (NRPS-4 to -8) (Fig. 3a-b), produced the resulting NRPSs heterologously in *E. coli* DH10B::*mtaA* (***Schimming et al., 2014***), and analysed the culture extracts by HPLC-MS/MS. Throughout the present work, the resulting peptides and yields were confirmed by HPLC-MS/MS (Tab. S7) and comparison of retention times with synthetic standards (see Supplementary Information).

**Figure 3.**
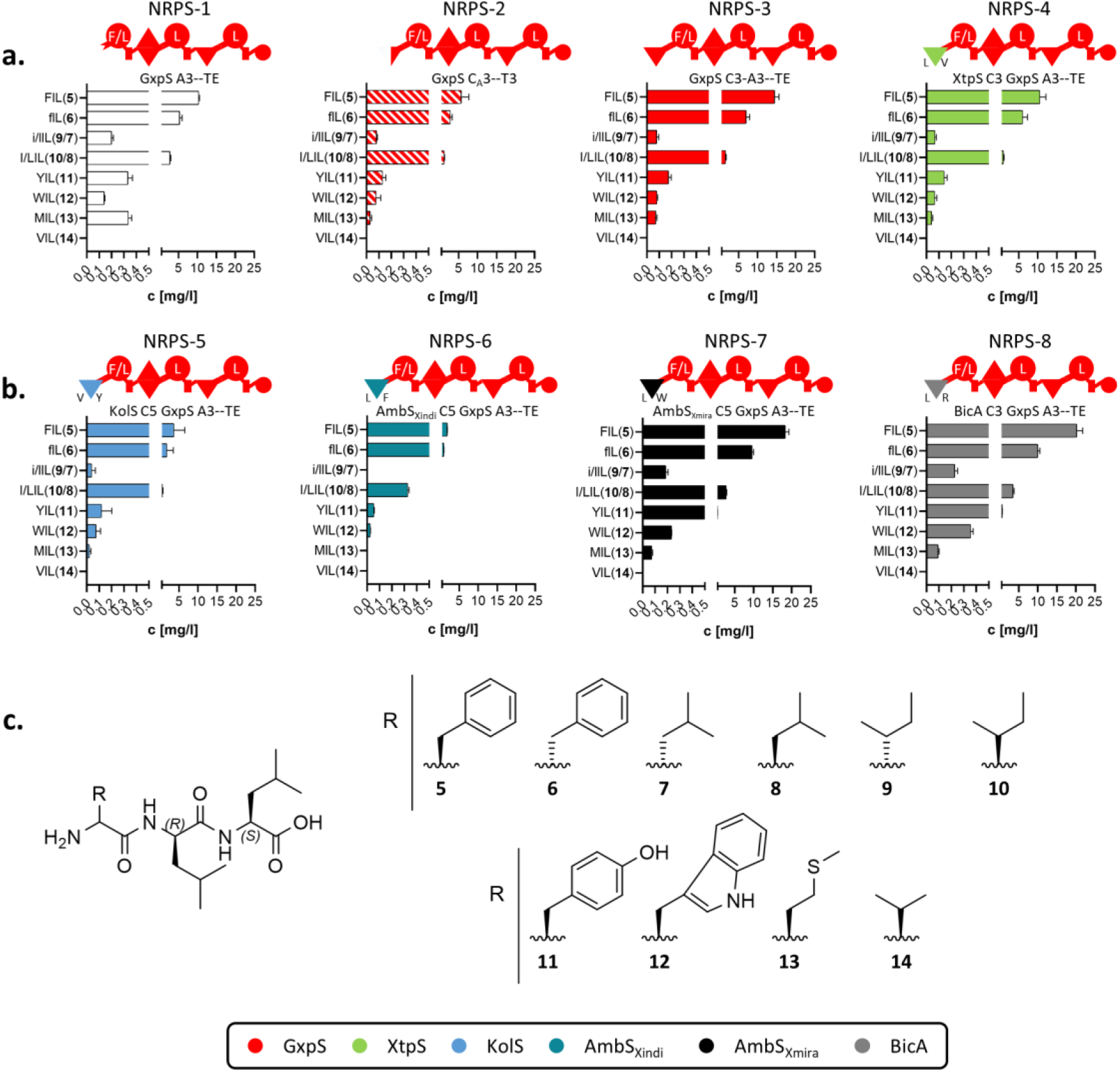
*In vivo* characterization of the GxpS A3 with varying C domains using truncated GxpS versions. (**a**) terminal GxpS_A3--TE with no, GxpS_C3_ASub_, GxpS_C3 or XtpS_C3 domain heterologous expressed in *E. coli* DH10B::*mtaA* and the extracts were measured via HPLC/MS; (**b**) terminal GxpS A3--TE with KolS_C5, BicA_C3, AmbS_Xmira__C5 or AmbS_Xindi__C5 domain heterologous expressed in *E. coli* DH10B::*mtaA* and the extracts were measured via HPLC/MS. The representation of the NRPS domains by symbols is according to Fig. 1, and C_DSub_ and C_ASub_ are labelled corresponding to the preferred up- and down-stream A domain substrate in WT NRPS; (**c**) Compounds **5** - **14** produced from NRPS-1 to -8 expressed in *E. coli* DH10B::*mtaA* and the extracts were measured via HPLC/MS. **Supplementary Information – Table S1** **Supplementary Information – Table S7** **Supplementary Information – Table S8**

Briefly, all of them, NRPS-1 to -8, were catalytically active showing biosynthesis of the same range of tripeptides (**5** - **14**), due to the promiscuity of GxpS_A3 – with FlL (**5**) having highest titres followed by flL (**6**) and l/LlL (**7** & **8**) (Fig. 3a-b). Interestingly, despite the C/E domain downstream of GxpS_A3, all peptides are produced with higher titers towards the L-configuration (**5**, **8,** & **10**). As the epimerization reaction is reversible and finds its end in the adjustment of an equilibrium between both isomers (***Stachelhaus and Walsh, 2000***), this might indicate that the downstream C/E domain, which usually expects a peptidyl chain, is unable to make sufficient contact with the activated amino acid, resulting in delayed condensation followed by late thiolation. Hence, this change in the reaction velocity caused by the length of the substrate (***Stein et al., 2005***) might lead to the observed diastereomer with a trend towards the L-isomer and not the expected D-isomer. However, titres of NRPS-2 are slightly lower but NRPS-3s’ are ∼30 % higher compared to NRPS-1 (Fig. 3a) – confirming the *in vitro* observed influence of the C-A interface on general biocatalytic activity of A domains (Fig. 2a-b). Remarkably, GxpS_A3 as part of NRPS-1, -2, and -3 showed a different substrate activation profile than as part of P1, P2, and P3, respectively. NRPS-1, -2, and -3 mainly synthesised the tripeptides **5** to **10**, known and expected from WT behaviour, illustrating why biochemical *in vitro* characterisations must always be treated with the utmost caution, especially with regard to multi-modular assembly lines.

Compared to NRPS-1, the chimeric proteins NRPS-4 to -8 showed no difference in the number of substrates activated by GxpS_A3, but the overall peptide production rates of NRPS-5 and -6 were ∼50 % lower and of NRPS-7 and -8 ∼200 % higher (Fig. 3b). For example, the latter produces **11** with 2-fold higher yield than NRPS-1 and 4-fold higher than NRPS-3. Consequently, NRPS-1 to -8 are supporting the *in vitro* observed extended gatekeeping function of C domains (Fig. 2) by fine tuning the A domains’ substrate selectivity.

### In situ recombination shows evidence for C domains’ extended gatekeeping function

To further investigate the influence of altered C-A interactions on the product spectra of NRPSs, we next sought to study homologous BGCs present in several bacterial strains producing the same peptide scaffold but resulting in different derivatives. The great advantage of such highly homologous systems is the possibility to study the effect of altered C-A interactions without having to consider further potential incompatibilities that could inhibit synthesis. We therefore targeted the fabclavine- producing BGCs (*fcl*; Fig. 4a) present in several *Xenorhabdus* strains (***Wenski et al., 2019***), including *X. budapestensis* DSM 16342 (*Xbud*), *X. hominickii* 17903 (*Xhom*), and *X. szentirmaii* DSM 16338 (*Xsze*), which were studied in present work.

**Figure 4.**
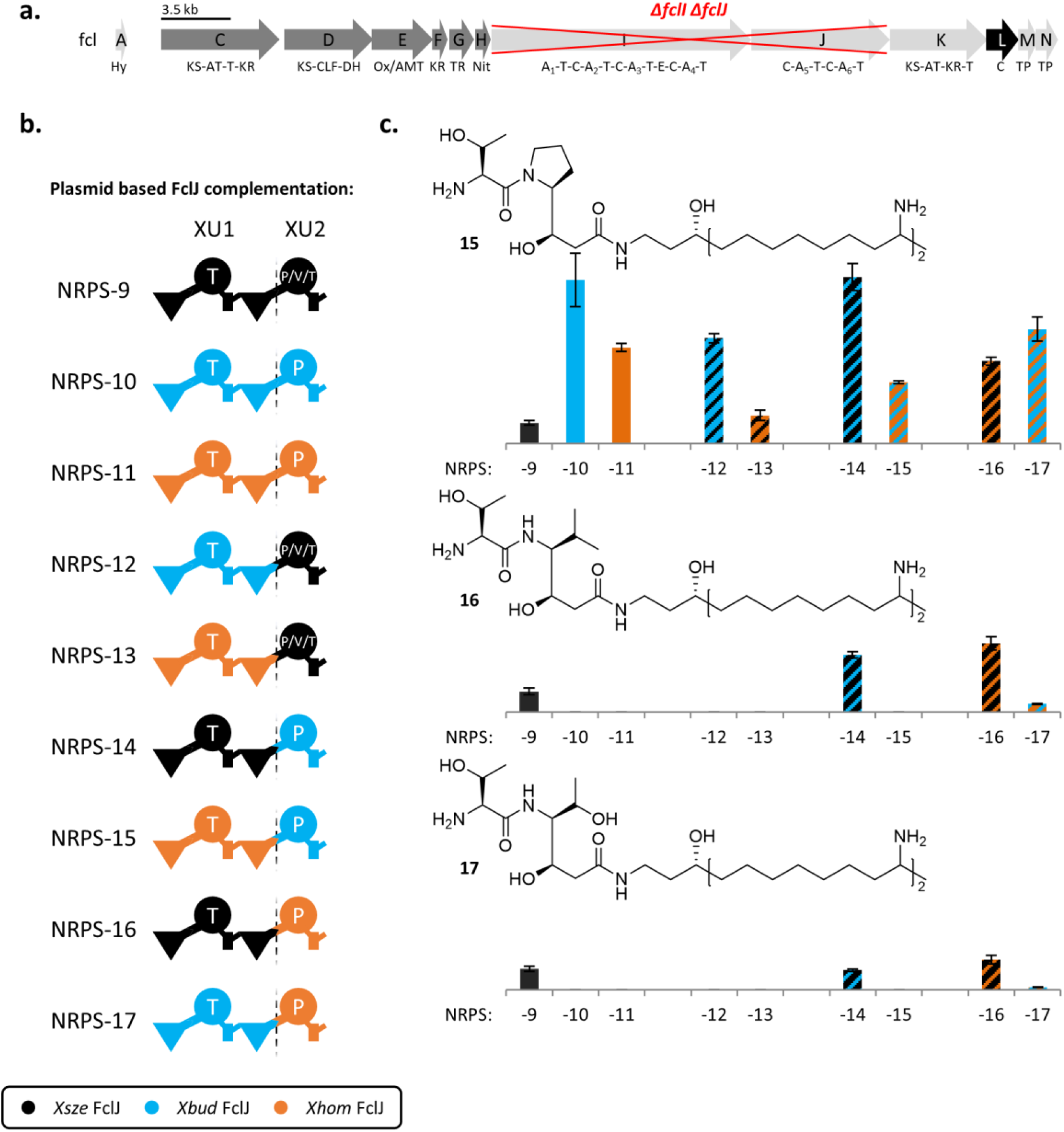
Fabclavines and the plasmid-based XU combinations for *fclJ* complementation in *X. szentirmaii ΔfclIJ*. (**a**) Fabclavine biosynthesis gene cluster (BGC) with the *ΔfclI ΔfclJ* deletion marked in red. (**b**) Schematic representation of the carried-out XU combinations of the *Xsze* FclJ_C5--C6/A6-T6 (black), *Xbud* FclJ C5-- C6/A6-T6 (light blue), and *Xhom* FclJ C5--C6/A6-T6 (orange). The representation of the NRPS domains by symbols is according to Fig. 1. (**c**) Hatched bar charts with corresponding colour code of the plasmid based FclJ insertions of the Pro derivative (Top), Val derivative (Middle), and the Thr derivative (Bottom). The produced quantity of each product **15**, **16** or **17** was compared in percentage relative to the amount of produced **15**, **16** or **17** by *Xsze* FclJ_C5--C6/A6-T6 (set as 100 %). **Supplementary Information – Table S1** **Supplementary Information – Table S8** **Supplementary Information – Table S9** **Supplementary Information – Figure S1** **Supplementary Information – Figure S2** **Supplementary Information – Figure S3**

Fabclavines are bioactive peptide-polyketide-polyamine hybrids with broad-spectrum activity against bacteria, fungi, and other eukaryotic cells (Fig. 4a). (***Donmez Ozkan et al., 2019***; ***Fuchs et al., 2014***; ***Wenski et al., 2019***) In previous work, the deletion of *fclI* of the NRPS encoding genes *fclIJ* led to shortened polyamine carrying fabclavine derivatives (**15** - **17**), and thus to the assumption that FclJ can also be used as a starting unit without FclI (***Wenski et al., 2019***). Accordingly, and due to *fclJ*’s rather small size, encoding two NRPS elongation modules (∼7 kbp), FclJ was chosen as promising starting point to investigate C-A interface substitutions *in situ*. FclJ, however, bears another advantage necessary to study the impact of an altered C-A interface on the A domain’s substrate recognition profile – namely FclJ_A6. While this particular A domain recognises and activates proline in *X. budapestensis* and *X. hominickii*, it also activates threonine and valine in *X. szentirmaii* (***Wenski et al., 2020***). As the promiscuity of the latter neither can be explained by differences within the respective proteins’ primary structure (∼89 % similarity) (Tab. S8) nor with changes within the substrate specificity conferring amino acid residues within the A domains active site (Tab. S9), we hypothesised that the respective upstream C domain (FclJ_C6) must be the reason for the product diversification in *X. szentirmaii* (Fig. 4).

To explore the substrate promiscuity of FclJ_A6 in *X. szentirmaii*, we generated a *X. szentirmaii ΔfclI ΔfclJ* double knockout mutant and prepared a library of plasmids encoding WT FclJ from *X. szentirmaii* (NRPS-9), *X. budapestensis* (NRPS-10) and *X. hominickii* (NRPS-11), as well as six chimeric FclJ combinations (Fig. 4b, NRPS-12 to -17) for plasmid-based gene complementation experiments (Fig. 4b). These six chimeric NRPSs represent all possible XU-C-A interface combinations from the chosen set of target-BGCs and therefore allows us to investigate whether the observed promiscuity of FclL_A6 is due to intrinsic proofreading of the given C domain or rather an effect of altered C-A interactions.

As intended, plasmid based-complementation and production of WT FclJs (NRPS-9 to -11) in *X. szentirmaii ΔfclI ΔfclJ* led to the production of the expected range of shortened fabclavines (**15** - **17**) – with NRPS-9 synthesising peptides **15** - **17** and NRPS-10 and -11 only the proline derivative **15**. For the chimeric NRPSs 12 and 13, both carrying the putative promiscuous XU2 of FclJ (A6T6) from *X. szentirmaii* (Fig. 4b-c), only synthesis of **15** could be detected, and thus FclJ_A6’s substrate promiscuity could not be transferred – indicating that production of derivatives other than **15** is not the sole result of the A domain’s substrate specificity. This indication is further supported by NRPS-14 and -16, both carrying XU1 (C5A5T5C6) of FclJ from *X. szentirmaii* and XU2 from *X. budapestensis* (NRPS-14) and *X. hominickii* (NRPS- 16), respectively, now capable to biosynthesise peptides **15** - **17**. Interestingly, production of NRPS-15 only led to detectable amounts of **15**, while NRPS-17 led to the synthesis of **15** -**17**, but to a much lesser extent than NRPS-14 and -16 (Fig. 4b-c). Taken together, however, obtained *in situ* results confirm that, at least in case of fabclavine biosynthesis, the observed NRP diversification (*X. szentirmaii*) or specification (*X. hominickii, X. budapestensis*) is neither the result of the respective A domains promiscuity nor the C domains proofreading, but due to an extended gatekeeping function.

The extended gatekeeping function seems to manifest itself via specific interactions in the C-A interface – presumably by influencing the geometric degrees of freedom of the A domain. In the course of their catalytic cycle, A domains must adopt an open and closed conformation as well as the *C*-terminal subdomain has to undergo a ∼140° rotation (***Drake et al., 2016***). Altered C and A domain interactions might therefore influence these precisely coordinated transitions, editing selectivity and activity of respective A domains. Eventually, to reveal the very nature of the C domains’ influence on downstream A domains, next we aimed to in depth investigate a series of WT and chimeric C-A interfaces on a structural level.

### *In silico* approach maps hot-spot areas to determine crucial C-A didomain interactions

Since the structural elucidation of the targeted C-A interface forming proteins of NRPS-11 and NRPS-13 was intended but failed, we chose an *in silico* approach to characterise the extended gatekeeping function of C domains, at least in first approximation. We combined protein homology modelling (***Nayeem et al., 2006***) by using the Molecular Operating Environment (MOE 2019.01 (Molecular Operating Environment (MOE), 2021)) along with HSPred (***Lise et al., 2011***). The latter is a support-vector-machine-based method to predict critical interaction partners across protein-protein interfaces. HSPred systematically mutates *in silico* individual amino acids (excluding Pro and Gly) to alanine and calculates the changes in free energy of binding (*ΔΔG*). ‘Critical Interaction Partners’ or ‘Hot Spots Residues’ are defined as those residues for which *ΔΔG* ≥ 2 kcal/mol. The HSPred output score predicts mutated residues with a score greater than zero as hot spots (*ΔΔG* ≥ 2 kcal/mol) and negative scores (*ΔΔG* < 2 kcal/mol) as non-hot spots. Others are not involved in interface formation (Fig. 5; Fig. S4).

**Figure 5.**
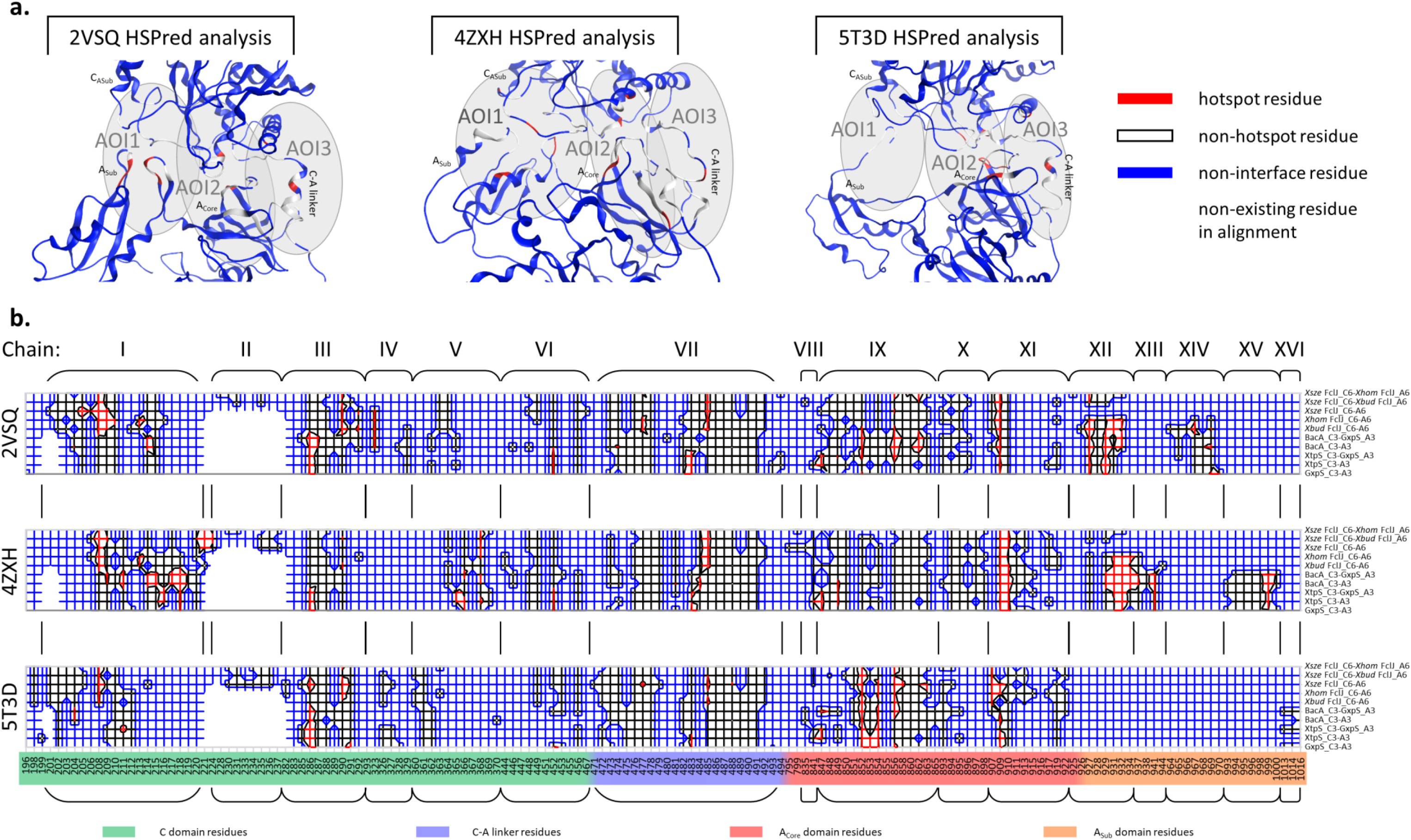
HSPred interface prediction for the created homology model. (**a**) Exemplary MOE models of the GxpS_C3-A3 homology model calculated with 2VSQ, 4ZXH, and the 5T3D as templates representative for all models from the HSPred analysis with highlighted hotspot residues (red), non-hotspot residues (white), non-interface residues (blue), and non-existing residues (colourless) in the reference alignment. The positions of the Area-Of-Interactions (AOI) 1-3 are grayed out in the structures. (**b**) Contour wireframe model only showing the interface forming residues of the HSPred interface prediction of the ten *Xsze* FclJ_C6-*Xhom* FclJ_A6, *Xsze* FclJ_C6-*Xbud* FclJ_A6, *Xsze* FclJ_C6-A6, *Xhom* FclJ_C6-A6, *Xbud* FclJ_C6-A6, BacA_C3-GxpS_A3, BacA_C3-A3, XtpS_C3-GxpS_A3, XtpS_C3-A3, and GxpS_C3-A3 models build in reference to the AB3403 (PDB ID: 4ZXH) (***Drake et al., 2016***), EntF (PDB ID: 5T3D) (***Drake et al., 2016***), and SrfA-C (PDB ID: 2SVQ) (***Tanovic et al., 2008***) templates. The interacting Chains I-XVI are indicated with black frames in the contour wireframe models. Colouring of the residue positions in the reference alignment domains (Tab. S11) is according to the C domain (green), C-A linker (blue), A_Core_ (red), and A_Sub_ (orange). **Supplementary Information – Table S8** **Supplementary Information – Table S10** **Supplementary Information – Table S11** **Supplementary Information – Figure S4** **Supplementary Information – Figure S5** **Supplementary Information – Figure S6** **Supplementary Information – Figure S7**

For comparative structural *in silico* analysis, we chose the *in vitro* assayed GxpS WT interface of P3 (GxpS_C3-A3) and hybrid interfaces of P4 (XtpS_C3-GxpS_A3) and P5 (BacA_C3-GxpS_A3) (cf. Fig. 2), respectively. In terms of sequence homology and catalytic activity, P4 and P5 were chosen to represent the two extremes, with P4 being almost WT-like and P5 completely unrelated. Additionally, we chose the *in situ* investigated FclJ_C6-A6 WT and hybrid interfaces presented above (Fig. 4b, NRPS-9 to -17). For homology modelling, we selected three different crystal structure templates of NRPS proteins with multiple domains: AB3403 (PDB ID: 4ZXH) (***Drake et al., 2016***), EntF (PDB ID: 5T3D) (***Drake et al., 2016***), and SrfA-C (PDB ID: 2SVQ) (***Tanovic et al., 2008***). This template diversity is intended to cover the majority of C-A interface- forming regions, such as the adenylate-forming conformation of AB3403, the thioester- forming conformation of EntF and the open conformation of SrfA-C (Fig. 5). Based on these templates, we created a total of 30 models of the selected C-A interfaces (Tab. S10), and then analysed the protein-protein interactions of the C and A domains via HSPred (Fig. S4).

The resulting interface plots in the contour wire model of the HSPred prediction (Fig. 5) show at a glance the distinct conformational changes of the different interfaces formed. In brief, numerous interactions can be ascribed to sixteen different Chains I- XVI that contribute to interface formation (Fig. 5b). These Chains, of which the C domain has six, the C-A linker one, and the A domain nine, interact in the so-called Area-Of-Interaction (AOI) 1-3 (Fig. 5a), which show highly dynamic conformational changes in the course of the catalytic NRP synthesis cycle. However, based on the chosen crystal structure templates, a comprehensive description of the structural C-A interface differences and the changes that occur during the transition of the individual catalytic states into each other can be found in the supplementary information (Explan. S1). In the following, the most important differences between the WT and hybrid C-A interfaces modelled and analysed in this work are highlighted. It should be noted that the amino acid numbering used below is based on the residue position in the protein sequence alignment of calculated models (Tab. S11).

When the WT C-A interface of P3 is compared to the hybrid interfaces of both, the C- A interface of P4 and P5, differences mainly are present in the AOI1 A_Sub_ area of all catalytic states (Fig. 5). P4 introduces additional hot spot residues in the adenylate forming conformation (Fig. 5, P4_4ZXH_) via Chain I (R216) & V (R365), loosened A_Core_/A_Sub_ transition of Chain III (D291) and a tightening to the C-A linker in AOI2 of Chain IX (R841, R847, Y849) in the thioester forming conformation (Fig. 5, P4_5T3D_). Although the exchange of GxpS_C3 for XtpS_C3 in P4 leads to a slightly closer interaction of the C domain with the A_Sub_ domain during adenylate and thioester formation as well as to a slight relaxation of the A_Core_/A_Sub_ hinge region (AA907 to AA944, Tab. S11), the hybrid interface of P4 appears to be very similar to the wild type one of P3 – as could be expected from their high sequence similarity. It is therefore not surprising that the introduced changes at the interface have almost no effect (cf. Fig. 2) on the catalytic activity and substrate activation profile of GxpS_A3, as evidenced in the *in vitro* and *in vivo* assays. In turn, when the interface of P5 is compared to the WT P3 interface, multiple additional hot residues within Chain I (R204, K209, D211, Y214, D215, K217, R218), and Chain V (R323) in all three models (Fig. 5, P5_2VSQ_, P5_4ZXH_, and P4_5T3D_) could be observed, indicating a much stronger association around the otherwise flexible A_Sub_ domain in AOI1. This increased rigidity of the C-A interface seems to interfere with the ability of the A domains to switch between the different conformations required for proper substrate binding/release and adenylate formation (***Reimer et al., 2018***), as highlighted by P5’s poor catalytic *in vitro* activity (cf. Fig. 2c).

Compared to all other interfaces, investigated within present work, the *Xsze* FclJ_C6- A6 WT interface (NRPS-9) as well as the recombinant interfaces from NRPS-12 to -17 show a novel interaction site with a significant impact on especially the substrate binding/release conformation (Fig. 5, models based on 2VSQ). All these constructs have the *Xsze* FclJ_C6 domain in common – introducing the unique Chain II (Fig. S5-7) that highly contributes to the C-A interface formation by tightly interacting with the respective A_Sub_ domains in AOI1. Chain II shows up in all conformations but with the highest abundance of hot residues in the adenylate forming state (models based on 4ZXH) at S220, H221 and E224. Chain XIV, which regularly participates in interface formation in substrate binding/release (Fig. 5, models based on 2VSQ models), disappears entirely in the constructs containing the respective *Xsze* C domain. Further, all of the studied fabclavine interfaces lack contribution of Chain XV and XVI, which have been involved in the adenylate-forming and thioester-forming conformation in the GxpS, XtpS and BacA interfaces (Fig. 5b). Consequently, the pronounced conformational changes of the A_Sub_ domain observed *in silico* are less determined by the opposite C-domain, suggesting dynamic detachment. Therefore, the previously described promiscuity at this position in fabclavine biosynthesis (***Wenski et al., 2020***) does not seem to be the exclusive result of the FclJ_A6 domains activity, but from the extended gatekeeping function of the FclJ_C6 domain that grants additional spatial flexibility mainly in AOI1.

In sum, the *Xsze* FclJ_C6 seems to follow its very own path in C-A interface formation with considerable differences, especially in the yet unreported interaction of the C domain’s Chain II, and loss of interaction of Chain XIV, XV & XVI with A_Sub_, extending its already dynamic 30° rotation from the substrate binding/release to the adenylate forming conformation and subsequent 140° body torsion in the thioester forming conformation (***Drake et al., 2016***). Interestingly, this extended gatekeeping, leading to less spatial restrictions on the A domain’s movement, is not only transferred on a structural level when chimeric interfaces are created, but also influences the substrate recognition capacity of the respective downstream A domain.

## Discussion

Although biochemical *in vitro* characterisations of individual domains or modules greatly contributed to our current advanced understanding of all fundamental catalytic reactions in NRP synthesis, obtained results are difficult to extrapolate to full length multi-domain and -modular mega-synthetases – as evidenced from long standing design paradigms currently under debate, such as the inseparability of C-A didomains and the C domains gatekeeping role (***Baltz, 2014***; ***Baunach et al., 2021***; ***Belshaw et al., 1999***; ***Bozhüyük et al., 2018***; ***Bozhüyük et al., 2019a***; ***Calcott et al., 2020***; ***Lautru and Challis, 2004***; ***Süssmuth and Mainz, 2017***). Especially the latter has most recently been revised by landmark contributions investigating the evolution of NRPSs, i.e. by drawing a landscape of evolutionary recombination events (***Baunach et al., 2021***), and exploring the C domains acceptor site specificity via gene shuffling experiments (***Calcott et al., 2020***). These contributions have led to the currently prevailing view that C domains, or rather the proofreading role attributed to them, can be neglected in the creation of hybrid NRPSs and unnecessarily has complicated NRPS engineering campaigns (***Alanjary et al., 2019***; ***Brown et al., 2018***). Although this view, to some extent, contradicts most recent structural insights (***Izoré et al., 2021***) as well as our own observations made when developing reproducible engineering strategies (***Bozhüyük et al., 2018***; ***Bozhüyük et al., 2019a***), we do not doubt that the core findings of these studies are accurate. Nevertheless, when reviewing our data of previous works (***Bozhüyük et al., 2018***; ***Bozhüyük et al., 2019a***; ***Bozhüyük et al., 2021***), we could notice that there must be more to it and that the current black and white view of this issue misses the complexity of this problem.

Therefore, with this work, we have attempted to shed light on the recently sparked debate about the role of C-domains in the non-ribosomal synthesis of peptides (***Dekimpe and Masschelein, 2021***). Based on our established expertise in engineering NRPSs, we have tried to rethink the problem and approach it from different angles, focusing particularly on the changing behaviour of A domains in the context of chimeric biosynthetic pathways. We have devised a comprehensive experimental procedure ranging from *in vitro* (Fig. 2) and *in vivo* (Fig. 3) characterisations targeting our preferred model system GxpS (Fig. 1) to *in situ* investigations of the fabclavine producing BGC (Fig. 4) and *in silico* characterisation of selected C-A didomain interfaces created in this study (Fig. 5).

Within this study, it has been our experience that *in vitro* results can differ significantly depending on the assay chosen and can paint a picture that contradict the results obtained *in vivo*. However, all *in vitro* and *in vivo* results concerning the selected promiscuous GxpS_A3 framework, revealed a significant influence of all C domains on both (Fig. 2-3), the general catalytic activity and the substrate recognition profile within the identified “substrate group specificity” of the GxpS_A3 domain. Interestingly, in terms of phylogenetic distance and sequence homology, less similar C domains (e.g. AmbS_indi.__C5 & BicA_C3) seem to have a more pronounced effect (Fig. 2 & 3) – in both directions (Fig. 2b, NRPS-6 & -8). This effect could be described as an extended gatekeeping function of the C-A interface on fine-tuning the A domains selectivity and thus contributing to its role as a primary substrate selectivity filter – as also reported from previous *in vitro* characterisations of A domains of the microcystin-producing NRPS (***Meyer et al., 2016***).

Noteworthy, BacA_C3 as a representative outside the *Photo-* and *Xenorhabdus* genus exerts significant influence on the interface-dynamics which almost abolished the activity *in vitro* (Fig. 2c). This observation now explains our previous inability to recombine building blocks of Gram-negative and -positive origin with each other by using the XU strategy in most cases (***Bozhüyük et al., 2018***). Interestingly, most recently we were able to functionally apply the very same interface (P5) by introducing synthetic leucin zippers (type S NRPSs) within the C-A linker region (***Bozhüyük et al., 2021***). The created type S NRPS not only synthesized a thiazoline containing peptide with high fidelity at high titre, but the GxpS_A3 domain exclusively activated leucine, thus completely omitting the domains *in vitro* confirmed preferred substrate phenylalanine (***Bozhüyük et al., 2021***) – representing another illuminating example of how altered C-A interactions are capable to contribute to the A domains attributed role as primary selectivity filter.

Eventually, by targeting the fabclavine BGCs from *X. szentirmaii*, *X. budapestensis*, and *X. hominicii* XU substitutions to alter C-A interface interactions could be made that were least out of their natural context (Fig. 4). Here, a more dominant and transferable extended gatekeeping effect of the *Xsze* FclJ_C6 domain could be observed, which, through an additional loop in the interface (Fig. S5-7), mainly interacts with the A_Sub_ domain. Introduction of *Xsze* XU1 (C5A5T5C6) upstream of FclJ_A6 from *X. budapestensis.* and *X. hominickii.* empowered the formerly proline specific A domains to also activate valine and threonine (Fig. 4b-c).

In the final analysis, along with most recently published findings, present data suggests that a general strong gatekeeping function of C domains can be excluded and might rather be the exception. Yet, we were able to highlight that C domains do have a great effect on selectivity of adjacent A domains via an extended gatekeeping function and should definitely be considered when artificial NRPSs are created. Our *in silico* analysis revealed that this extended gatekeeping function manifests itself within the respective formed C-A interfaces during all catalytic stages (Fig. 5).

At this point, we must revise the established XU concept assembly rules to guide the debate about the specificity-imparting properties of C domains. Accordingly, the second XU rule (’The specificity of the downstream C domain must be respected’ (***Bozhüyük et al., 2018***)) should not have been focusing on the attributed C domain’s acceptor site specificity, but on the very nature of interfaces that C domains can form with an A domain of interest. Nevertheless, the second rule could still serve as a rule of thumb to directly guide engineering attempts without prior in-depth analysis, as it is more likely that C domains upstream of A domains with the same or similar specificity can form a more similar and thus functional interface (Fig. S8). With our current knowledge, C domains that do not directly conform to the C domain dogma no longer need to be excluded. Therefore, the C-A interface is assumed to have a more significant contribution to a selectivity filter function, in turn, highlighting the great advantage of the XUC concept which preserves these interfaces (***Bozhüyük et al., 2019a***).

We hope that the present work can make a constructive contribution to the ongoing debate and is just one more viewpoint of many to follow. We look forward to the forthcoming results and intend to contribute further insights soon, yet again from a different angle. Therefore, we would like to conclude with the words of a great scientist: ‘When you change the way you look at things, the things you look at change.’ – Max Planck

## Material and methods

### Cultivation of strains

All *E. coli* and *Xenorhabdus* strains were cultivated in liquid or on solid LB-medium (pH 7.5, 10 g/L tryptone, 5 g/L yeast extract and 5 g/L NaCl). Solid media contained 1 % (w/v) agar. Kanamycin (50 μg/ml) and chloramphenicol (34 μg/ml) were used as selection markers. All *E. coli* cultures were cultivated at 37 °C, 22 °C, or 16 °C for peptide or protein production purposes. *Xenorhabdus* strains were grown at 30 °C.

### Cloning of biosynthetic gene clusters

Genomic DNA of selected *Xenorhabdus* and *Photorhabdus* strains were isolated using the Qiagen Gentra Puregene Yeast/Bact Kit. All PCRs were performed with oligonucleotides obtained from Eurofins Genomics (Tab. S4). NRPS fragments for HiFi cloning (NEB) were amplified with primers coding for the respective homology arms (20-30 bp) in a two-step PCR program. The coding sequences for the His6-Tag were amplified with the pCOLADUET^TM^-1 (Merck/Millipore) plasmid backbone. Polymerases Phusion High-Fidelity DNA polymerase (Thermo Scientific), Q5 High-Fidelity DNA polymerase (New England BioLabs), and Velocity DNA polymerase (Bioline) were used according to the manufacturers’ instructions. DNA purification was performed using Invisorb Fragment CleanUp or MSB Spin PCRapace Kits (stratec molecular). All generated plasmids (Tab. S3) were introduced into *E. coli* DH10B::*mtaA* (***Schimming et al., 2014***) by either electroporation. Each NRPS (subunit) was under the control of a *P_BAD_* promotor for peptide production or a *tac*I promotor for protein expression. Plasmid isolation from *E. coli* was achieved with the Invisorb Spin Plasmid Mini Two Kit (stratec molecular).

### Generation of deletion mutants

The generation of deletion mutants was performed as described previously (***Brachmann et al., 2007***; ***Wenski et al., 2019***): The upstream and downstream flanking regions of the corresponding gene (approximately 1000 bp) were amplified and cloned into the either PCR-amplified or digested vector pEB17 to generate deletion vectors (***Bode et al., 2019***). After the Hot Fusion Assembly *E. coli* S17 were transformed with the vectors, followed by conjugation with the corresponding *Xenorhabdus* strain as described previously (***Fu et al., 2014***; ***Philippe et al., 2004***; ***Simon et al., 1983***; ***Thoma and Schobert, 2009***).

### Transformation of *X. szentirmaii*

Hetero- and homologous complementation as well as NRPS-engineering plasmids were transformed into the corresponding *X. szentirmaii* strain by heatshock transformation by an adapted protocol of Xu et al. as described previously (***Wenski et al., 2019***; ***Xu et al., 1989***).

### Heterologous expression of NRPS templates and LC-MS analysis

Constructed plasmids were transformed into *E. coli* DH10B::*mtaA* (***Schimming et al., 2014***). Cells were grown overnight in LB medium containing the necessary antibiotics (50 μg/ml kanamycin, or 34 μg/ml chloramphenicol). 100 μl of an overnight culture were used for inoculation of 10 ml LB-cultures supplemented with the respective antibiotics as selection markers and additionally containing 0.002 mg/ml L-arabinose and 2 % (v/v) XAD-16. After incubation for 72 h at 22 °C the XAD-16 was harvested. One culture volume methanol was added and incubated for 30 min at RT. The organic phase was filtrated, and a sample was taken of the cleared extract. After centrifugation (17,000 x *g*, 20 min) the methanol extracts were used for LC-MS analysis. All measurements were performed by using an Ultimate 3000 LC system (Dionex) with an ACQUITY UPLC BEH C18 column (130 Å, 2.1 x 50 mm, 1.7 μm particle size; Waters) at a flow rate of 0.4 ml min^-1^ using acetonitrile (ACN) and water containing 0.1 % formic acid (v/v) in a gradient ranging from 5-95 % of ACN over 16 min (40 °C) coupled to an AmaZonX (Bruker) electron spray ionization mass spectrometer. High-resolution mass spectra were obtained on an Ultimate 3000 RSLC (Dionex) coupled to an Impact II qTOF (Bruker) equipped with an ESI Source set to positive ionization mode. The software DataAnalysis 4.3 (Bruker) was used to evaluate the measurements.

### Expression and purification of His6-tagged proteins

For overproduction and purification of the His6-tagged ∼72 kDa GxpS A3-T3, ∼98 kDA GxpS C3_ASub_-A3-T3, ∼122 kDa GxpS C3-A3-T3 and ∼122 kDa XtpS C3 GxpS A3-T3 a 5 mL overnight culture in LB medium of *E. coli* BL21 (DE3) Gold cells harboring the corresponding expression plasmid and the TaKaRa chaperone-plasmid pTF16 (TAKARA BIO INC.) were used to inoculate 500 mL of autoinduction medium (464 mL LB medium, 500 μL 1M MgSO4, 10 mL 50×5052, 25 mL 20xNPS) containing 50 μg/mL kanamycin and 34 μg/mL chloramphenicol. The cells were grown at 37 °C up to an OD_600_ of 0.6. Following the cultures were cultivated for additional 72 h at 16 °C. The cells were pelleted (10 min, 4,000 rpm, 4 °C) and stored overnight at -20 °C.

For protein purification the cells were resuspended in binding buffer (500 mM NaCl, 20 mM imidazol, 50 mM HEPES, 10 % (w/v) glycerol, pH 8.0). For cell lysis benzonase (500 U, Fermentas), protease inhibitor (Complete EDTA-free, Roche), 0.1 % Triton-X and lysozym (0.5 mg/mL, ∼20,000 U/mg, Roth) were added and the cells were incubated rotating for 30 min at 4 °C. After this the cells were placed on ice and lysed by ultra-sonication. Subsequently, the lysed cells were centrifuged (25,000 rpm, 30 min, 4°C).

The yielded supernatant was passed through a 0.2 μm filter and loaded with a flow rate of 0.5 mL/min on a 5 mL HisTrap HP column (GE Healthcare) equilibrated with 12 CV binding buffer. Unbound protein was washed off with 8 CV with 4 % and 8 CV with 8 % elution buffer (500 mM NaCl, 500 mM imidazol, 50 mM HEPES, 10 % (w/v) glycerol, pH 8.0). The purified protein of interest was eluted with 35 % elution buffer. Following, the purified protein containing fraction was concentrated (Centriprep^®^ Centrifugal Filters Ultacel^®^ YM - 50, Merck Millipore).

### MALDI-Orbitrap-MS

Samples were prepared for MALDI-analysis as a 1:1 dilution in 9-aminoacridine in acetone (10 mg/mL in 99 % aceton), spotted onto a polished stainless-steel target, and air-dried. MALDI-Orbitrap-MS analyses were performed with a MALDI LTQ Orbitrap XL (Thermo Fisher Scientific,Inc., Waltham, MA) equipped with a nitrogen laser at 337 nm. The following instrument parameters were used: laser energy, 27 μJ; automatic gain control, on; auto spectrum filter, off; resolution, 30,000; plate motion, survey CPS. Mass spectra were obtained in negative ion mode over a range of 500 to 540 m/z. The mass spectra for γ-[^16^O_4_]-ATP exchange analysis were acquired by averaging 50 consecutive laser shots. Spectral analysis was conducted using XCalibur Qual Browser (version 2.0.7; Thermo Fisher Scientific, Inc., Waltham, MA).

### γ-[^18^O_4_]-ATP-Pyrophosphat Exchange Assay

The γ-[^18^O_4_]-ATP-Pyrophosphat Exchange Assay was performed as published previously (***Phelan et al., 2009***) with the following changes described below.

The 2 µl amino acid solution (3 mM amino acid, 15 mM PP_i_/Tris), 2 μL γ-[^18^O_4_]-ATP (3 mM γ-[^18^O_4_]-ATP, 15 mM MgCl_2_/Tris) and 2 μL of purified protein (c = 2 mg/ml) were incubated for 2 h at RT. The reactions were stopped by freezing the sample at -20 °C and addition of 6 μL 9-aminoacridine in acetone (10 mg/mL) for MALDI-Orbitrap-MS analysis.

### Multiplexed hydroxamate assay (HAMA)

The hydroxamate formation assay was performed as published previously (***Stanišić et al., 2019***). The 100 μL reaction volume containing 50 mM TRIS (pH 7.6), 5 mM MgCl_2_, 150 mM hydroxylamine (pH 7.5 - 8, adjusted with HCl), 5 mM ATP, 1 mM TCEP and 2 µM of purified enzyme were stared by adding 1 mM proteinogenic amino acid mix (in 100 mM TRIS, pH 8) and incubated for 30 min at RT. The reactions were stopped by diluting in 10-fold 95 % acetonitrile (ACN) and water containing 0.1 % formic acid. All measurements were performed by using an Ultimate 3000 RSLC (Dionex) with an ACQUITY UPLC BEH Amide Column, 130 Å, 1.7 µm, 2.1 mm X 50 mm, 1/pkg coupled to an Impact II qTOF (Bruker) equipped with an ESI Source set to positive ionization mode. UPLC conditions were performed as published previously (***Stanišić et al., 2019***). The software DataAnalysis 4.3 (Bruker) was used to evaluate the measurements.

### Homology modelling and interface identification

The homology-modelling was performed with the homology modeling algorithm within MOE (Molecular Operating Environment). This process undergoes an (I) initial partial geometry, where all coordinates are copied if residue identity is conserved. Next, a (II) Boltzmann-weighted randomized sampling, which (IIa) consists of a backbone fragments collection from a high-resolution structural database, and alternative side chain conformations assembly from an extensive rotamer library for non-identical residues and (IIb) a creation of independent number models based upon loop and side chain placements scored by a contact energy function (***Nayeem et al., 2006***).

For homology modelling, the C-A didomains within the crystal structure of AB3403 (PDB-ID: 4ZXH), EntF (PDB-ID: 5T3D) and SrfA-C (PDB-ID: 2SVQ) were selected as homologous-protein-templates.

With the homology models build, HSPred (***Lise et al., 2011***), a support vector machine(SVM)-based method, was used to predict the critical interaction partners across the interface. This approach systematically mutated individual amino acids s (excluding Pro and Gly) to alanine and calculates the changes in free energy of binding (*ΔΔG*). ‘Critical Interaction Partners’ or ‘Hot Spots Residues’ are defined as those residues for which *ΔΔG* ≥ 2 kcal/mol. The HSPred output score (its exact calculation can be read here (***Lise et al., 2011***)) predicts mutated residues with a score greater than zero as hot spots (*ΔΔG* ≥ 2 kcal/mol) and negative scores (*ΔΔG* < 2 kcal/mol) as non-hot spots. Others are not involved in the interface.

### Peptide quantification

The absolute production titers of selected peptides were calculated with calibration curves based on pure synthetic **1**, (for quantification of **1**–**4**), **5** (for quantification of **5**, **9**-**10**, **12**-**14**), **6** (for quantification of **6**), **7** (for quantification of **7**), **8** (for quantification of **8**), **11** (for quantification of **11**), **13** (for quantification of **13**). Therefore, the pure compounds were prepared at different concentrations (100, 50, 25, 12.5, 6.25, 3.125, 1.56, 0.78, 0.39, and 0.195 μg/ml) and measured by LC-MS using HPLC/MS measurements as described above. The peak area for each compound at different concentrations was calculated using Compass Data Analysis and used for the calculation of a standard curve passing through the origin. Triplicates of all *in vivo* experiments were measured. The pure peptide standards **1**, **5**, **6**, **7**, **8**, **11**, and **13** were synthesized in-house (***Bozhüyük et al., 2018***).

### Chemical synthesis

Chemical synthesis of all peptides was performed as described previously (***Bozhüyük et al., 2018***).

## Acknowledgements

This work was supported by the LOEWE research cluster MegaSyn, the LOEWE center TBG, both funded by the state of Hesse and an ERC advanced grant (grant agreement number 835108).

## Competing interests

The authors declare no competing interests.

## Notes

### Competing Interest Statement

The authors have declared no competing interest.

