## Supplementary Information for "Influence of condensation domains on activity and specificity of adenylation domains"

#### Supplementary Tables

**Table S1.** ESI-MS data of all produced compounds. The compound amino acid compositions marked with an asterisk are attached with one malonate unit and two polyamine units corresponding to fabclavines biosynthesis (***Wenski et al., 2019***) to obtain the denoted molecular mass.

| Compound | amino acid composition | theoretical mass-to-charge ratio (*m/z*) [M+H]$^+$ | Molecular formular | Reference |
|---|---|---|---|---|
| **1** | cyclo(vLflL) | 586.3962 | $C_{32}H_{51}N_5O_5$ | (***Bozhüyük et al., 2018***) |
| **2** | cyclo(ILflL) | 600.4119 | $C_{33}H_{53}N_5O_5$ | (***Bozhüyük et al., 2018***) |
| **3** | cyclo(vLIlL) | 552.4119 | $C_{29}H_{53}N_5O_5$ | (***Bozhüyük et al., 2018***) |
| **4** | cyclo(ILIlL) | 566.4275 | $C_{30}H_{55}N_5O_5$ | (***Bozhüyük et al., 2018***) |
| **5** | FIL | 392.2544 | $C_{21}H_{33}N_3O_4$ | - |
| **6** | fIL | 392.2544 | $C_{21}H_{33}N_3O_4$ | - |
| **7** | IIL | 358.2700 | $C_{21}H_{35}N_3O_4$ | - |
| **8** | LIL | 358.2700 | $C_{21}H_{35}N_3O_4$ | - |
| **9** | iIL | 358.2700 | $C_{21}H_{35}N_3O_4$ | - |
| **10** | lIL | 358.2700 | $C_{21}H_{35}N_3O_4$ | - |
| **11** | YIL | 408.2493 | $C_{21}H_{33}N_3O_5$ | - |
| **12** | WIL | 431.5405 | $C_{23}H_{34}N_4O_4$ | - |
| **13** | MIL | 376.2265 | $C_{17}H_{33}N_3O_4S$ | - |
| **14** | VIL | 344.2544 | $C_{17}H_{33}N_3O_4$ | - |
| **15** | TP* | 713.6263 | $C_{39}H_{80}N_6O_5$ | (***Wenski et al., 2020***) |
| **16** | TV* | 715.6420 | $C_{39}H_{82}N_6O_5$ | (***Wenski et al., 2020***) |
| **17** | TT* | 717.62.12 | $C_{38}H_{80}N_6O_6$ | (***Wenski et al., 2020***) |

**Table S2.** Strains used in this work.

| Strain | Genotype | Reference |
|---|---|---|
| *E. coli* DH10B | $F^-$ *mcr*A Δ(*mrr*-*hsd*RMS-*mcr*BC) φ80*lac*ZΔM15 Δ*lac*X74 *rec*A1 *end*A1 *ara*D139 Δ(*ara*-*leu*)7697 *gal*U *gal*K $\lambda^-$ *rps*L($Str^R$) *nup*G/ - | Invitrogen |
| *E. coli* DH10B::*mtaA* | *E. coli* DH10B with Δ*entD*::*mtaA* | (***Schimming et al., 2014***) |
| *E. coli* BL21 (DE3) Gold | *E. coli* B $F^-$ *ompT hsdS*($r_B^-$ $m_B^-$) *dcm*$^+$ $Tet^r$ *gal* λ(DE3) *endA* Hte | Invitrogen |
| *E. coli* S17-1 λpir | $Tp^r$ $Sm^r$ *rec*A, *thi*, *pro*, *hsd*R-M+RP4: 2- Tc:Mu:Km Tn7 λpir | Invitrogen |
| *P. luminescens* TTO1 | wildtype | NC_005126.1_NCBI |
| *X. nematophila* ATCC 19061 | wildtype | FN667742_NCBI |
| *B. licheniformis* ATCC 10716 | wildtype | M. A. Marahiel / ATCC |
| *X. szentirmaii* DSM 16338 | wildtype | NIBV00000000_NCBI |
| *X. budapestensis* DSM 16342 | wildtype | NIBS00000000_NCBI |
| *X. hominickii* DSM 17903 | wildtype | NJAI00000000_NCBI |
| *X. szentirmaii* Δ*fclIJ* | wildtype *ΔfclI*, *ΔfclJ* | this work |

**Table S3.** Plasmids used in this work.

| Plasmid | Genotype | Reference |
|---|---|---|
| pTF16 | Chaperone tig, L-arabinose inducible Promotor *araB*, $cm^R$ | TaKaRa Bio Inc., Singapore |
| pCOLA_GxpS A3-T3 | ori ColA, $kan^R$, *tac*I, HIS-*gxpS*_A3T3 | this work |
| pAD_GxpS $C_A$3-A3-T3 | ori ColA, $kan^R$, *tac*I, *gxpS*_CA3A3T3 | this work |
| pAD_GxpS C3-A3-T3 | ori ColA, $kan^R$, *tac*I, HIS-*gxpS*_C3A3T3 | this work |
| pAD_XtpS C3 GxpS A3-T3 | ori ColA, $kan^R$, *tac*I, HIS-*xtpS*_C3 *gxpS*_A3T3 | this work |
| pEB17 | R6Kγ ori, oriT, *araC*, *ara*BAD promoter, $Km^r$ | (***Bode et al., 2015***) |
| pEB17_Δ*fclIJ* *X. szentirmaii* | pEB17 with *Xsze fclI, Xsze fclJ* | this work |
| pCOLA_ara_tacI_*fclJ Xsze* | ori ColA, $kan^R$, *araC-$P_{BAD}$ Xsze fclj*_C5A5T6C6A6T6 and *tac*I | this work |
| pCOLA_ara_tacI_*fclJ Xbud* | ori ColA, $kan^R$, *araC-$P_{BAD}$ Xbud fclj*_C5A5T6C6A6T6 and *tac*I | this work |
| pCOLA_ara_tacI_*fclJ Xhom* | ori ColA, $kan^R$, *araC-$P_{BAD}$ Xhom fclj*_C5A5T6C6A6T6 and *tac*I | this work |
| pDD3 | ori ColA, $kan^R$, *araC-$P_{BAD}$ Xbud fclj*_C5A5T6C6 *Xsze. fclj*_A6T6 and *tac*I | this work |
| pDD4 | ori ColA, $kan^R$, *araC-$P_{BAD}$ Xhom fclj*_C5A5T6C6 *Xsze fclj*_A6T6 and *tac*I | this work |
| pDD5 | ori ColA, $kan^R$, *araC-$P_{BAD}$ Xsze fclj*_C5A5T6C6 *Xbud fclj*_A6T6 and *tac*I | this work |
| pDD6 | ori ColA, $kan^R$, *araC-$P_{BAD}$ Xhom fclj*_C5A5T6C6 *Xbud fclj*_A6T6 and *tac*I | this work |
| pDD7 | ori ColA, $kan^R$, *araC-$P_{BAD}$ Xsze fclj*_C5A5T6C6 *Xhom fclj*_A6T6 and *tac*I | this work |
| pDD8 | ori ColA, $kan^R$, *araC-$P_{BAD}$ Xbud fclj*_C5A5T6C6 *Xhom fclj*_A6T6 and *tac*I | this work |

**Table S4.** Oligonucleotides used in this work. Oligonucleotides marked with an asterisk were used in combination with pEB74_rv (5'-GGAATTCCTCCTGTTAGCCCAA-3').

| Plasmid | Oligonucleotide | Sequence (5'->3') | Template |
|---|---|---|---|
| pCOLA_GxpS A3-T3 | pCOLADuet_Gibson Primer A3 Insert FW | CATCACCATCATCACCACCCTCAACAACCTGTCACGGC | *P. laumondii* TTO1 |
| | pCOLADuet_Gibson Primer A3 Insert RV | CAGCCTAGGTTAATTAAGCTGTTATTTTTCGAACTGCGGGTGGCTCCAAGT CAGATCAATCAGCGGCAAC | *P. laumondii* TTO1 |
| | DUET_Gib_FW | CAGCTTAATTAACCTAGGCTG | pCOLADuet_ara/tacI |
| | DUET_Gib_RV | GTGGTGATGATGGTGATG | pCOLADuet_ara/tacI |
| pAD_GxpS $C_A$3-A3-T3 | AD21 | TCGAGTCTGGTAAAGAAACC | *P. laumondii* TTO1 |
| | AD22 | CTGGCTGTGGTGATGAT | *P. laumondii* TTO1 |
| | AD12 | CATCATCACCACAGCCAGTCAGGTGAAGGAGTACAGGC | *P. laumondii* TTO1 |
| | AD14 | CTTTACCAGACTCGATTACGGTACTTGCTCAACAATACG | *P. laumondii* TTO1 |
| | DUET_Gib_FW | CAGCTTAATTAACCTAGGCTG | pCOLADuet_ara/tacI |
| | DUET_Gib_RV | GTGGTGATGATGGTGATG | pCOLADuet_ara/tacI |
| pAD_GxpS C3-A3-T3 | AD21 | TCGAGTCTGGTAAAGAAACC | *P. laumondii* TTO1 |
| | AD22 | CTGGCTGTGGTGATGAT | *P. laumondii* TTO1 |
| | AD13 | CCATCATCACCACAGCCAGATCTGTGCACAACGTAATACG | *P. laumondii* TTO1 |
| | AD14 | CTTTACCAGACTCGATTACGGTACTTGCTCAACAATACG | *P. laumondii* TTO1 |
| | DUET_Gib_FW | CAGCTTAATTAACCTAGGCTG | pCOLADuet_ara/tacI |
| | DUET_Gib_RV | GTGGTGATGATGGTGATG | pCOLADuet_ara/tacI |
| | AD10 | ATCATCACCACAGCCAGCGTCATGCGAACAGTGATC | *X. nematophila* ATCC 19061 |

| | | | |
|---|---|---|---|
| pAD_XtpS C3 GxpS A3-T3 | AD74 | GGTTCTTCGGTCGCATTCCAGGTTTTTAACAACAATGTGC | *X. nematophila* ATCC 19061 |
| | AD75 | GCACATTGTTGTTAAAAACCTGGAATGCGACCGAAG | *P. laumondii* TTO1 |
| | AD14 | CTTTACCAGACTCGATTACGGTACTTGCTCAACAATACG | *P. laumondii* TTO1 |
| | DUET_Gib_FW | CAGCTTAATTAACCTAGGCTG | pCOLADuet_ara/tacI |
| | DUET_Gib_RV | GTGGTGATGATGGTGATG | pCOLADuet_ara/tacI |
| pEB17_Δ*fclIJ* *X. szentirmaii* | SW112_I_LF_fw | CGATCCTCTAGAGTCGACCTGCAGCTATTTCAATTACTTGATTGAATGCGG | pEB17 |
| | SW485_XszIJ_rv | AGCTCTGTTCCCTTTTCCCATAGCCATCATTCCTTCAATAAGAATTTATTACC | *X. szentirmaii* DSM 16338 |
| | SW486_XszIJ_fw | TATTGAAGGAATGATGGCTATGGGAAAAGGGAACAGAGC | *X. szentirmaii* DSM 16338 |
| | SW390_XszJ_RF_rv | GAGAGCTCAGATCTACGCGTTTCATATGGGTCGCTGTACCGTGTGC | pEB17 |
| pCOLA_ara_tacI_ fclJ *Xsze.* | SW440_XsJ_KoL_fw | CCATACCCGTTTTTTTGGGCTAACAGGAGGAATTCCATGCAGCAAGATACA TTTAAATTTAAAGCATCCC | *X. szentirmaii* DSM 16338 |
| | W441_XsJ_KoL_rv | CGAGCCGATGATTAATTGTCAACAGCTCCTGCAGTTACCCTTTGTGCTGCA TTTTTTTCTGC | *X. szentirmaii* DSM 16338 |
| pCOLA_ara_tacI_ fclJ *Xbud* | SW478_XbudJ_fw | CCATACCCGTTTTTTTGGGCTAACAGGAGGAATTCCATGCAGGAAAATACAT TTCAATTTAAGGC | *X. budapestensis* DSM 16342 |
| | SW479_XbudJ_rv | CGAGCCGATGATTAATTGTCAACAGCTCCTGCAGCTATTCCTTTGCCTGTTT CTGCTGC | *X. budapestensis* DSM 16342 |
| pCOLA_ara_tacI_ fclJ *Xhom* | SW480_XhomJ_fw | CCATACCCGTTTTTTTGGGCTAACAGGAGGAATTCCATGCAGGAAAACACA TATAAATTCAAAGC | *X. hominickii* DSM 17903 |
| | SW481_Xhom_rv | CGAGCCGATGATTAATTGTCAACAGCTCCTGCAGTCATTGTTGCTGCATTTT TTTCTGC | *X. hominickii* DSM 17903 |
| pDD3 | SW491 | CCATACCCGTTTTTTTGGGCTAACAGGAGGAATTCCATGCAGGAAAATACAT TTCAATTTAAGGC | *X. budapestensis* DSM 16342 |

| | | | |
|---|---|---|---|
| | SW492 | CCGGATGCCCCCTGACTGAAACTTTCAATCTTCCGTTGTTGGATTATCG | *X. budapestensis* DSM 16342 |
| | SW493* | AACAACGGAAGATTGAAAGTTTCAGTCAGGGGGCATCCGG | *X. szentirmaii* DSM 16338 |
| pDD4 | SW494 | CCATACCCGTTTTTTTGGGCTAACAGGAGGAATTCCATGCAGGAAAACACA TATAAATTCAAAGC | *X. hominickii* DSM 17903 |
| | SW495 | CCGGATGCCCCCTGACTGAAACTCTCAATCAGCCGTTGTTGTG | *X. hominickii* DSM 17903 |
| | SW496* | AACAACGGCTGATTGAGAGTTTCAGTCAGGGGGCATCC | *X. szentirmaii* DSM 16338 |
| pDD5 | SW497 | CCATACCCGTTTTTTTGGGCTAACAGGAGGAATTCCATGCAGCAAGATACA TTTAAATTTAAAGCATCCC | *X. szentirmaii* DSM 16338 |
| | SW498 | CCCGGCGCTCCCTGATTGAAGGCTTCAATTTTCCGCTGCTGGG | *X. szentirmaii* DSM 16338 |
| | SW499* | AGCAGCGGAAAATTGAAGCCTTCAATCAGGGAGCGCCG | *X. budapestensis* DSM 16342 |
| pDD6 | SW500 | CCATACCCGTTTTTTTGGGCTAACAGGAGGAATTCCATGCAGGAAAACACA TATAAATTCAAAGC | *X. hominickii* DSM 17903 |
| | SW501 | CCCGGCGCTCCCTGATTGAAACTCTCAATCAGCCGTTGTTGTG | *X. hominickii* DSM 17903 |
| | SW502/514 | AACAACGGCTGATTGAGAGTTTCAATCAGGGAGCGCCGG | *X. budapestensis* DSM 16342 |
| pDD7 | SW503 | CCATACCCGTTTTTTTGGGCTAACAGGAGGAATTCCATGCAGCAAGATACA TTTAAATTTAAAGCATCCC | *X. szentirmaii* DSM 16338 |
| | SW504 | CCGGCTTCGCCATGACTGAAGGCTTCAATTTTCCGCTGCTGG | *X. szentirmaii* DSM 16338 |
| | SW505* | AGCAGCGGAAAATTGAAGCCTTCAGTCATGGCGAAGCCG | *X. hominickii* DSM 17903 |
| pDD8 | SW506 | CCATACCCGTTTTTTTGGGCTAACAGGAGGAATTCCATGCAGGAAAATACAT TTCAATTTAAGGCATCCC | *X. budapestensis* DSM 16342 |
| | SW507 | CCGGCTTCGCCATGACTGAAACTTTCAATCTTCCGTTGTTGGATTATCGG | *X. budapestensis* DSM 16342 |
| | SW508/509* | AACAACGGAAGATTGAAAGTTTCAGTCATGGCGAAGCCG | *X. hominickii* DSM 17903 |

**Table S5.** Raw data refer to Fig. 2a of the γ-[$^{18}O_4$]-ATP *in vitro* assay.

| | **GxpS_A3-T3** | | **GxpS_C$_{ASub}$3-A3-T3** | | **GxpS_C3-A3-T3** | | **XtpS_C3-GxpS_A3-T3** | |
|---|---|---|---|---|---|---|---|---|
| | **MEAN %** | **SD %** | **MEAN %** | **SD %** | **MEAN %** | **SD %** | **MEAN %** | **SD %** |
| Ala | 6.0 | 1.7829 | 3.2 | 1.5923 | 6.5 | 0.8946 | 0.0 | - |
| Val | 13.1 | 1.5273 | 7.3 | 0.7999 | 28.1 | 4.1965 | 18.4 | 0.2795 |
| Met | 27.8 | 0.9616 | 21.7 | 2.4104 | 28.9 | 4.2377 | 22.2 | 0.2132 |
| Leu | 20.8 | 1.5242 | 1.5 | 2.1195 | 83.3 | 1.7940 | 80.2 | 1.3005 |
| Ile | 8.0 | 1.4162 | 1.5 | 1.8464 | 5.5 | 1.4720 | 4.5 | 0.5767 |
| Pro | 4.2 | 1.0358 | 1.7 | 1.7206 | 3.5 | 1.7304 | 1.3 | 0.2233 |
| Trp | 6.4 | 1.3769 | 4.9 | 4.3964 | 41.8 | 7.7732 | 36.2 | 1.1851 |
| Phe | 73.1 | 3.9659 | 4.1 | 1.3018 | 82.4 | 4.8753 | 86.2 | 0.7535 |
| Lys | 2.8 | 2.1504 | 0.4 | 0.6176 | 3.2 | 0.1311 | 2.0 | 0.1628 |
| Arg | 1.9 | 0.9436 | 4.5 | 3.3481 | 2.9 | 0.5006 | 1.3 | 0.1958 |
| His | 2.5 | 0.1231 | 1.1 | 1.6076 | 9.5 | 0.5304 | 4.2 | 0.2325 |
| Tyr | 22.8 | 0.9783 | 10.2 | 1.2426 | 82.9 | 0.3427 | 62.4 | 0.5753 |
| Thr | 1.6 | 2.2140 | 0.6 | 0.9052 | 4.4 | 0.4515 | 3.0 | 0.6157 |
| Gln | 2.0 | 2.8822 | 0.0 | - | 3.2 | 0.5854 | 1.5 | 0.6882 |
| Gly | 3.4 | 2.4223 | 0.0 | - | 2.3 | 0.6520 | 1.4 | 0.1089 |
| Ser | 2.4 | 3.4284 | 0.0 | - | 4.9 | 0.2329 | 1.7 | 0.4954 |
| Cys | 2.9 | 1.6624 | 3.1 | 4.4023 | 2.7 | 0.7036 | 1.4 | 0.1321 |
| Asn | 1.9 | 1.3335 | 1.0 | 1.3608 | 2.6 | 1.0490 | 2.7 | 0.1687 |
| Glu | 1.3 | 1.8734 | 0.4 | 0.6029 | 2.3 | 0.8902 | 0.9 | 0.7045 |
| Asp | 3.9 | 3.8562 | 0.3 | 0.3921 | 3.8 | 0.5405 | 1.3 | 0.3334 |
| PAPA | 96.8 | 2.2374 | 0.0 | - | 85.8 | 1.4628 | 86.2 | 0.5908 |

**Table S6.** Raw data refer to Fig. 2b-c of the multiplexed hydroxamate *in vitro* assay.

| | GxpS_A3-T3 | | GxpS_C$_{ASub}$3-A3-T3 | | GxpS_C3-A3-T3 | | XtpS_C3-GxpS_A3-T3 | | BacA_C3-GxpS_A3-T3 | |
|---|---|---|---|---|---|---|---|---|---|---|
| | **MEAN mg/l** | **SD mg/l** | **MEAN mg/l** | **SD mg/l** | **MEAN mg/l** | **SD mg/l** | **MEAN mg/l** | **SD mg/l** | **MEAN mg/l** | **SD mg/l** |
| Ala | - | - | - | - | - | - | - | - | - | - |
| Val | - | - | - | - | - | - | - | - | - | - |
| Met | - | - | - | - | - | - | - | - | - | - |
| Leu | 0.0757 | 0.0229 | - | - | 0.1080 | 0.0144 | 0.1080 | 0.0323 | - | - |
| Ile | 0.0757 | 0.0229 | - | - | 0.1080 | 0.0144 | 0.1080 | 0.0323 | - | - |
| Pro | - | - | - | - | - | - | - | - | - | - |
| Trp | 0.1050 | 0.0185 | - | - | 0.4880 | 0.0625 | 0.1360 | 0.0175 | - | - |
| Phe | 1.0300 | 0.1597 | 0.0504 | 0,0015 | 2.9100 | 0.3972 | 1.6500 | 0.2294 | 0.0280 | 0.0043 |
| Lys | - | - | - | - | - | - | - | - | - | - |
| Arg | - | - | - | - | - | - | - | - | - | - |
| His | - | - | - | - | - | - | - | - | - | - |
| Tyr | 0.0382 | 0.0037 | - | - | 0.1090 | 0.0171 | 0.0466 | 0.0106 | - | - |
| Thr | - | - | - | - | - | - | - | - | - | - |
| Gln | - | - | - | - | - | - | - | - | - | - |
| Gly | - | - | - | - | - | - | - | - | - | - |
| Ser | - | - | - | - | - | - | - | - | - | - |
| Cys | - | - | - | - | - | - | - | - | - | - |
| Asn | - | - | - | - | - | - | - | - | - | - |
| Glu | - | - | - | - | - | - | - | - | - | - |
| Asp | - | - | - | - | - | - | - | - | - | - |

**Table S7.** Raw data refer to Fig. 3 of the GxpS with deleted first two modules (A1 to C3) and varying preceding C domains *in vivo* assay.

| | GxpS_A3--TE | | GxpS_C3--TE | | GxpS_C$_{ASub}$3--TE | | XtpS_C3-GxpS_A3--TE | | KolS_C5-GxpS_A3--TE | | AmbS$_{indica}$_C5-GxpS_A3--TE | | AmbS$_{mira}$_C5-GxpS_A3--TE | | BicA_C3-GxpSA3--TE | |
|---|---|---|---|---|---|---|---|---|---|---|---|---|---|---|---|---|
| | **MEAN mg/l** | **SD mg/l** | **MEAN mg/l** | **SD mg/l** | **MEAN mg/l** | **SD mg/l** | **MEAN mg/l** | **SD mg/l** | **MEAN mg/l** | **SD mg/l** | **MEAN mg/l** | **SD mg/l** | **MEAN mg/l** | **SD mg/l** | **MEAN mg/l** | **SD mg/l** |
| AIL | - | - | - | - | - | - | - | - | - | - | - | - | - | - | - | - |
| VIL | - | - | - | - | - | - | - | - | - | - | - | - | - | - | - | - |
| MIL | 0.4510 | 0.0598 | 0.0711 | 0.0105 | 0.0304 | 0.0115 | 0.0412 | 0.0094 | 0.0222 | 0.0113 | - | - | 0.0740 | 0.0066 | 0.0933 | 0.0077 |
| I/LIL | 2.7240 | 0.3624 | 1.5264 | 0.1965 | 1.2375 | 0.1136 | 0.9644 | 0.1902 | 0.5896 | 0.2422 | 0.3275 | 0.0102 | 2.8215 | 0.0902 | 3.5202 | 0.2752 |
| i/lIL | 0.1758 | 0.0247 | 0.0792 | 0.0161 | 0.0824 | 0.0057 | 0.0681 | 0.0120 | 0.0398 | 0.0296 | - | - | 0.1868 | 0.0175 | 0.2273 | 0.0234 |
| PIL | 0.0038 | 0.0085 | 0.4944 | 0.0619 | 0.0919 | 0.0418 | 0.4964 | 0.1275 | 0.0107 | 0.0815 | 0.0467 | 0.0047 | 0.1606 | 0.0382 | 0.0683 | 0.0244 |
| WIL | 0.1516 | 0.0126 | 0.0805 | 0.0078 | 0.0772 | 0.0371 | 0.0663 | 0.0164 | 0.0753 | 0.0355 | 0.0255 | 0.0013 | 0.2334 | 0.0010 | 0.3548 | 0.0199 |
| FIL | 9.6263 | 0.5551 | 14.4737 | 1.1411 | 5.7133 | 1.9941 | 10.5606 | 1.5582 | 3.6512 | 2.8903 | 1.8598 | 0.0457 | 18.3480 | 0.9115 | 20.2288 | 1.5753 |
| fIL | 5.3513 | 0.5291 | 6.9523 | 0.9291 | 2.8380 | 0.4718 | 6.0191 | 1.2855 | 1.8271 | 1.6909 | 0.9494 | 0.0735 | 9.5853 | 0.3810 | 9.9711 | 0.6059 |
| KIL | - | - | - | - | - | - | - | - | - | - | - | - | - | - | - | - |
| RIL | - | - | - | - | - | - | - | - | - | - | - | - | - | - | - | - |
| HIL | - | - | - | - | - | - | - | - | - | - | - | - | - | - | - | - |
| YIL | 0.3356 | 0.1154 | 0.1749 | 0.0217 | 0.1289 | 0.0265 | 0.1428 | 0.0230 | 0.1197 | 0.0818 | 0.0543 | 0.0039 | 0.5044 | 0.0175 | 0.6745 | 0.0432 |
| TIL | - | - | - | - | - | - | - | - | - | - | - | - | - | - | - | - |
| QIL | - | - | - | - | - | - | - | - | - | - | - | - | - | - | - | - |
| GIL | - | - | - | - | - | - | - | - | - | - | - | - | - | - | - | - |
| SIL | - | - | - | - | - | - | - | - | - | - | - | - | - | - | - | - |
| CIL | - | - | - | - | - | - | - | - | - | - | - | - | - | - | - | - |
| NIL | - | - | - | - | - | - | - | - | - | - | - | - | - | - | - | - |
| EIL | - | - | - | - | - | - | - | - | - | - | - | - | - | - | - | - |
| DIL | - | - | - | - | - | - | - | - | - | - | - | - | - | - | - | - |

**Table S8.** HeatMap of the Clustal Omega alignment of C domains investigated in this work. Higher similarity is represented by darker plot colors.

| Identity [%] | GxpS_C3 | XtpS_C3 | BacA_C3 | KolS_C5 | AmbS$_{X. indi.}$_C5 | AmbS$_{X. mira.}$_C5 | BicA_C3 | *Xsze* FclJ_C6 | *Xbud* FclJ_C6 | *Xhom* FclJ_C6 |
|---|---|---|---|---|---|---|---|---|---|---|
| **GxpS_C3** | | 86.33 | 41.38 | 67.29 | 64.43 | 63.87 | 62.10 | 27.52 | 28.15 | 28.22 |
| **XtpS_C3** | 86.33 | | 41.38 | 68.86 | 64.35 | 65.05 | 63.04 | 28.59 | 28.33 | 28.51 |
| **BacA_C3** | 41.38 | 41.38 | | 41.80 | 39.96 | 42.34 | 42.03 | 27.69 | 28.56 | 29.42 |
| **KolS_C5** | 67.29 | 68.86 | 41.80 | | 69.95 | 71.14 | 70.72 | 27.91 | 28.56 | 28.45 |
| **AmbS$_{X. indi.}$_C5** | 64.43 | 64.35 | 39.96 | 69.95 | | 67.26 | 65.10 | 30.66 | 30.79 | 30.45 |
| **AmbS$_{X. mira.}$_C5** | 63.87 | 65.05 | 42.34 | 71.14 | 67.26 | | 70.40 | 28.03 | 29.07 | 27.82 |
| **BicA_C3** | 62.10 | 63.04 | 42.03 | 70.72 | 65.10 | 70.40 | | 27.69 | 28.56 | 29.42 |
| ***Xsze* FclJ_C6** | 27.52 | 28.59 | 27.69 | 27.91 | 30.66 | 28.03 | 27.69 | | 74.59 | 73.91 |
| ***Xbud* FclJ_C6** | 28.15 | 28.33 | 28.56 | 28.56 | 30.79 | 29.07 | 28.56 | 74.59 | | 80.54 |
| ***Xhom* FclJ_C6** | 28.22 | 28.51 | 29.42 | 28.45 | 30.45 | 27.82 | 29.42 | 73.91 | 80.54 | |

**Table S9.** The selectivity-conferring code (***Stachelhaus et al., 1999***) of the FclJ A6 domains and antiSMASH (***Blin et al., 2021***) substrate prediction.

| FclJ A6 | AA1 | AA2 | AA3 | AA4 | AA5 | AA6 | AA7 | AA8 | AA9 | AA10 | antiSMASH prediction |
|---|---|---|---|---|---|---|---|---|---|---|---|
| ***Xbud*** | D | A | W | F | I | G | G | H | - | K | Leu |
| ***Xhom*** | D | A | L | F | I | G | G | Y | -(E) | K | Leu |
| ***Xsze*** | D | A | L | F | I | G | G | Y | -(E) | K | Leu |

**Table S10.** RMSD to mean of the MOE homology models.

| | Template | | | |
|---|---|---|---|---|
| **model** | **2VSQ** | **4ZXH** | **5T3D** | |
| *Xsze* FclJ_C6-*Xhom.* FclJ_A6 | 1.26 | 1.75 | 1.47 | **RMSD [Å]** |
| *Xsze* FclJ_C6-*Xbud.* FclJ_A6 | 1.27 | 1.68 | 1.52 | |
| *Xsze.* FclJ_C6-A6 | 1.49 | 1.86 | 1.77 | |
| *Xhom.* FclJ_C6-A6 | 1.29 | 1.67 | 1.50 | |
| *Xbud.* FclJ_C6-A6 | 1.30 | 1.70 | 1.47 | |
| BacA_C3-GxpS_A3 | 1.14 | 1.69 | 1.57 | |
| BacA_C3-A3 | 1.31 | 1.77 | 1.52 | |
| XtpS_C3-GxpS_A3 | 1.11 | 1.67 | 1.39 | |
| XtpS_C3-A3 | 1.35 | 1.68 | 1.48 | |
| GxpS_C3-A3 | 1.13 | 1.60 | 1.28 | |

**Table S11.** Clustal Omega alignment of the C-A didomains used for homology modeling and in HSPred analysis including the AB3403 (PDB-ID: 4ZXH), EntF (PDB-ID: 5T3D) and SrfA-C (PDB-ID: 2SVQ) templates. The interacting Chains I - XVI are highlighted in grey and entitled below. Colouring of the consensus sequence is according to the C domain (green), C-A linker (blue), $A_{Core}$ (red), and $A_{Sub}$ (orange).

```
                                       10        20        30        40        50        60
Consensus                   LSRMQQGMXYHMNLXPDXNVYHXX--TXLRVNGRXDEXAFREALAQTVARHDVLRTAFDM   58
2VSQ.A                      LSPMQEGMLFHAILNPGQSFYLEQ--ITMKVKGSLNIKCLEESMNVIMDRYDVFRTVFIH   58
5T3D.A                      LVAAQPGIWMAEKLSELPSAWSVA--HYVELTGEVDSPLLARAVVAGLAQADTLRMRFTE   58
4ZXH.A                      ISSEQLGIWYIQRLEPTCSAYNMVVAFDVKVNQSLGNKPI-EILEAVMHDYPLLRVSMPA   59
BacA_C3-A3                  TSPAQQRMYMLSMLENERGAYHIP--MALLVEGRINAMQLENALKTFLQRHEILRTGFEI   58
BacA_C3-GxpS_A3             TSPAQQRMYMLSMLENERGAYHIP--MALLVEGRINAMQLENALKTFLQRHEILRTGFEI   58
Xbud FclJ_C6-A6             LSRMQAGMFYHMNMSPDANVYHCTGTSHLRVNGRFDEKAFIEATAQTVAAHDVLRTAFDM   60
Xhom FclJ_C6-A6             FSRMQEGMFYHMDLSPESNVYHCTGTSHLRINGPFDERAFREATAQTVAAHDVLRTAFDM   60
Xsze FclJ_C6-A6             LSRMQAGMFYHMNLSPDANVYHCTGSTHIFINGQFDETAFREATAQTVAAHDVLRTAFDM   60
Xsze FclJ_C6-Xbud FclJ_A6   LSRMQAGMFYHMNLSPDANVYHCTGSTHIFINGQFDETAFREATAQTVAAHDVLRTAFDM   60
Xsze FclJ_C6-Xhom FclJ_A6   LSRMQAGMFYHMNLSPDANVYHCTGSTHIFINGQFDETAFREATAQTVAAHDVLRTAFDM   60
GxpS_C3-A3                  LSFGQQRLWFLAQFEGVSDTYHIP--TALRLRGRLNFSAWQQALNRLFARHEALRSVFIT   58
XtpS_C3-A3                  LSFGQQRLWFLAQFEGISDTYHIP--TALRLRGRLDLSAWQQALNCLFARHEALRSVFIT   58
XtpS_C3-GxpS_A3             LSFGQQRLWFLAQFEGISDTYHIP--TALRLRGRLDLSAWQQALNCLFARHEALRSVFIT   58

                                       70        80        90       100       110       120
Consensus                   EN-SEPLQLVLEFAEFPJVVEDL-RHLSDEEQDEQXKALLEXERVNPFDLLRPPLLRFFJ  116
2VSQ.A                      EKVKRPVQVVLKKRQFHIEEIDL-THLTGSEQTAKINEYKEQDKIRGFDLTRDIPMRAAI  117
5T3D.A                      DN-GEVWQWVDDALTFELPEIIDLRTNIDPHGTAQALMQADLQQDLRVDSGKP-LVFHQL  116
4ZXH.A                      ND--QGIEQLIWDRVYPNIIFSDARHIEAS----DLTQLVEQDTKQPFDLTQPPLWRIHC  113
BacA_C3-A3                  QN-NELIQKIYENVDFRLEYECLDASITDQHALMEITSRYCKESIKPFDLSRPPLMRAKL  117
BacA_C3-GxpS_A3             QN-NELIQKIYENVDFRLEYECLDASITDQHALMEITSRYCKESIKPFDLSRPPLMRAKL  117
Xbud FclJ_C6-A6             ENYSEPLQLVYQFAELPIVVEDL-RHLSEEEQDERAKALLEAERVNPFDLLNPTLLRFFI  119
Xhom FclJ_C6-A6             ENYSEPLQLVHEFAELPIVVEDL-RHLSVEEQDDQAMALLEAERVNPFDLLKPTLLRFFI  119
Xsze FclJ_C6-A6             ESYSEPLQLVLEFAELPIVVEDL-RHLSDEEQDERVKALLEAERVNPFDLLNPTLLRFFI  119
Xsze FclJ_C6-Xbud FclJ_A6   ESYSEPLQLVLEFAELPIVVEDL-RHLSDEEQDERVKALLEAERVNPFDLLNPTLLRFFI  119
Xsze FclJ_C6-Xhom FclJ_A6   ESYSEPLQLVLEFAELPIVVEDL-RHLSDEEQDERVKALLEAERVNPFDLLNPTLLRFFI  119
GxpS_C3-A3                  VD-GQPQVELLP-AEFGLPVKKYDLRKASN-VDKQFKRLCVQEAKTPFDLARGPLIRCAL  115
XtpS_C3-A3                  VE-GQPQVELLP-AEFALPMQKYDLRKASD-VDEQLAHLCAQEAKTPFDLARGPLIRCAL  115
XtpS_C3-GxpS_A3             VE-GQPQVELLP-AEFALPMQKYDLRKASD-VDEQLAHLCAQEAKTPFDLARGPLIRCAL  115

                                      130       140       150       160       170       180
Consensus                   XLRXDXSFQFTMTEHHIIXDGWSYGILIVEVFXRYATLTGQXPWPXPXK-SLEYRDYIAW  175
2VSQ.A                      FKKAEESFEWVWSYHHIILDGWCFGIVVQDLFKVYNALREQKPYSLPPV--KPYKDYIKW  175
5T3D.A                      IQVADNRWYWYQRYHHLLVDGFSFPAITRQIANIYCTWLRGEPTPASPF--TPFADVVEE  174
4ZXH.A                      YECGQNHYVIAFVIHHALMDFWSIGLLLRDVSKRFGLVA-----ESDAVNGIEFVQYADK  168
BacA_C3-A3                  IKIDDIRHILVINFHHIISDGVSQGILMNEILELYSNV------PLPEV-NVQYKDYVEW  170
BacA_C3-GxpS_A3             IKIDDIRHILVINFHHIISDGVSQGILMNEILELYSNV------PLPEV-NVQYKDYVEW  170
Xbud FclJ_C6-A6             QLRTDNSFQFTMTECHPVFDGWSYNTMIVEVFNRYATLTGQASWQAPDKKSLEYRDFIAL  179
Xhom FclJ_C6-A6             QLRTDNSFQFTMTECHPVFDGWSYNTMIVEVFNRYATLTGQVNWQAPDKKSLEYRDFIAL  179
Xsze FclJ_C6-A6             YLRTDHSFQFTMTECHPIFDGWSYNTMIVEVFSRYATLTGLDNWQAPDKKSLEYRDFIAL  179
Xsze FclJ_C6-Xbud FclJ_A6   YLRTDHSFQFTMTECHPIFDGWSYNTMIVEVFSRYATLTGLDNWQAPDKKSLEYRDFIAL  179
Xsze FclJ_C6-Xhom FclJ_A6   YLRTDHSFQFTMTECHPIFDGWSYNTMIVEVFSRYATLTGLDNWQAPDKKSLEYRDFIAL  179
GxpS_C3-A3                  IQLADEEHIFLLTQHHIISDAWSLKILESELVALYAAGLNQQPDPLLPL-TIQYPDYAAW  174
XtpS_C3-A3                  VRLAEEEHLFLLTQHHIISDGWSLGILESELTALYVACLNQQPDPLPPL-TIQYPDYAAW  174
XtpS_C3-GxpS_A3             VRLAEEEHLFLLTQHHIISDGWSLGILESELTALYVACLNQQPDPLPPL-TIQYPDYAAW  174

                                      190       200       210       220       230       240
Consensus                   EQAAIADEKXKMXWAYXLD--TJXXLPRKLXXPVDRDRPSX-------------------  214
2VSQ.A                      LEKQDKQASLRYWREY------LEGFEGQTTFAEQRKKQK--------------------  209
5T3D.A                      YQQYRESEAWQRDAAFWAEQRRQLPPPASLSPAPLPGR----------------------  212
4ZXH.A                      QQSSVIDDTDESLIFWKN---ALKHAPHVHSIPLDYPRPAV-------------------  206
BacA_C3-A3                  NHTFNQSAAMKKQEAYWLD--VYRDIPSKLDFPYDYKRHHI-------------------  209
BacA_C3-GxpS_A3             NHTFNQSAAMKKQEAYWLD--VYRDIPSKLDFPYDYKRHHI-------------------  209
Xbud FclJ_C6-A6             EQAAIADEKQKDYWAQKLDDCTILELPRKQHTDGINDVVASA------------------  221
Xhom FclJ_C6-A6             EQAAIADEQQKAYWTQKLDDCTILELPRKQHTDVIDDTGVPA------------------  221
Xsze FclJ_C6-A6             EKAAIADEKNKMYWAQKLDDCTIMELPRKLNAGVSGDAASHAGEGTVDGNNGDDNDSGTL  239
Xsze FclJ_C6-Xbud FclJ_A6   EKAAIADEKNKMYWAQKLDDCTIMELPRKLNAGVSGDAASHAGEGTVDGNNGDDNDSGTL  239
Xsze FclJ_C6-Xhom FclJ_A6   EKAAIADEKNKMYWAQKLDDCTIMELPRKLNAGVSGDAASHAGEGTVDGNNGDDNDSGTL  239
GxpS_C3-A3                  QRQVFSGEGVQAQSDYWRR--TLADAPVLLELPIDRPRPSQ-------------------  213
XtpS_C3-A3                  QRQVFSDEGVQVQSDYWRR--TLADAPVLLELPLDRPRPSQ-------------------  213
XtpS_C3-GxpS_A3             QRQVFSDEGVQVQSDYWRR--TLADAPVLLELPLDRPRPSQ-------------------  213
                                                      Chain I             Chain II

                                      250       260       270       280       290       300
Consensus                   -XSXKGKSXXLXLEXTXYQGLRHLMQQLGVTLKSVLLTGHIKLLSXXSGEHDIVTGIPAA  273
2VSQ.A                      -DGYEPKELLFSPSEAETKAFTELAKSQHTTLSTALQAVWSVLISRYQQSGDLAFGTVVS  268
5T3D.A                      --SASADILRLKLEFTDGEFRQLATQLSGVQRTDLALALAALWLGRLCNRMDYAAGFIFM  270
4ZXH.A                      -QQHKGSSLVFRVSESVSSGLVNLAKDYEITLFGLVLSGFYVLLHKLSNENNLVIATPVA  265
BacA_C3-A3                  -DTFEGSSVFLEMERELSDHIRKLAKHNGTTLYTVMLSAYYVLLNKYTNQTDIVVGTAAA  268
BacA_C3-GxpS_A3             -DTFEGSSVFLEMERELSDHIRKLAKHNGTTLYTVMLSAYYVLLNKYTNQTDIVVGTAAA  268
Xbud FclJ_C6-A6             -TSPKLKSFSLILDSTVYQGLRHLMQQLGVPLKSILLTGHIKVMSIFSGEHDILTGITAN  280
Xhom FclJ_C6-A6             ANRPKLKSFSLILDTHIYQGLRHLMQQLSVPMKSLLLTGHIKVMSIFSGENDILTGITAN  281
```

```
Xsze FclJ_C6-A6              ATSPKIKSFTLTLEGTVYQGLRHLMQQLGVPLKSVLLTGHIKVMSIFSGEHDILTGISTN  299
Xsze FclJ_C6-Xbud FclJ_A6    ATSPKIKSFTLTLEGTVYQGLRHLMQQLGVPLKSVLLTGHIKVMSIFSGEHDILTGISTN  299
Xsze FclJ_C6-Xhom FclJ_A6    ATSPKIKSFTLTLEGTVYQGLRHLMQQLGVPLKSVLLTGHIKVMSIFSGEHDILTGISTN  299
GxpS_C3-A3                   -QSFTGGRVAVHIDAPLVQALKHLGQQHGATLFMTLLTAWAALLSRLSGQDDVVIGIPSA  272
XtpS_C3-A3                   -QSFAGGQVAVHFDATLVQDLKHLGQQHGTTLFMTLLTAWATLLSRLSGQDDVVIGIPSA  272
XtpS_C3-GxpS_A3              -QSFAGGQVAVHFDATLVQDLKHLGQQHGTTLFMTLLTAWATLLSRLSGQDDVVIGIPSA  272
                                                                      Chain III

                                       310       320       330       340       350       360
Consensus                    GRP-EAQG-DNLYGLFXNTLPLRQILSPV-SWHELLRQVFANXIEAIPHQRYPXAEIQRQ  330
2VSQ.A                       GRPAEIKGVEHMVGLFINVVPRRVKLSEGITFNGLLKRLQEQSLQSEPHQYVPLYDIQSQ  328
5T3D.A                       RRLGSAAL--TATGPVLNVLPLGIHIAAQETLPELATRLAAQLKKMRRHQRYDAEQIVRD  328
4ZXH.A                       GRL--ERSLRNALGQFVNTIAIHMDIDADQTLRQFTQQVQEQLRQSLKHQKIAFSRVVEA  323
BacA_C3-A3                   GRL-HPDL-QDVFGVFVNTLALRNEVDTSYSFKEFLQQTKERTIAAFDNSEYPFDDLIRK  326
BacA_C3-GxpS_A3              GRL-HPDL-QDVFGVFVNTLALRNEVDTSYSFKEFLQQTKERTIAAFDNSEYPFDDLIRK  326
Xbud FclJ_C6-A6              GRP-EAQGGDHLYGLFLNILPFRQILTPV-SWHELIRQVFANEIEAIPYRRYPLAEIQRQ  338
Xhom FclJ_C6-A6              GRP-EAQGGDKLYGLFLNILPFRQILTPV-SWNELIRQVFANEIEAIPYRRYPLAEIQRQ  339
Xsze FclJ_C6-A6              GRP-EAQGGDNLYGLFLNILPFRQILSPV-SWHELIRQVFANEIEAIPHRRYPLAEIQRQ  357
Xsze FclJ_C6-Xbud FclJ_A6    GRP-EAQGGDNLYGLFLNILPFRQILSPV-SWHELIRQVFANEIEAIPHRRYPLAEIQRQ  357
Xsze FclJ_C6-Xhom FclJ_A6    GRP-EAQGGDNLYGLFLNILPFRQILSPV-SWHELIRQVFANEIEAIPHRRYPLAEIQRQ  357
GxpS_C3-A3                   NRN-LREI-EPLLGFFVNTLALRIDLSDMPDVATLLRRVRQTTLGAQEHQDLPFEQVVEI  330
XtpS_C3-A3                   NRN-RREI-ESLLGFFVNTLALRIDLSGMPDVVTLLQRVRQTTLGAQEHQDLPFEQVVEI  330
XtpS_C3-GxpS_A3              NRN-RREI-ESLLGFFVNTLALRIDLSGMPDVVTLLQRVRQTTLGAQEHQDLPFEQVVEI  330
                                                  Chain IV

                                       370       380       390       400       410       420
Consensus                    FGQQRLLDEVPLFYIXFHIYDQMAAEXXLNVXXXLXTQDVYEGXXFDL--------XVHF  382
2VSQ.A                       ADQPKLIDHIIVFE-NYPLQDAKNEESSENGFDMVDVH-VFEKSNYDL----------NL  376
5T3D.A                       SGRAAGDEPLFGPVLNIKVFDYQLDIPDVQAQTHTLATGPVNDLELAL-----------F  377
4ZXH.A                       VSPKRDGSINPLAQIGMFWERLGGMDEFKELLLPIQTPATLVGQDLTLGSFPVRQQEGQL  383
BacA_C3-A3                   LNGVRESNRNPLFDTMFVLEDARMFTKQKGDVKLSPIIFELDNAKFDM--------IFNV  378
BacA_C3-GxpS_A3              LNGVRESNRNPLFDTMFVLEDARMFTKQKGDVKLSPIIFELDNAKFDM--------IFNV  378
Xbud FclJ_C6-A6              FGQQPLLDEVLFNYIDFHIYDQMVPEVGLEVVDRLHTQDVYEGTHFTL--------DVHF  390
Xhom FclJ_C6-A6              FGQQPLLDEVLFNYIDFHIYDQMAPEIGLEVVDRLHTQDVYEGTHFTL--------DIHF  391
Xsze FclJ_C6-A6              FGQQPLLDEVLFNYIDFHIYDQMAAGLGLNVIGKLHTQDVYEGTHFAL--------TVHF  409
Xsze FclJ_C6-Xbud FclJ_A6    FGQQPLLDEVLFNYIDFHIYDQMAAGLGLNVIGKLHTQDVYEGTHFAL--------TVHF  409
Xsze FclJ_C6-Xhom FclJ_A6    FGQQPLLDEVLFNYIDFHIYDQMAAGLGLNVIGKLHTQDVYEGTHFAL--------TVHF  409
GxpS_C3-A3                   VQPPRRPEHTPLFQVMFAWQESETEEWQLPELAVTPFELGYDIAKFDL--------QLEL  382
XtpS_C3-A3                   VQPPRRPEHTPLFQVMFAWQESETKEWQLPELAVTPFELGYDIAKFDL--------QLEL  382
XtpS_C3-GxpS_A3              VQPPRRPEHTPLFQVMFAWQESETKEWQLPELAVTPFELGYDIAKFDL--------QLEL  382
                               Chain V

                                       430       440       450       460       470       480
Consensus                    QHKT-------DZIXIQIDYXENLFSRETIARMAECYXAVLXAMVAEPQ-XLHCIDXFSP  434
2VSQ.A                       MASP------GDEMLIKLAYNENVFDEAFILRLKSQLLTAIQQLIQNPDQPVSTINLVDD  430
5T3D.A                       PDVH-------GDLSIEILANKQRYDEPTLIQHAERLKMLIAQFAADPALLCGDVDIMLP  430
4ZXH.A                       DITLEMGGEYQGELVGVLKYNTDLFSAQSAENMVQLLQAVLSEMVAHPERKIVELDIAPD  443
BacA_C3-A3                   LDFE-------QKIVLNIEYSTSLFKDETIQKIAEDYFRILEEVSENLDVALHQIDMISR  431
BacA_C3_GxpS_A3              LDFE-------QKIVLNIEYSTSLFKDETIQKIAEDYFRILEEVSENLDVALHQIDMISR  431
Xbud FclJ_C6-A6              QHLTLSSALASDQISVQLDYDENKISRGLAANMAECYAAVFAAMVADPQ-ALHCASHFLP  449
Xhom FclJ_C6-A6              QHLTLSSALASDQISVQIDYDETKLSRGQVANMAECYSAVFASMVAEPQ-ALHCASHFLP  450
Xsze FclJ_C6-A6              QHKTLTSSLINDQVSIQIDYDENRLSRDLAADMAECYSAVFTAMVTEAQ-SLHCANHFVP  468
Xsze FclJ_C6-Xbud FclJ_A6    QHKTLTSSLINDQVSIQIDYDENRLSRDLAADMAECYSAVFTAMVTEAQ-SLHCANHFVP  468
Xsze FclJ_C6-Xhom FclJ_A6    QHKTLTSSLINDQVSIQIDYDENRLSRDLAADMAECYSAVFTAMVTEAQ-SLHCANHFVP  468
GxpS_C3-A3                   TEKA-------GEIVGELNYSSALFDHETIERQVGYLQAILWAIVNQPQQPVTAIDILSS  435
XtpS_C3-A3                   TEKA-------GEIVGVLNYSSALFDHETIERQMGYLQAILRAMVNQPQQPVAVIDILSS  435
XtpS_C3-GxpS_A3              TEKA-------GEIVGVLNYSSALFDHETIERQMGYLQAILRAMVNQPQQPVAVIDILSS  435
                                                    Chain VI                   Chain VII

                                       490       500       510       520       530       540
Consensus                    MEQQRLLEXXNATEEGYPXQVTLXQLFAEQXAKTPDACAVXY----GNQTLSYLELNQQA  490
2VSQ.A                       REREFLLTGLNPPAQAHETKP-LTYWFKEAVNANPDAPALTY----SGQTLSYRELDEEA  485
5T3D.A                       GEYAQLAQ-LNATQVEIP-ETTLSALVAEQAAKTPDAPALAD----ARYLFSYREMREQV  484
4ZXH.A                       YKDGIQFEALRGKATDYA-QHDLFAMILKQIDERGDNHALTS----NDHTVSYRELGQHI  498
BacA_C3-A3                   QEKRTLLESWNATEEPYPTQVCVHQLFEQQIEKTPDAIAVIY----ENQTLSYAELNARA  487
BacA_C3-GxpS_A3              QEKRTLLESWNATEEPYPTQVCVHQLFEQQIEKTPDAIAVIY----ENQTLSYAELNARA  487
Xbud FclJ_C6-A6              IIQQRKIESFNQGAPGYQGQETLAALFAEQAARTPSACAVEY----GDRQLSYLELHQQS  505
Xhom FclJ_C6-A6              VTQQRLIESFSHGEAGYQGQATLADLFAEQVARSPSACVVEL----GERQLSYLALHQQS  506
Xsze FclJ_C6-A6              MAQQRKIEAFSQGASGYRGQQTLAELFAEQVARSPSACAVEFSKPSGKQQLTYLALDQQS  528
Xsze FclJ_C6-Xbud FclJ_A6    MAQQRKIEAFNQGAPGYQGQETLAALFAEQAARTPSACAVEY----GDRQLSYLELHQQS  524
Xsze FclJ_C6-Xhom FclJ_A6    MAQQRKIEAFSHGEAGYQGQATLADLFAEQVARSPSACVVEL----GERQLSYLALHQQS  524
GxpS_C3-A3                   SERELLLENWNATEEPYPTQVCVHQLFEQQIEKTPDAIAVIY----ENQTLSYAELNARA  491
XtpS_C3-A3                   TERTLLLKTWNATETVYPESLCIHQLFEQQVEKTPQATALIA----GEKHLSYSELNVWA  491
XtpS_C3-GxpS_A3              TERTLLLKTWNATEEPYPTQVCVHQLFEQQIEKTPDAIAVIY----ENQTLSYAELNARA  491
                               Chain VII

                                       550       560       570       580       590       600
Consensus                    NRLAHLLIXKGVKPGQRVALLLGRSIELIIXJLAVLKAGAAYLPLPDPEXPDERLXXMJED  550
2VSQ.A                       NRIARRLQKHGAGKGSVVALYTKRSLELVIGILGVLKAGAAYLPVDPKLPEDRISYMLAD  545
5T3D.A                       VALANLLRERGVKPGDSVAVALPRSVFLTLALHAIVEAGAAWLPLDTGYPDDRLKMMLED  544
4ZXH.A                       AGIAEYLRAHGITQGDRVGLMLDRTALLPAAILGIWAAGAAYVPLDPNFPTERLQNIIED  558
BacA_C3-A3                   NRLAHQLIALGVAPDQRVAICVTRSLARIIGLLAVLKAGGAYVPLDPAYPGERLAYMLTD  547
BacA_C3-GxpS_A3              NRLAHQLIALGVAPDQRVAICVTRSLARIIGLLAVLKAGGAYVPLDPAYPGERLAYMLTD  547
Xbud FclJ_C6-A6              NQLAHLLAQKGVKQGQYVALLLGRSIELVISMLALVKLGAVYLPLNTEDPDIRLIEQIED  565
Xhom FclJ_C6-A6              NQLAHLLAQKGVKPGQRVALLLGRSIELIVSIMALAKLEAVYLPLNTEDPAARLIEQIED  566
Xsze FclJ_C6-A6              NQLAHLLVQKGVQQGQRVALLLGRSIELIVSILALVKLKAVYLPLNPEDPDARLLEQVED  588
Xsze FclJ_C6-Xbud FclJ_A6    NQLAHLLAQKGVKQGQYVALLLGRSIELVISMLALVKLGAVYLPLNTEDPDIRLIEQIED  584
Xsze FclJ_C6-Xhom FclJ_A6    NQLAHLLAQKGVKPGQRVALLLGRSIELIVSIMALAKLEAVYLPLNTEDPAARLIEQIED  584
GxpS_C3-A3                   NRLAHQLIALGVAPDQRVAICVTRSLARIIGLLAVLKAGGAYVPLDPAYPGERLAYMLTD  551
```

```
XtpS_C3-A3              NRLARQLIGQGVGSGDHIALLFERSIKLVVAQLAVLKAGAVYVPLDPDMPDGRKNWLIND  551
XtpS_C3-GxpS_A3         NRLAHQLIALGVAPDQRVAICVTRSLARIIGLLAVLKAGGAYVPLDPAYPGERLAYMLTD  551

                                  610       620       630       640       650       660
Consensus               AQPVLLXTDXRXTAALXEXILATLTXL-----DQSTLXEXPVXDLQXSG------TPNXP  599
2VSQ.A                  SAAACLLTHQEMKEQAAELPYTGTTLFI----DDQTRFEEQASDPATAI------DPNDP  595
5T3D.A                  ARPSLLITTDDQLPRFSDVPNLTSLCY------NAPLTPQGSAPLQLSQ-------PHHT  591
4ZXH.A                  AEPKVILTQTELMDGLNVSVPRLDI-------NQAGVVALEQVRETLAF--------GDI  603
BacA_C3-A3              ATPVILMADNVGRAALSEDILATLTVL-----DPNTLLEQPDHNPQVSG-----LTPQHL  597
BacA_C3-GxpS_A3         ATPVILMADNVGRAALSEDILATLTVL-----DPNTLLEQPDHNPQVSG-----LTPQHL  597
Xbud FclJ_C6-A6         TQCDLLIIDRRITDRLLPSISPHLPPILWLDEEQSALAQMPVTDLPDNLSHHAENQANLP  625
Xhom FclJ_C6-A6         TQCHLLVSDRRITNSLLSLTEMSLPPVLWLDAEQSALAQMPVTDLTLSTNNQT---TNLP  623
Xsze FclJ_C6-A6         AQCEFFITDQRVIDSSLPMKAANLSAVIWLDAEQSSLSRMPVSPLPDKGSHPDENQANLP  648
Xsze FclJ_C6-Xbud FclJ_A6  TQCDLLIIDRRITDRLLPSISPHLPPILWLDEEQSALAQMPVTDLPDNLSHHAENQANLP  644
Xsze FclJ_C6-Xhom FclJ_A6  TQCHLLVSDRRITNSLLSLTEMSLPPVLWLDAEQSALAQMPVTDLTLSTNNQT---TNLP  641
GxpS_C3-A3              ATPVILMADNVGRAALSEDILATLTVL-----DPNTLLEQPDHNPQVSG-----LTPQHL  601
XtpS_C3-A3              CAAKLLLTD-IQTAIPTDLIVPLLRLS-----DESEAVSEQASGQKNSTDLDLPRTSTEL  605
XtpS_C3-GxpS_A3         ATPVILMADNVGRAALSEDILATLTVL-----DPNTLLEQPDHNPQVSG-----LTPQHL  601

                                  670       680       690       700       710       720
Consensus               AYVMYTSGSTGKPKGVLIGHXXIIXL-XKDVSYXDFCTFGRXLQLASVSFDASTWEIWGP  658
2VSQ.A                  AYIMYTSGTTGKPKGNITTHANIQGL-VKHVDYMAFSDQDTFLSVSNYAFDAFTFDFYAS  654
5T3D.A                  AYIIFTSGSTGRPKGVMVGQTAIVNRLLWMQNHYPLTGEDVVAQKTPCSFDVSVWEFFWP  651
4ZXH.A                  AYVMYTSGSTGKPKGVRIGHPSIINFLLSMNDRLQVTTETQLLAITTYAFDISILELLIP  663
BacA_C3-A3              AYVIYTSGSTGRPKGVMIEHRSVVNLTLTQITQFDVCATSRMLQFASFGFDASVWEIMMA  657
BacA_C3-GxpS_A3         AYVIYTSGSTGRPKGVMIEHRSVVNLTLTQITQFDVCATSRMLQFASFGFDASVWEIMMA  657
Xbud FclJ_C6-A6         AYVMYTSGSTGKPKGALIGQKGIIRL-VKDVSYIDFQTFGRCLQLASVSFDASTWEIWGP  684
Xhom FclJ_C6-A6         AYIMYTSGSTGKPKGALIGQKGIIRL-VKDVSYIDFHTFGRFLQLAAVSFDASTLEIWGP  682
Xsze FclJ_C6-A6         AYVMYTSGSTGKPKGALIGQKGIIRL-VKDVSYADFRVFGRFLQLASVNFDASTLEIWGP  707
Xsze FclJ_C6-Xbud FclJ_A6  AYVMYTSGSTGKPKGALIGQKGIIRL-VKDVSYIDFQTFGRCLQLASVSFDASTWEIWGP  703
Xsze FclJ_C6-Xhom FclJ_A6  AYIMYTSGSTGKPKGALIGQKGIIRL-VKDVSYIDFHTFGRFLQLAAVSFDASTLEIWGP  700
GxpS_C3-A3              AYVIYTSGSTGRPKGVMIEHRSVVNLTLTQITQFDVCATSRMLQFASFGFDASVWEIMMA  661
XtpS_C3-A3              AYIMYTSGSTGTPKGVLVPHRAVARLVINN-GYAAIEPDDRVAFTANPAFDASTFDVWAP  664
XtpS_C3-GxpS_A3         AYVIYTSGSTGRPKGVMIEHRSVVNLTLTQITQFDVCATSRMLQFASFGFDASVWEIMMA  661

                                  730       740       750       760       770       780
Consensus               LLNGGSLVIYPZXAIXDPQXLEXYIEEXGVTSLFLTPXLFNLIVDELPP-----ALXTVK  713
2VSQ.A                  MLNAARLIIADEHTLLDTERLTDLILQENVNVMFATTALFNLLTDAGED-----WMKGLR  709
5T3D.A                  FIAGAKLVMAEPEAHRDPLAMQQFFAEYGVTTTHFVPSMLAAFVASLTPQTARQSCATLK  711
4ZXH.A                  LMYGGVVHVCPREVSQDGIQLVDYLNAKSINVLQATPATWKMLLDSEW------SGNAGL  717
BacA_C3-A3              LSCGAMLVIPTETVRQDPQRLWRYLEEQAITHACLTPAMFHD-GTDL-P-----AIAIKP  710
BacA_C3-GxpS_A3         LSCGAMLVIPTETVRQDPQRLWRYLEEQAITHACLTPAMFHD-GTDL-P-----AIAIKP  710
Xbud FclJ_C6-A6         LLNGGSIVIYPQGAISVLL-LEKIIKESGVESLFLTSSLFNLIVDERPQ-----TLQTVK  738
Xhom FclJ_C6-A6         LLNGGSIVIYPQGAISIPQ-LEKMINDSGVESLFLTASLFNLIVDERPQ-----VLQTVK  736
Xsze FclJ_C6-A6         LLNGGSIVIYPQSAISVLL-LEKIIKDGRVDSLFLTTSLFNLIVDERPQ-----TLQTVK  761
Xsze FclJ_C6-Xbud FclJ_A6  LLNGGSIVIYPQGAISVLL-LEKIIKESGVESLFLTSSLFNLIVDERPQ-----TLQTVK  757
Xsze FclJ_C6-Xhom FclJ_A6  LLNGGSIVIYPQGAISIPQ-LEKMINDSGVESLFLTASLFNLIVDERPQ-----VLQTVK  754
GxpS_C3-A3              LSCGAMLVIPTETVRQDPQRLWRYLEEQAITHACLTPAMFHD-GTDL-P-----AIAIKP  714
XtpS_C3-A3              LLNGGALVVIDRAMLLTPVELVRAVQNHGITVMWLTVGLFNRLSTELSP-----ALPQIK  719
XtpS_C3-GxpS_A3         LSCGAMLVIPTETVRQDPQRLWRYLEEQAITHACLTPAMFHD-GTDL-P-----AIAIKP  714

                                  790       800       810       820       830       840
Consensus               QLIXGGEAMSPAHAXRJCSRYP-XDLINGYGPTENTVFTTXYCYRA-----TEGVSXPIG  767
2VSQ.A                  CILFGGERASVPHVRKALRIMGPGKLINCYGPTEGTVFATAHVVHDL---PDSISSLPIG  766
5T3D.A                  QVFCSGEAL-PADLCREWQQLTGAPLHNLYGPTEAAVDVSWYPAFGEELAQVRGSSVPIG  770
4ZXH.A                  TALCGGEALDTILAEKLLGKV--GCLWNVYGPTETTVWSSAARI-------TDAKYIDLG  768
BacA_C3-A3              TLIFAGEAPSPALFQALCSR---ADLFNAYGPTEITVCATTWDCPAD---YTGGV-IPIG  763
BacA_C3-GxpS_A3         TLIFAGEAPSPALFQALCSR---ADLFNAYGPTEITVCATTWDCPAD---YTGGV-IPIG  763
Xbud FclJ_C6-A6         QLISGGEAMSSWHAARIKERYPELLLINGYGPTENTTFTTSHCYRA-----TEGNSVPIG  793
Xhom FclJ_C6-A6         QLISGGEAMSAAHAARISKHYPQLRLINGYGPTENTTFTTSYCYRG-----DEGTSVPIG  791
Xsze FclJ_C6-A6         QLISGGEAMSSAHANRICKYYPELHLINGYGPTENTTFTTSYRYRG-----TEGNSVPIG  816
Xsze FclJ_C6-Xbud FclJ_A6  QLISGGEAMSSWHAARIKERYPELLLINGYGPTENTTFTTSHCYRA-----TEGNSVPIG  812
Xsze FclJ_C6-Xhom FclJ_A6  QLISGGEAMSAAHAARISKHYPQLRLINGYGPTENTTFTTSYCYRG-----DEGTSVPIG  809
GxpS_C3-A3              TLIFAGEAPSPALFQALCSR---ADLFNAYGPTEITVCATTWDCPAD---YTGGV-IPIG  767
XtpS_C3-A3              ILIVGGDVLDPQVISQVLNTNPPQQLLNGYGPSEGTTFTTTYCIRAL---AQSATNIPIG  776
XtpS_C3-GxpS_A3         TLIFAGEAPSPALFQALCSR---ADLFNAYGPTEITVCATTWDCPAD---YTGGV-IPIG  767
                                                                          Chain VIII

                                  850       860       870       880       890       900
Consensus               KPXANNXLYILDEXRQPVPJGTVGELYIGGDGLARGYLNRPELTAERFIEBPFSDSLSAR  827
2VSQ.A                  KPISNASVYILNEQSQLQPFGAVGELCISGMGVSKGYVNRADLTKEKFIENPF--KPGET  824
5T3D.A                  YPVWNTGLRILDAMMHPVPPGVAGDLYLTGIQLAQGYLGRPDLTASRFIADPF--APGER  828
4ZXH.A                  EPLANTQLYVLDEQQRLVPPGVMGELWIGGDGLAVDYWQRPELTDAQFRTLP-SLPNAGR  827
BacA_C3-A3              SPVANKRLYLLDEHRQPVPLGTVGELYIGGVGVARGYLNRPELTAERFLNDPFSDETNAR  823
BacA_C3-GxpS_A3         SPVANKRLYLLDEHRQPVPLGTVGELYIGGVGVARGYLNRPELTAERFLNDPFSDETNAR  823
Xbud FclJ_C6-A6         KPNEGNLVYILDTFHQPVPIGTAGELYIGGDGLAMTYLNQESLYQDLFIENPFDKSLSAR  853
Xhom FclJ_C6-A6         KPNAGNLAYILDAFRQPVPIGTVGELYIGGDGLAITYLNQEQLYQKLFIENPFDSSLSAR  851
Xsze FclJ_C6-A6         KPNEGNLVYILNAFRQPVPIGTTGELYIGGKGLALAYLNQDSLYQNLFIENPFDKSLSTR  876
Xsze FclJ_C6-Xbud FclJ_A6  KPNEGNLVYILDTFHQPVPIGTAGELYIGGDGLAMTYLNQESLYQDLFIENPFDKSLSAR  872
Xsze FclJ_C6-Xhom FclJ_A6  KPNAGNLAYILDAFRQPVPIGTVGELYIGGDGLAITYLNQEQLYQKLFIENPFDSSLSAR  869
GxpS_C3-A3              SPVANKRLYLLDEHRQPVPLGTVGELYIGGVGVARGYLNRPELTAERFLNDPFSDETNAR  827
XtpS_C3-A3              RPIANTRVYLLDNHGQPVPLGAIGEIYIGGDGVACGYLNRPELTADRFLIDPFSDVPDAR  836
XtpS_C3-GxpS_A3         SPVANKRLYLLDEHRQPVPLGTVGELYIGGVGVARGYLNRPELTAERFLNDPFSDETNAR  827
                                   Chain IX                               Chain X
```

```
                                      910       920       930       940       950       960
Consensus                   LYRTGDLARYLPDGNJEYLGRIDQQVKIRGFRIETGEIEAVLCEHPXVEXAVVLX-J---  883
2VSQ.A                      LYRTGDLARWLPDGTIEYAGRIDDQVKIRGHRIELEEIEKQLQEYPGVKDAVVVADR---  881
5T3D.A                      MYRTGDVARWLDNGAVEYLGRSDDQLKIRGQRIELGEIDRVMQALPDVEQAVTHACVINQ  888
4ZXH.A                      LYRTGDKVCLRTDGRLTHHGRLDFQVKIRGFRIELGEIENVLKQIDGITDAVVLVKT---  884
BacA_C3-A3                  MYRAGDLARYLPDGNLVFVGRNDQQVKIRGFRIEPGEIEARLVEHSEVSEALVLA-L---  879
BacA_C3-GxpS_A3             MYRAGDLARYLPDGNLVFVGRNDQQVKIRGFRIEPGEIEARLVEHSEVSEALVLA-L---  879
Xbud FclJ_C6-A6             LYKTGDRGRYLPNGDIEYLGRIDNQNKIRGYRVETGEIEAVLCKHPDVERAVVRV-I---  909
Xhom FclJ_C6-A6             LYKTGDRGRYLPDGEIEYLGRIDSQNKILGYRIETKEIEAVLCQHPIVERAVVRV-I---  907
Xsze FclJ_C6-A6             LYKTGDRGRYLPNGDIEYLGRIDNQNKIRGYRIETGEIEAVLCKHPDIERAAVRI-I---  932
Xsze FclJ_C6-Xbud FclJ_A6   LYKTGDRGRYLPNGDIEYLGRIDNQNKIRGYRVETGEIEAVLCKHPDVERAVVRV-I---  928
Xsze FclJ_C6-Xhom FclJ_A6   LYKTGDRGRYLPDGEIEYLGRIDSQNKILGYRIETKEIEAVLCQHPIVERAVVRV-I---  925
GxpS_C3-A3                  MYRAGDLARYLPDGNLVFVGRNDQQVKIRGFRIEPGEIEARLVEHSEVSEALVLA-L---  883
XtpS_C3-A3                  LYRTGDLARYLPDGNLEFLGRNDQQVKIRGFRIELGEIEARLAEYPAVREATVLV-L---  892
XtpS_C3-GxpS_A3             MYRAGDLARYLPDGNLVFVGRNDQQVKIRGFRIEPGEIEARLVEHSEVSEALVLA-L---  883
                                      Chain XI       Chain XII  Chain XIII

                                      970       980       990      1000      1010
Consensus                   ---GEGXDKRLVAYVVARAXQGLXAMKLRSHLSERLPXYMIPAAFVRLDXLPLTPNG     937
2VSQ.A                      ---HESGDASINAYLVNRTQ--LSAEDVKAHLKKQLPAYMVPQTFTFLDELPLTTNG     933
5T3D.A                      AAATGGDARQLVGYLVSQSGLPLDTSALQAQLRETLPPHMVPVVLLQLPQLPLSANG     945
4ZXH.A                      ---TGDNDQKLVAYV---TGQELDIAGLKKNLQIHLPAYMVPSAFIRLDEFPMTANK     935
BacA_C3-A3                  ---GDGQDKRLVAYVVALADDGL-ATKLREHLSDILPDYMIPAAFVRLDAFPLTPNG     932
BacA_C3-GxpS_A3             ---GDGQDKRLVAYVVALADDGL-ATKLREHLSDILPDYMIPAAFVRLDAFPLTPNG     932
Xbud FclJ_C6-A6             ---EENRGKRIAAYVVLRAGQTLNTMELRSYLAERLPRYMIPAYFRALSSLPLNDNG     963
Xhom FclJ_C6-A6             ---EEARGKRIAAYVVLRAQQILDVMALRSYLSERLPRYMIPTYFQALSSLPLKENG     961
Xsze FclJ_C6-A6             ---KETRGKRIASYIVPRAQKTLDMMELRIYLAERLPRYMLPTYFQTLPSLPLNANG     986
Xsze FclJ_C6-Xbud FclJ_A6   ---EENRGKRIAAYVVLRAGQTLNTMELRSYLAERLPRYMIPAYFRALSSLPLNDNG     982
Xsze FclJ_C6-Xhom FclJ_A6   ---EEARGKRIAAYVVLRAQQILDVMALRSYLSERLPRYMIPTYFQALSSLPLKENG     979
GxpS_C3-A3                  ---GDGQDKRLVAYVVALADDGL-ATKLREHLSDILPDYMIPAAFVRLDAFPLTPNG     936
XtpS_C3-A3                  ---GDGQDKRLVAYIVADVNEEL-VNNLRSHLSKVLPDYMVPAAFMRLDAFPLTPNG     945
XtpS_C3-GxpS_A3             ---GDGQDKRLVAYVVALADDGL-ATKLREHLSDILPDYMIPAAFVRLDAFPLTPNG     936
                             Chain XIV                      Chain XV          Chain XVI
```

### Supplementary Figures

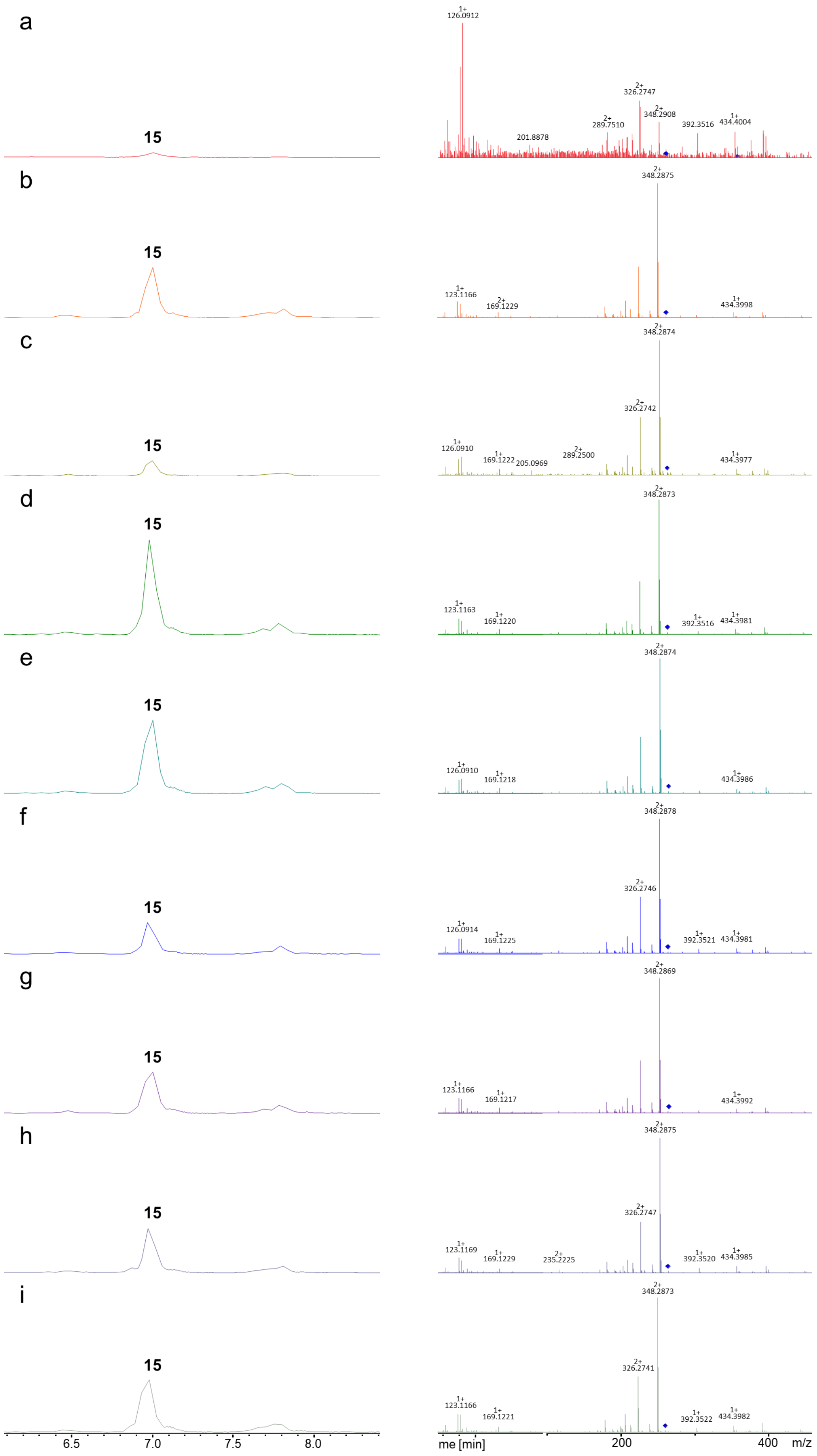

a
15
1+ 126.0912
2+ 326.2747
2+ 348.2908
2+ 289.7510
201.8878
392.3516
1+ 434.4004
b
15
2+ 348.2875
1+ 123.1166
2+ 169.1229
1+ 434.3998
c
15
2+ 348.2874
2+ 326.2742
1+ 126.0910
1+ 169.1222
205.0969
2+ 289.2500
1+ 434.3977
d
15
2+ 348.2873
1+ 123.1163
1+ 169.1220
1+ 392.3516
1+ 434.3981
e
15
2+ 348.2874
1+ 126.0910
1+ 169.1218
1+ 434.3986
f
15
2+ 348.2878
2+ 326.2746
1+ 126.0914
1+ 169.1225
1+ 392.3521
1+ 434.3981
g
15
2+ 348.2869
1+ 123.1166
1+ 169.1217
1+ 434.3992
h
15
2+ 348.2875
2+ 326.2747
1+ 123.1169
1+ 169.1229
2+ 235.2225
1+ 392.3520
1+ 434.3985
i
15
2+ 348.2873
2+ 326.2741
1+ 123.1166
1+ 169.1221
1+ 392.3522
1+ 434.3982
6.5
7.0
7.5
8.0
me [min]
200
400
m/z

**Figure S1.** HPLC/MS data refers to Figure 4 (Plasmid based FclJ complementation: NRPS-9 to -17) of compound **15** produced in *X. szentirmaii ΔfclJ*.

(a) Extracted ion chromatogram (EIC)/$MS^2$ of **15** (m/z $[M+H]^{2+}$ = 357.32; NRPS-9).
(b) Extracted ion chromatogram (EIC)/$MS^2$ of **15** (m/z $[M+H]^{2+}$ = 357.32; NRPS-10).
(c) Extracted ion chromatogram (EIC)/$MS^2$ of **15** (m/z $[M+H]^{2+}$ = 357.32; NRPS-11).
(d) Extracted ion chromatogram (EIC)/$MS^2$ of **15** (m/z $[M+H]^{2+}$ = 357.32; NRPS-12).
(e) Extracted ion chromatogram (EIC)/$MS^2$ of **15** (m/z $[M+H]^{2+}$ = 357.32; NRPS-13).
(f) Extracted ion chromatogram (EIC)/$MS^2$ of **15** (m/z $[M+H]^{2+}$ = 357.32; NRPS-14).
(g) Extracted ion chromatogram (EIC)/$MS^2$ of **15** (m/z $[M+H]^{2+}$ = 357.32; NRPS-15).
(h) Extracted ion chromatogram (EIC)/$MS^2$ of **15** (m/z $[M+H]^{2+}$ = 357.32; NRPS-16).
(i) Extracted ion chromatogram (EIC)/$MS^2$ of **15** (m/z $[M+H]^{2+}$ = 357.32; NRPS-17).

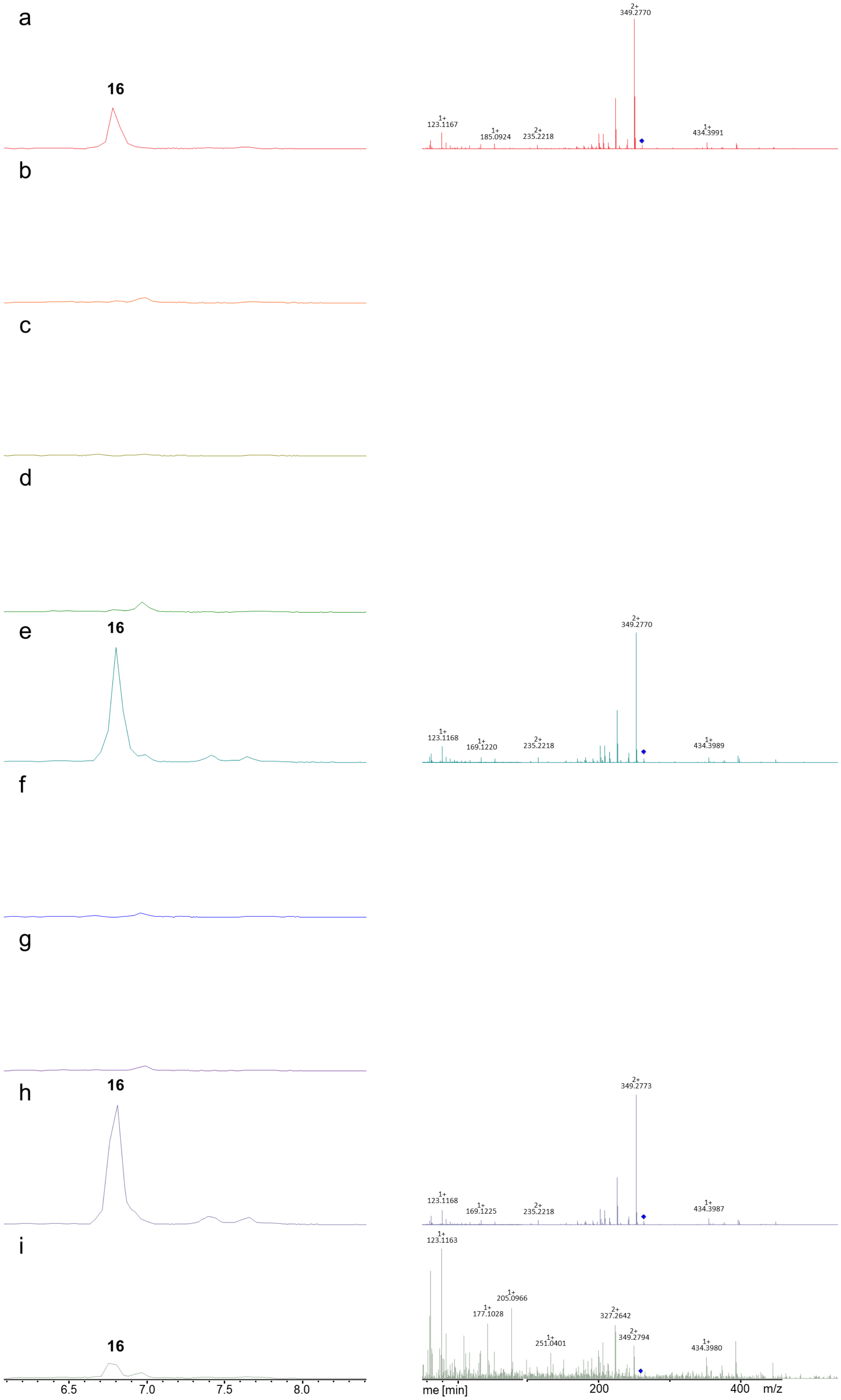

a
16
2+
349.2770
1+
123.1167
1+
185.0924
2+
235.2218
1+
434.3991
b
c
d
2+
349.2770
e
16
1+
123.1168
1+
169.1220
2+
235.2218
1+
434.3989
f
g
2+
349.2773
h
16
1+
123.1168
1+
169.1225
2+
235.2218
1+
434.3987
1+
123.1163
i
1+
205.0966
1+
177.1028
2+
327.2642
1+
251.0401
2+
349.2794
1+
434.3980
16
6.5
7.0
7.5
8.0
me [min]
200
400
m/z

**Figure S2.** HPLC/MS data refers to Figure 4 (Plasmid based FclJ complementation: NRPS-9 to -17) of compound **16** produced in *X. szentirmaii ΔfclJ*.

(a) Extracted ion chromatogram (EIC)/$MS^2$ of **16** (m/z $[M+H]^{2+}$ = 358.32; NRPS-9).
(b) Extracted ion chromatogram (EIC)/$MS^2$ of **16** (m/z $[M+H]^{2+}$ = 358.32; NRPS-10).
(c) Extracted ion chromatogram (EIC)/$MS^2$ of **16** (m/z $[M+H]^{2+}$ = 358.32; NRPS-11).
(d) Extracted ion chromatogram (EIC)/$MS^2$ of **16** (m/z $[M+H]^{2+}$ = 358.32; NRPS-12).
(e) Extracted ion chromatogram (EIC)/$MS^2$ of **16** (m/z $[M+H]^{2+}$ = 358.32; NRPS-13).
(f) Extracted ion chromatogram (EIC)/$MS^2$ of **16** (m/z $[M+H]^{2+}$ = 358.32; NRPS-14).
(g) Extracted ion chromatogram (EIC)/$MS^2$ of **16** (m/z $[M+H]^{2+}$ = 358.32; NRPS-15).
(h) Extracted ion chromatogram (EIC)/$MS^2$ of **16** (m/z $[M+H]^{2+}$ = 358.32; NRPS-16).
(i) Extracted ion chromatogram (EIC)/$MS^2$ of **16** (m/z $[M+H]^{2+}$ = 358.32; NRPS-17).

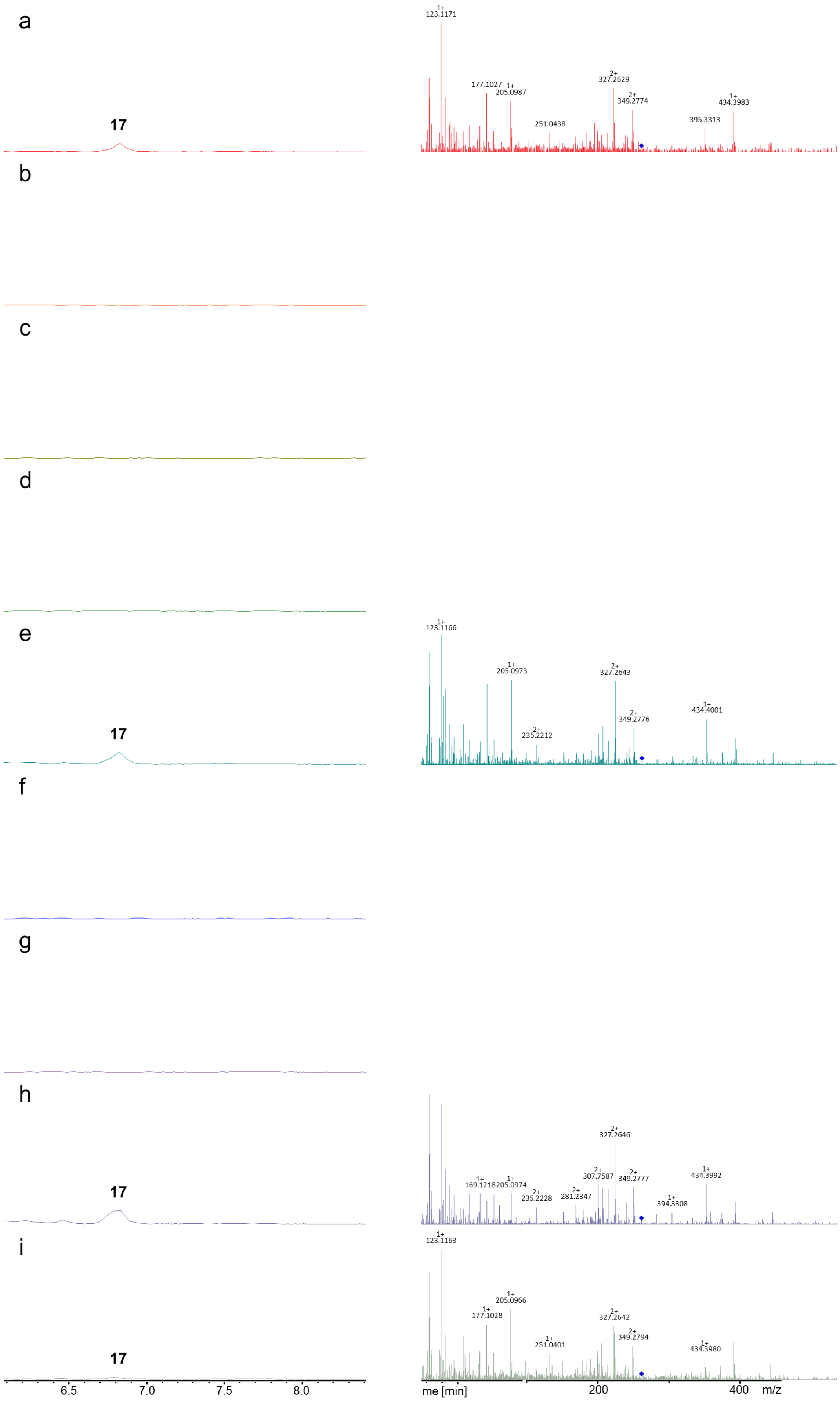

a
17
1+
123.1171
177.1027
1+
205.0987
251.0438
2+
327.2629
2+
349.2774
395.3313
1+
434.3983
b
c
d
e
17
1+
123.1166
1+
205.0973
2+
235.2212
2+
327.2643
2+
349.2776
1+
434.4001
f
g
h
17
1+
169.1218
1+
205.0974
2+
235.2228
2+
281.2347
2+
307.7587
2+
327.2646
2+
349.2777
1+
394.3308
1+
434.3992
i
17
1+
123.1163
1+
177.1028
1+
205.0966
1+
251.0401
2+
327.2642
2+
349.2794
1+
434.3980
6.5
7.0
7.5
8.0
me [min]
200
400
m/z

**Figure S3.** HPLC/MS data refers to Figure 4 (Plasmid based FclJ complementation: NRPS-9 to -17) of compound **17** produced in *X. szentirmaii ΔfclJ*.

(a) Extracted ion chromatogram (EIC)/$MS^2$ of **17** (m/z $[M+H]^{2+}$ = 359.31; NRPS-9).
(b) Extracted ion chromatogram (EIC)/$MS^2$ of **17** (m/z $[M+H]^{2+}$ = 359.31; NRPS-10).
(c) Extracted ion chromatogram (EIC)/$MS^2$ of **17** (m/z $[M+H]^{2+}$ = 359.31; NRPS-11).
(d) Extracted ion chromatogram (EIC)/$MS^2$ of **17** (m/z $[M+H]^{2+}$ = 359.31; NRPS-12).
(e) Extracted ion chromatogram (EIC)/$MS^2$ of **17** (m/z $[M+H]^{2+}$ = 359.31; NRPS-13).
(f) Extracted ion chromatogram (EIC)/$MS^2$ of **17** (m/z $[M+H]^{2+}$ = 359.31; NRPS-14).
(g) Extracted ion chromatogram (EIC)/$MS^2$ of **17** (m/z $[M+H]^{2+}$ = 359.31; NRPS-15).
(h) Extracted ion chromatogram (EIC)/$MS^2$ of **17** (m/z $[M+H]^{2+}$ = 359.31; NRPS-16).
(i) Extracted ion chromatogram (EIC)/$MS^2$ of **17** (m/z $[M+H]^{2+}$ = 359.31; NRPS-17).

| | 2vsq | 2vsq | 2vsq | 2vsq | 2vsq | 4zxh | 4zxh | 4zxh | 4zxh | 4zxh | 5t3d | 5t3d | 5t3d | 5t3d | 5t3d | 2vsq | 2vsq | 2vsq | 2vsq | 2vsq | 4zxh | 4zxh | 4zxh | 4zxh | 4zxh | 5t3d | 5t3d | 5t3d | 5t3d | 5t3d |
|---|---|---|---|---|---|---|---|---|---|---|---|---|---|---|---|---|---|---|---|---|---|---|---|---|---|---|---|---|---|---|
| | gxps3-A3 | xtpsc3-a3 | xtpsc3_gxpsa3 | bacAC3-A3 | bacac3_gxpsa3 | gxpsC3-A3 | xtpsc3-a3 | xtpsc3_gxpsa3 | bacAC3-A3 | bacac3_gxpsa3 | gxpsC3-A3 | xtpsc3-a3 | xtpsc3_gxpsa3 | bacAC3-A3 | bacac3_gxpsa3 | FclJX.bud.C6-A6 | FclJX.hom.C6-A6 | fcljx.sze.c6-a6 | FclJX.szeC6_FclJX.bud.A6 | FclJX.szeC6_FclJX.hom.A6 | FclJX.bud.C6-A6 | FclJX.hom.C6-A6 | Fcljx.sze.c6-a6 | FclJX.szeC6_FclJX.bud.A6 | FclJX.szeC6_FclJX.hom.A6 | FclJX.bud.C6-A6 | FclJX.hom.C6-A6 | fcljx.sze.c6-a6 | FclJX.szeC6_FclJX.bud.A6 | FclJX.szeC6_FclJX.hom.A6 |
| 196 | -1,4880 | | | | | | | | | | | | | | | | | | | | | | | | | | | | | |
| 198 | | | | | | | | | | | | | | | | | | | | | | | | | | -1,3520 | -1,3060 | -1,5470 | -1,2770 | -1,2050 |
| 199 | | | | | | | | | | | | -0,1360 | | | | | | | | | | | | | | | | | | |
| 201 | x | x | x | x | x | x | x | x | x | x | x | x | x | x | x | | | -1,5830 | | | | | | | | -1,3010 | -1,2180 | -1,4620 | -1,2300 | -1,3010 |
| 202 | | | | | | | | | | | -1,3100 | -1,2280 | -1,3670 | -1,5140 | -1,4030 | -0,8490 | -1,6700 | -1,1640 | -1,3100 | -1,3060 | | | | | | -1,3800 | -1,2830 | -1,3140 | -1,0920 | -1,2770 |
| 203 | | | | | | | | | | | | | | | | -1,2620 | | -1,3570 | -1,3660 | -1,4570 | | | | | | | -1,4590 | -1,3530 | -1,4010 | -1,5150 |
| 204 | | | | 0,0330 | -0,1420 | | | | | | | | | 0,5410 | 0,8460 | | -1,2650 | -0,8390 | -1,3910 | | | | | | | -1,0640 | -0,9400 | -1,1180 | -1,1360 | -1,4020 |
| 205 | -0,5070 | -1,1400 | -0,9740 | -0,4590 | -0,3860 | | | | | | | | | | | -0,0820 | -0,5740 | 0,9730 | -1,0830 | -1,1000 | | | | | | | | -0,3100 | -0,6610 | |
| 206 | | | | | | | | | | | | | | | | | | -0,1480 | | | | -1,3330 | | -1,2330 | | | | | | |
| 208 | -0,8830 | -0,5880 | -0,7560 | -0,7750 | -1,0030 | -0,8560 | -1,0520 | -1,0270 | -0,6900 | -0,4730 | | | | | | 2,1650 | 1,8060 | 1,8160 | 1,0860 | 1,1390 | 0,9610 | 0,8250 | 0,2810 | 2,0060 | 0,8800 | 1,2750 | 2,4410 | 1,9620 | 1,8600 | 2,0380 |
| 209 | -0,7820 | -0,6970 | -0,7050 | -0,4370 | -0,5970 | -0,7160 | -0,5520 | -0,5180 | 0,0670 | 0,1430 | | -1,3220 | -1,2270 | | | 0,0360 | 0,1550 | 0,2920 | -0,6380 | -0,3910 | -0,3620 | -0,3270 | -0,2610 | 0,1510 | -0,3740 | | | -0,8910 | | -1,2450 |
| 210 | -0,9300 | -0,8330 | -0,9010 | -0,6800 | -0,9220 | | | | -0,5080 | -0,6100 | -1,0830 | -1,0760 | -0,8750 | -1,0110 | -0,8750 | -0,1020 | 0,1240 | -0,7290 | -0,7140 | -0,5890 | | | -1,0490 | | | -1,3060 | -0,9660 | -0,8530 | -1,1660 | -1,2790 |
| 211 | | | | | | -0,4200 | -0,3520 | -0,2900 | 0,4520 | 0,4140 | -1,2200 | -0,9710 | 0,4590 | -0,5260 | -0,7940 | | | | | | -0,4040 | -0,2000 | -0,9800 | -0,3980 | -0,7400 | | -0,4950 | | | |
| 212 | | | | | | -1,1870 | | | | | -0,9350 | -1,2630 | -0,7490 | -0,7930 | -1,1060 | -1,2780 | | | | | -1,2210 | | | | | -1,4610 | -0,9470 | | | |
| 213 | | | | | | | | | | | | | | | | -0,3790 | 1,0620 | | | | 0,2460 | 0,4550 | | | | | | | | |
| 214 | -0,6940 | -0,8720 | -1,0010 | 0,6860 | 0,4850 | -0,9430 | -0,5600 | -0,7480 | 0,4970 | 0,4730 | | | | | | | -0,4270 | -0,9760 | -0,8480 | -0,5020 | | -0,3370 | -0,8540 | -1,0610 | -1,0460 | | | -1,3960 | | |
| 215 | -0,3170 | -0,1220 | -0,1100 | -0,2430 | -0,2620 | -1,0020 | -0,5460 | -0,9720 | 0,3690 | 0,3420 | | | | | | -1,0260 | -0,8930 | -1,0180 | -0,9040 | -0,7600 | -1,0600 | -0,9360 | | | | | | | | |
| 216 | | | | | | | 1,5210 | 1,0260 | -0,7910 | -0,7950 | | | | | | | -0,6620 | | | | -1,2670 | -0,9960 | | | | | | | | |
| 217 | | | | | | | | | 0,5660 | 0,4980 | | | | | | | | | | | -0,8840 | -1,1020 | -1,0430 | -1,0850 | -1,0120 | | | | | |
| 218 | | | | | | 0,9410 | 1,5840 | 1,4500 | 2,2300 | 2,3120 | | | | | | | | | | | -0,8390 | -0,4540 | | | | | | | | |
| 219 | | | | | | | | | -0,6180 | -0,5950 | | | | | | | | | | | | | | | | | | | | |
| 220 | | | | | | -0,7830 | -0,8090 | -0,8500 | -0,4910 | -0,5350 | | | | | | | | | | | | -0,4580 | -0,6100 | 0,0340 | -0,4140 | | | | | |
| 221 | | | | | | | | | | | | | | | | | | | | | -1,1480 | | 0,8750 | 1,2850 | 0,7650 | | | | | |
| 224 | x | x | x | x | x | x | x | x | x | x | x | x | x | x | x | x | x | | | | x | x | -0,3460 | 0,3860 | -0,2620 | x | x | | | |
| 228 | x | x | x | x | x | x | x | x | x | x | x | x | x | x | x | x | x | | | -0,5860 | x | x | | | | x | x | | | |
| 230 | x | x | x | x | x | x | x | x | x | x | x | x | x | x | x | x | x | | | -0,1380 | x | x | | | -1,2170 | x | x | -1,2380 | | -1,3530 |
| 231 | x | x | x | x | x | x | x | x | x | x | x | x | x | x | x | x | x | | -1,1260 | -0,9770 | x | x | | | -1,5820 | x | x | -0,8910 | -1,3290 | -1,1880 |
| 233 | x | x | x | x | x | x | x | x | x | x | x | x | x | x | x | x | x | | | | x | x | | | | x | x | -1,1440 | | |
| 234 | x | x | x | x | x | x | x | x | x | x | x | x | x | x | x | x | x | | | | x | x | | | | x | x | -0,1810 | | -0,8450 |
| 235 | x | x | x | x | x | x | x | x | x | x | x | x | x | x | x | x | x | | -1,0600 | -0,7560 | x | x | -1,2960 | -1,2970 | | x | x | -0,1790 | -0,6900 | -0,6650 |
| 236 | x | x | x | x | x | x | x | x | x | x | x | x | x | x | x | x | x | | | | x | x | -0,6480 | -1,1910 | -0,8390 | x | x | -0,1250 | -1,0000 | -0,9580 |
| 237 | x | x | x | x | x | x | x | x | x | x | x | x | x | x | x | x | x | | | | x | x | -1,4080 | | -1,4620 | x | x | | | |
| 282 | | | | | | | | | | | | | | | | | | | | | | | | | | | -0,8500 | -0,2360 | | |
| 283 | | | | | | | | | | | | | | | | | | | | | | | | | | | | | | |
| 285 | | | | | -0,9010 | | | | | | | -0,7870 | -0,7480 | | | | | | | | | | | | | | | -0,8590 | | |
| 286 | 1,3230 | 0,6050 | 0,9720 | 0,2640 | 0,6510 | 1,6020 | 1,0710 | 1,0270 | -1,0480 | -1,0550 | 2,1720 | 1,8810 | 1,6890 | 0,4520 | 0,5030 | -0,2790 | -0,3890 | -0,5610 | -0,3990 | -0,3360 | -1,1410 | -1,2840 | -1,0200 | -1,1780 | -1,2490 | -0,1570 | -0,0780 | 0,4370 | -0,1020 | -0,2510 |
| 287 | -0,6690 | -0,4600 | -0,7920 | 0,0490 | 0,1890 | -0,8870 | -1,0200 | -0,9430 | -0,4030 | -0,2160 | -0,8630 | -1,0770 | -0,9830 | -0,6200 | -0,2510 | -0,5630 | -0,1260 | -0,8180 | -0,6950 | -0,4460 | -0,5810 | -0,6980 | -0,7860 | -0,6860 | -0,5530 | -0,9440 | -0,5810 | -0,2680 | -0,6790 | -0,8610 |
| 288 | -1,3150 | -1,1910 | -1,2070 | -0,7710 | -0,7840 | -0,9210 | -0,9440 | -0,8420 | -0,7020 | -0,6690 | -1,3800 | -1,3100 | -1,1660 | | -0,7960 | -0,9060 | -0,4570 | -1,1970 | | -0,8830 | -1,0360 | -1,0750 | | | -1,2210 | -0,9660 | -1,0190 | -0,4000 | -0,9060 | -1,1380 |
| 289 | | | | -0,2030 | -0,3900 | | | | -0,7640 | -0,8310 | | | | -0,6300 | 0,0100 | | | | | | | | | | | | | | | |
| 290 | -0,8210 | -0,5260 | -0,7450 | -0,0070 | -0,6990 | -0,4760 | -0,4480 | -0,3550 | -0,5220 | -0,5600 | -0,3330 | -1,0030 | -0,8810 | -1,4510 | -0,6780 | 1,1760 | 1,0000 | 1,6790 | -0,1890 | 0,4180 | 0,3940 | -0,8860 | -1,4810 | 0,1990 | -0,6690 | -0,2050 | 0,5450 | 1,4630 | -0,2770 | 0,1300 |
| 291 | | | | | -1,5060 | | | | | | 0,6460 | -0,4490 | -0,3550 | -1,5190 | -0,9310 | -0,5930 | | | -1,1620 | | | | | | | -0,5410 | -0,6810 | -0,3200 | -0,6580 | -0,3430 |
| 292 | | | | -0,2530 | -0,8140 | | | | | | | | | | | 0,3120 | -0,3490 | 0,2340 | | | | -0,8830 | | | | | | | | |
| 293 | | | | | | | | | | | | | | | | | | | | | | | | | | | | | | |
| 323 | | | | 2,2280 | 2,0460 | | | | | | | | | | | 2,4420 | 2,1740 | 2,3980 | | | | | | | | | | | | |
| 326 | | | | | | | | | | | -0,3450 | -0,3170 | -0,8730 | | | | | | | | | | | | | -1,0790 | -1,1130 | -0,7280 | -1,1280 | |
| 327 | | | | | | | | | | | -1,5220 | -1,4650 | -1,5540 | | | | | | | | | | | | | | | -1,7620 | | |
| 328 | | | | -1,5300 | | | | | | | -0,5900 | | | | | | | | | | | | | | | | | | | |
| 329 | -1,5120 | -1,6000 | -1,5480 | -1,3420 | -1,4940 | | | | | | | | | | | -1,5300 | | | | | | | | | | | | | | |
| 360 | | | | | | | | | | | | | | | | | | | | | | | | | | -1,3130 | -1,2330 | | -1,3610 | -1,1400 |
| 361 | | | | | | | | | | | -1,5260 | -1,3130 | -1,5170 | -1,5400 | -1,3430 | | | | | | | | | | | -0,7530 | -0,3040 | -0,8150 | -0,6440 | -0,6030 |
| 362 | -1,1670 | -1,1200 | | -0,8320 | -1,2220 | | | | | | -1,1090 | -1,0190 | -1,1090 | -1,7150 | -1,3020 | | | | | | | | | | | | | | | |
| 363 | | | | | | | | | | | | | | | | -0,5270 | -0,9040 | -0,7350 | -0,7460 | -0,9740 | | | | | | | -1,1830 | | | -1,3970 |
| 364 | | | | | | | | | -1,1610 | -0,8400 | | | | | | -0,5840 | -0,9400 | -0,9780 | -0,8940 | -1,0030 | -0,7500 | | -0,2230 | -0,5440 | -0,8680 | | | | | |
| 365 | | 0,4660 | | -0,4690 | -0,3090 | | 0,3420 | 0,4050 | | | | | | | | | | | | | | | | | | | | | | |
| 366 | | | | | | 0,3290 | 0,1100 | -0,0900 | -0,6660 | 0,6010 | | | | | | | | | | | -0,5100 | -0,5440 | -0,1290 | -0,0060 | -0,1310 | | | | | |
| 367 | | | | | | | | | | | | | | | | | | | | | -0,8990 | | | | | | | | | |
| 368 | | | | | | -0,5740 | 0,2430 | -0,2520 | 0,0040 | 0,0380 | | | | | | | | | | | -0,3880 | 0,1370 | 0,1130 | 0,7620 | 0,1600 | | | | | |
| 369 | | | | | | | -0,5920 | -0,6170 | | | | | | | | | | | | | | | | | -0,6350 | | | | | |
| 370 | | | | | | | | | | | | | | -1,6120 | | | | | | | | | | | | | | | | |
| 444 | | | | | | | | | | | | | | | | | | | | | -0,5880 | -0,7980 | | | | | | | | |
| 446 | | | | -1,1130 | | | | | -0,4220 | -0,3790 | | | | | | | | | | | | | | | | | | | | |
| 447 | | | | | | | | | | | | | | | | | | | | | | | | | | | | | | |
| 448 | | | | -0,7220 | | -1,4720 | -1,0610 | -1,4940 | -1,5200 | -1,4950 | | | | | | | | -1,1700 | | -1,2410 | | | -0,5080 | -0,4850 | -0,6750 | | | | | |
| 449 | | | | | | -1,0400 | -0,7930 | -1,0500 | -0,3990 | -0,3470 | | | | | | | -0,8240 | -0,7930 | -1,0080 | -0,6560 | -0,8580 | -0,6930 | -0,6510 | -0,6520 | -0,8190 | -1,5940 | -0,9490 | -1,3020 | -1,2820 | -1,4650 |
| 451 | | | | | | | | | | | | | | | | | | | | | | | | | | | | | | |
| 452 | 1,7760 | 2,0920 | 2,1600 | -0,1990 | -0,5680 | 1,6800 | 1,1240 | 0,8180 | -0,7610 | -0,6980 | 1,4550 | | | | -0,7330 | -1,3990 | -0,8220 | -0,2960 | -0,3540 | -0,1190 | -0,7140 | -0,6990 | -0,2350 | -0,1610 | -0,3710 | | | -0,9320 | -0,9010 | -1,2050 |
| 453 | | | | | | | | | | -0,7630 | -0,9630 | | | | -0,9400 | | | | | | | | | | | -1,5090 | -1,2840 | -1,1760 | -1,1170 | -1,2530 |
| 455 | | | | | | | | | | | | | | | | | | | | | | | | | | | | | | |
| 456 | -0,1800 | -0,5630 | -0,2820 | -0,7320 | -0,6880 | -0,2610 | -0,3910 | -0,3900 | -0,5890 | -0,3820 | -0,5560 | -0,9550 | -0,8320 | | -0,6010 | -1,5420 | -1,2280 | -1,3870 | -1,3130 | -1,2960 | | | | | | -1,6860 | -1,5970 | -1,4490 | -1,4710 | -1,4830 |
| 467 | | | | | | | | | | | | | | | | | | -0,5620 | | | | | | | | | | | | |
| 471 | | | | | | | | | | | | | | | | | | | | | | | | | | | | -1,6960 | | |
| 472 | | | | | | | | | | | -0,2020 | -0,8950 | -1,1790 | | | | | | | | | | | | | -1,1400 | -0,9610 | -0,9860 | -1,2860 | |
| 473 | -0,7180 | | | -0,1240 | -0,2980 | -1,2390 | | | -1,0880 | -1,1510 | -0,2010 | | | -0,5610 | -0,5230 | -0,7470 | -0,9040 | -1,0870 | -1,1590 | -0,8440 | -1,5070 | -1,4270 | -1,2410 | -1,3750 | | -0,8970 | -0,8140 | -0,7990 | -1,0550 | -0,6640 |
| 474 | | -1,0760 | -1,0300 | 0,0930 | -0,4770 | | -1,2190 | | -1,4150 | -1,4580 | | | | -0,6640 | -0,8540 | -1,5450 | -1,4800 | -1,2480 | -1,5130 | -1,4210 | | | | | | -1,3410 | -1,2480 | -1,2210 | -1,4740 | -1,3780 |
| 475 | -0,7680 | -0,9240 | -0,7880 | -0,5080 | -0,6790 | | | | | | -0,3110 | -0,7050 | -0,8320 | -0,7250 | -0,8620 | | | | | | | | | | | | | | | |
| 476 | -0,4300 | -0,5970 | -0,4240 | -0,4610 | -0,4530 | -0,1540 | -0,3710 | -0,4210 | | -0,9760 | -0,1230 | -0,3560 | -0,6350 | -0,7460 | -0,5570 | -1,4510 | -1,3960 | -0,7900 | -1,0480 | -0,9490 | -1,3390 | -1,4700 | -0,7060 | | -1,4670 | -0,5280 | -0,7330 | -0,4160 | -0,7110 | -0,2310 |
| 477 | -0,4950 | -0,6560 | -0,3010 | -0,3610 | -0,5890 | -0,9360 | -1,0730 | -0,9720 | -1,1310 | -1,1250 | -0,1090 | -0,7040 | -0,6390 | -0,5000 | -0,4850 | 0,1920 | 0,1120 | -0,1400 | -0,1550 | 0,0270 | -0,9450 | -1,0750 | -0,9200 | -1,0230 | -1,1670 | -0,5220 | -0,5690 | 0,2040 | -0,3820 | -0,3300 |
| 478 | -0,5230 | -0,6850 | -0,4510 | -0,4820 | -0,1160 | -0,0910 | -0,3010 | -0,3600 | -0,5930 | -0,7200 | -0,5490 | -0,8690 | -0,6730 | -0,5660 | -0,5400 | -0,4720 | -0,2180 | -0,1060 | -0,0270 | -0,0870 | -0,6630 | -0,5530 | -0,4480 | -0,8900 | -0,4390 | -0,5950 | -0,9640 | -0,4130 | -0,6070 | -0,6380 |
| 479 | | | | | | -1,2460 | -1,1940 | -1,0130 | -1,3790 | -1,3410 | | | | | | | | | | | -1,2450 | -1,3710 | -1,1030 | -1,1160 | -1,1430 | | | -1,2730 | | |
| 480 | | | | | -1,3980 | | | | | | | | | | | | | | | | | | | | | | | | | |
| 481 | | | | | | -0,8750 | | -0,4080 | 0,0500 | -0,7730 | | | | | | | | | | | -0,9550 | -0,7390 | -0,9010 | -0,8160 | -0,6200 | | | | | |
| 482 | -0,1680 | -0,4340 | -0,5600 | -0,5870 | -0,6920 | | | | | | -1,0290 | | -1,3120 | -1,0010 | -0,7960 | | -0,8210 | | | | | | | | | -0,9350 | | | | |
| 483 | 1,0350 | 0,6440 | 1,2680 | -1,0330 | -0,5920 | 1,3560 | -0,1770 | 0,2570 | -6,9200 | -0,8620 | 0,1150 | 0,5520 | -0,6680 | -0,7850 | -0,3760 | -0,8480 | -0,8320 | -0,6880 | -0,7350 | -0,9790 | | | | | | -0,8950 | -0,9960 | -0,4870 | -0,9960 | -0,3720 |
| 484 | | | | -0,9190 | | -0,8180 | -0,6200 | -0,3960 | 0,8320 | -0,2810 | | | | | | -1,1880 | -7,3600 | -7,2770 | -7,1090 | -7,5160 | -0,6980 | -0,3690 | -0,8740 | -0,6200 | -0,1670 | | | | | |
| 485 | -6,8610 | -6,8040 | -6,7860 | -6,6600 | -7,1060 | -5,6960 | -5,8630 | -5,9140 | -5,8570 | -6,1500 | -7,1500 | -7,2470 | -7,2690 | -7,0730 | -7,4340 | 1,2430 | 1,0550 | 2,4450 | 2,3330 | 2,4360 | 1,8780 | 1,8300 | 1,7150 | 2,2640 | 1,6090 | 1,5480 | -0,7940 | 1,7010 | -0,0500 | -0,1480 |
| 486 | -6,1690 | -6,0370 | -5,8960 | -6,1960 | -6,3330 | -5,7860 | -5,8870 | -5,8850 | -6,0510 | -6,2470 | -6,5000 | -6,3530 | -6,5800 | -6,4800 | -6,5890 | -5,0490 | -6,3270 | -5,0310 | -4,8800 | -5,0570 | -5,4070 | -6,3330 | -5,1850 | -5,0830 | -5,4420 | -5,5950 | -6,4480 | -5,5250 | -5,5740 | -5,2880 |
| 487 | -6,4390 | -6,1610 | -6,2210 | -6,8470 | -6,9090 | -6,3000 | -6,5730 | -6,5910 | -6,6850 | -6,5980 | -7,0030 | -6,2970 | -7,0190 | -6,9880 | -6,6320 | -6,3460 | -6,6910 | -6,5710 | -6,3230 | -6,6550 | -6,4120 | -6,7710 | -6,5500 | -6,3360 | -6,6350 | -6,8450 | -6,6870 | -6,5030 | -6,6360 | -6,9410 |
| 488 | -1,4880 | -5,7470 | -5,5870 | -0,9030 | -0,9270 | -0,2420 | -6,6230 | -7,1530 | -0,3960 | -1,0530 | -1,1350 | -6,5730 | -7,2990 | -0,6860 | -1,0660 | -0,1510 | -1,1590 | -0,3970 | -1,4920 | -1,5550 | -0,2100 | -1,0230 | -1,2060 | -0,7860 | -1,0320 | -0,9770 | -0,7220 | 0,0550 | -0,1510 | -0,4770 |
| 489 | -7,1260 | -7,0390 | -7,1890 | -7,1470 | -7,4120 | -6,1490 | -6,6020 | -6,9010 | -6,9650 | -6,7590 | -7,4220 | -7,3880 | -7,0720 | -7,4160 | -7,7070 | -7,0280 | -7,5270 | | | | -6,5470 | -6,9480 | | | | -6,9810 | -6,5600 | | | |
| 490 | -5,8850 | -5,6570 | -5,8450 | -5,9040 | -6,1230 | -5,0580 | -5,3300 | -5,4910 | -5,3950 | -5,3350 | -6,1060 | -5,7130 | -6,1250 | -6,0210 | -6,3820 | -6,1370 | -6,7610 | -6,5270 | -6,3790 | -6,2800 | -5,4510 | -5,9470 | -5,7340 | -5,4280 | -5,7070 | -5,9700 | -6,0580 | -5,8380 | -5,8180 | -6,0000 |
| 491 | -6,8290 | -6,1610 | -6,3940 | -6,7640 | -6,9890 | -6,6700 | -6,9180 | -6,8470 | -6,2060 | -6,3970 | -6,6990 | -6,2950 | -6,7690 | -6,7090 | -6,7990 | -6,5750 | -7,2940 | -7,0310 | -6,7650 | -7,1020 | -6,5040 | -6,8940 | -6,7610 | -6,4520 | -7,3390 | -6,7030 | -6,9610 | -6,6860 | -6,4180 | -7,0790 |
| 492 | | | | | | | | | | | | | | | | -7,2460 | -7,5770 | -7,5240 | -7,3590 | -7,4240 | | -6,9930 | -7,1120 | | | -6,8160 | -6,8000 | -6,6810 | -6,5210 | -7,2210 |
| 493 | -1,5300 | -1,3690 | | | | | | | | | -7,2800 | | | | | | | | | | | | | | | | | | | |
| 494 | | -0,6610 | -0,7770 | | | | | | | | | | | | | | | | | -2,0240 | | | | | | | | | | |
| 795 | | | | | | | | | | | | | | | | | | | | | | | -0,9500 | | | | | | | |
| 799 | | | | | | | | | | | | | | | | | | | | | | | -1,2780 | | -1,1380 | | | | | |
| 835 | | | | | | | | | | | | | | -1,2260 | -1,5430 | | | | -1,7640 | | | | -1,4280 | | | | | | | |
| 841 | | -1,2240 | | | | -7,2740 | 0,4600 | -7,4860 | -7,6330 | -7,3200 | | -0,7670 | -7,7070 | | | | | | | | -6,6710 | -6,8370 | -6,9800 | -6,8750 | -7,1820 | | | -7,1110 | -7,1880 | -6,8480 |
| 847 | | 2,5600 | 2,9850 | | | 1,2860 | 2,1900 | 1,2330 | | | | 2,6180 | 2,2190 | | 2,9310 | -1,1760 | | -1,1730 | | | -6,8400 | | | | -6,9180 | | | | | |
| 848 | -1,0090 | -7,4000 | -7,1920 | -7,2070 | -7,3580 | -6,5570 | -6,8060 | -6,7350 | -6,8770 | -6,6880 | | | | | -1,0120 | | | -1,2960 | -1,4320 | | -6,9630 | | -7,0880 | -6,6530 | | | | | -6,8920 | |
| 849 | -0,4160 | -0,1050 | 0,0450 | -0,6840 | -0,7220 | -0,0830 | 0,1110 | 0,0380 | 0,2540 | -0,3340 | | -0,2980 | -0,8320 | | -0,5740 | -0,6290 | -0,4910 | -0,5940 | -0,0770 | -0,3620 | -0,2530 | -0,4040 | -0,4480 | -0,1300 | -0,2410 | | | | | |
| 850 | | -0,9310 | | | | -0,9500 | -1,0110 | -1,0400 | -1,0430 | -1,1710 | | | | | | -0,6180 | | -0,7690 | -0,6460 | -0,7590 | -1,0310 | -1,1870 | -1,0080 | -0,8750 | -0,9370 | | | | -0,8610 | |
| 851 | | | -0,4470 | | | | | | | | | | | -1,0770 | -0,7730 | | -0,9140 | -0,7780 | -0,7890 | -0,5630 | | | | | | -0,5940 | | -0,6940 | -0,8040 | -0,3900 |
| 852 | 0,1030 | -0,2550 | 0,1900 | 0,2700 | 0,4640 | -0,2300 | -0,2270 | -0,6800 | -0,6760 | -0,9680 | 1,2840 | 0,6380 | 0,6470 | 0,2550 | 0,5020 | 0,1490 | -0,2810 | -0,8390 | -0,2970 | 0,0430 | -1,0090 | -1,0110 | -0,8340 | -1,0740 | | 0,2280 | 0,5450 | 0,9790 | 0,5520 | 0,4910 |
| 853 | -1,1880 | | -1,2280 | -0,9820 | -0,8060 | | -1,1970 | | -2,0110 | -1,8080 | 2,3470 | 0,2290 | -0,7040 | -1,1400 | -1,1230 | | | | | | | | | -1,7000 | | -0,5120 | | | -0,3760 | |
| 854 | -0,4310 | -0,7720 | -0,3060 | 0,0190 | -0,3710 | -0,9300 | -0,7230 | | -0,7300 | -1,2100 | 1,7810 | 0,3550 | 0,5600 | 0,0750 | 0,2030 | -0,6890 | -0,7200 | -0,9170 | -0,8590 | -0,7150 | -1,2350 | -1,1740 | -1,0110 | | | -0,0680 | 0,1200 | 0,4150 | -0,3480 | -0,3010 |
| 855 | | | -0,7720 | -0,4300 | -1,0670 | | | | | | | | | | | | | | | | | | | | | | | | | |
| 856 | -0,0670 | -0,1170 | 0,2730 | 0,4760 | 0,2300 | -0,7350 | -0,7710 | -0,5220 | -0,4720 | -1,0430 | 0,1700 | -0,2110 | -0,4820 | -0,0610 | 0,2850 | 0,0550 | -0,0660 | 0,0320 | -0,0770 | 0,1470 | -1,0400 | -1,0620 | -0,7510 | | -0,8690 | 0,3100 | 0,1270 | 0,8720 | -0,0890 | 0,5620 |
| 858 | -0,3910 | -0,6660 | -0,3790 | -0,2860 | -0,0150 | -0,1320 | -0,3760 | -0,3250 | -0,8620 | -0,8990 | -0,2440 | -0,4240 | -0,4470 | -0,5680 | -0,1840 | -0,3630 | -0,4300 | -0,5750 | -0,2330 | -0,1050 | -0,8570 | -0,9180 | -0,6930 | -0,9980 | -0,9680 | -0,4470 | -0,7350 | 0,2760 | -0,3800 | -0,4010 |
| 860 | -6,7290 | -6,4710 | -0,3850 | -0,0630 | -6,7900 | -0,1970 | -0,3250 | -0,4100 | -0,0990 | -0,3380 | -0,7680 | -1,2070 | -1,1090 | -1,0330 | -0,8670 | -6,8700 | -0,7040 | -6,4780 | -6,2590 | 0,1670 | -0,6490 | -0,8350 | -0,4120 | -0,4560 | -0,5690 | -0,8490 | -0,9310 | -0,7920 | -0,6050 | -0,8790 |
| 862 | -0,2930 | | -0,2880 | 0,4470 | 0,6050 | -0,0770 | | -0,3990 | -0,4060 | -0,3240 | -0,2510 | | -0,7090 | -0,5870 | -0,3910 | -0,1680 | -0,1470 | -0,3230 | 0,0070 | 0,3090 | -0,6690 | -0,6980 | -0,7600 | -0,9470 | -0,8340 | -0,8230 | -0,6430 | 0,0140 | -0,5180 | -0,8490 |
| 863 | -1,0530 | -0,7040 | -0,9590 | -0,3800 | -0,3620 | -1,2260 | -1,0240 | -1,1610 | -0,8890 | -0,7880 | -0,9340 | -0,8500 | -0,7480 | -0,8430 | -0,6710 | | -0,4450 | -0,6650 | | -0,2410 | | -0,9410 | -1,4280 | | -1,1480 | | -0,0360 | 0,0880 | | -0,7150 |
| 865 | | | | | | | | | | | | | | -0,5370 | -0,1460 | | | | | | | | | | | | | | | |
| 893 | -0,1610 | -0,8030 | -0,1970 | -0,9000 | -0,9910 | | | | -1,1230 | | -1,1260 | -0,4710 | -1,0540 | -1,2120 | -0,8800 | | -1,3880 | -0,6330 | | -1,0420 | | | | | | -0,7510 | -1,1430 | -1,1750 | | -0,7190 |
| 894 | -1,2170 | -0,8830 | | -1,4770 | -1,2080 | -1,1040 | -1,2560 | -0,9030 | -0,8510 | -1,4690 | | | | | -1,3110 | | -0,5480 | | -1,3250 | | -0,9190 | -0,3710 | -0,6700 | -0,9510 | -0,0760 | | | -0,3820 | | -0,3020 |
| 895 | -0,9160 | -0,8250 | -1,0330 | | -1,2940 | -0,6710 | -0,5160 | -0,2040 | -0,4930 | -0,6800 | | -0,3930 | | | -0,8340 | -0,4030 | -1,3600 | -0,6090 | -0,8370 | -1,6570 | -0,9390 | -1,3060 | -0,7240 | -0,6030 | -1,1370 | | | | | |
| 896 | | -1,3530 | -0,1730 | | -1,1230 | -0,9040 | -1,4180 | -0,6720 | | | | | | | | | | | -1,5060 | | | -1,5210 | | -1,7060 | -1,4040 | | | | | |
| 897 | | | | | | -1,0110 | | -1,0590 | -1,0080 | -1,3610 | | | | | | | | | | | -1,4270 | -1,3050 | -1,4480 | -1,4550 | -1,3050 | | | | | |
| 898 | | | | | | -0,5360 | -0,2670 | -0,3840 | -0,4450 | -0,9920 | | | | | | | | | | | | -1,2320 | | | -1,1590 | | | | | |
| 907 | | | | -0,8450 | | | | | | | | | | | -1,1370 | -1,0500 | -0,5170 | -1,2270 | -1,0010 | -0,7200 | | | | | | 0,5770 | 0,5320 | 0,7060 | 0,9420 | 0,5750 |
| 909 | -0,1000 | 1,1590 | -0,0740 | 0,8100 | 0,6120 | 0,2390 | 1,4450 | 0,1780 | 1,0350 | 0,6870 | 0,1570 | 0,9420 | -0,1210 | 0,1740 | -0,0020 | 0,9110 | 0,9750 | 0,7570 | 1,6210 | 1,0910 | 1,8420 | 1,7710 | 1,0700 | 0,8750 | 0,9320 | | 1,3990 | 0,7110 | | |
| 910 | -6,5350 | -5,9370 | -6,2390 | -5,9870 | -6,5620 | 0,3180 | 0,1820 | 0,0730 | 0,3460 | 0,0380 | -0,3620 | -0,5330 | -0,6730 | -0,5290 | -0,5930 | -6,1470 | -6,4140 | -6,2330 | -6,1640 | -5,8360 | 0,1010 | -0,2660 | 0,0350 | 0,1470 | -0,0060 | -0,1810 | -6,3780 | -0,3330 | -6,0190 | -0,3930 |
| 911 | | | | | | -1,0120 | | -1,1610 | -0,4140 | -0,2470 | | | | | | | | | | | -0,9490 | -0,8290 | | -0,5720 | | | -0,9970 | | -1,2270 | |
| 913 | | | | | | -1,1500 | | | -0,5820 | -0,5140 | | | | | | | | | | | -7,4370 | | -1,1190 | -1,2480 | -1,2000 | -7,3530 | -1,3400 | -7,3870 | -1,3100 | -7,3590 |
| 915 | | | | | | -7,2650 | | -1,4150 | | -0,7860 | | | | | | | | | | | | | | | | | -0,2870 | | | |
| 916 | | | | | | | | | | | | | | | | | | | | | | | | | -6,8530 | | | | | |
| 917 | | -0,5960 | | | | | -0,9600 | | | | | | | | | | | | | | -0,8850 | -0,4580 | -0,9500 | -0,8200 | | | -0,8070 | | | |
| 919 | | -1,4380 | -1,3770 | -0,8840 | | | | | | | -1,3670 | -1,1420 | -1,1310 | -1,0320 | -1,1050 | | -1,0160 | | -1,2560 | | -0,9790 | -0,8430 | | | | -1,2690 | -0,8300 | -0,8020 | | -1,3910 |
| 922 | | | | | | | | | | | | | | | -1,2880 | | | | | | | | | | | -1,1160 | -1,0080 | | -1,0540 | -1,0510 |
| 925 | | | | | | | | | | | | | | | | | | | | -0,8670 | | | | | | | | | | |
| 926 | | | | | | | | | | | | | | | | -0,8910 | | | | | | | | | | | | | | |
| 927 | 0,2230 | 0,3180 | 0,1680 | 0,9890 | 0,2060 | -0,1520 | -0,1400 | -0,1150 | -0,2050 | -0,1590 | | | | | | 1,0630 | 1,2910 | | 0,1420 | 0,3760 | -0,6450 | -0,5840 | | -0,2770 | -0,7620 | | | | | |
| 928 | -0,9470 | -0,8450 | -0,9890 | -0,6490 | -0,9320 | | | | | | | | | | | -0,6110 | -0,6960 | | -0,7880 | -0,9830 | | | | | | | | | | |
| 929 | 0,4410 | 1,0020 | 0,2440 | 0,4180 | 0,2590 | -1,0720 | -0,3330 | -0,7340 | 0,9010 | 0,4340 | | | | | | -0,0090 | -0,1570 | | -0,5250 | -0,7990 | -0,9800 | -1,1690 | | -0,8280 | -1,3070 | | | | | |
| 931 | -0,8260 | -0,7750 | -0,7430 | -0,0590 | -0,2870 | 0,0080 | -0,0490 | 0,0760 | 0,3780 | 0,6420 | | | | | | 0,3760 | 0,6280 | | | -0,5210 | 0,0770 | 0,3900 | | | 0,0270 | | | | | |
| 932 | | | | 0,7870 | 0,6060 | 0,9340 | 1,5960 | 0,9890 | 1,6530 | 1,1910 | | | | | | -0,1630 | 2,9770 | | | 0,7430 | 1,1590 | 0,2540 | | | -0,1790 | | | | | |
| 934 | | | | | | | -0,8050 | -0,8190 | 1,3270 | 1,3470 | | | | | | | | | | | -0,2110 | 0,5060 | | | | | | | | |
| 937 | | | | | | -0,8740 | -0,5770 | -0,5850 | 0,1540 | -0,0400 | | | | | | | | | | | -1,4410 | -0,3840 | | | | | | | | |
| 938 | | | | | | | | | -0,9340 | -1,0580 | | | | | | | | | | | | | | | | | | | | |
| 941 | | | | | | 1,6840 | 1,5190 | 1,6400 | 1,8460 | 2,0890 | | | | | | | | | | | | | | | | | | | | |
| 944 | | | | | | | | | | | | | | | | | | | | | -1,3140 | | | | | | | | | |
| 964 | | | | | | | | | | | | | | | | 0,0680 | | | | | | | | | | | | | | |
| 965 | | | -1,5280 | -1,0940 | -1,1120 | | | | | | | | | | | -1,8100 | | | | | | | | | | | | | | |
| 966 | | | | | | | | | | | | | | | | -0,9660 | | | | | | | | | | | | | | |
| 967 | -1,3700 | -1,2750 | -1,0350 | -0,4970 | -0,7830 | | | | | | | | | | | 1,0280 | -0,2760 | | | | | | | | | | | | | |
| 968 | -0,3870 | -0,8730 | -1,0140 | -0,0430 | -0,6000 | | | | | | | | | | | | | | | | | | | | | | | | | |
| 969 | 0,1410 | -0,4220 | -0,8590 | 0,2860 | -0,7140 | | | | | | | | | | | 0,1100 | -0,7900 | | | | | | | | | | | | | |
| 970 | 1,3470 | | | | | | | | | | | | | | | | | | | | | | | | | | | | | |
| 993 | | | | | | -1,6380 | | -1,5750 | -1,5300 | | | | | | | | | | | | | | | | | | | | | |
| 994 | | | | | | -1,4780 | -1,5260 | -1,4740 | -1,4030 | -1,1850 | | | | | | | | | | | | | | | | | | | | |
| 995 | | | | | | -1,1750 | -1,2790 | -1,2230 | -0,8710 | -0,8900 | | | | | | | | | | | | | | | | | | | | |
| 996 | | | | | | -0,4930 | -0,6110 | -0,5740 | -0,2690 | -0,3450 | | | | | | | | | | | | | | | | | | | | |
| 998 | | | | | | -0,0240 | -0,2390 | -0,2720 | -0,4370 | -0,2810 | | | | | | | | | | | | | | | | | | | | |
| 999 | | | | | | -0,0100 | 0,1130 | 0,1180 | 0,8110 | 0,9640 | | | | | | | | | | | | | | | | | | | | |
| 1000 | | | | | | -0,6080 | | | -0,3150 | -0,1590 | | | | | | | | | | | | | | | | | | | | |
| 1013 | | | | | | | | | | | -1,4460 | | -1,4330 | | -1,5450 | | | | | | | | | | | | | | | |
| 1014 | | | | | | | | | | | -1,3470 | | -1,3420 | | -1,4150 | | | | | | | | | | | | | | | |
| 1016 | | | | | | | | | | | -1,2180 | | -1,2630 | -1,4160 | -1,2580 | | | | | | | | | | | | | | | |

**Figure S4.** HSPred analysis. Related to Figure 5 only showing the interface forming residues with highlighted hotspot residues (red), non-hotspot residues (white), non-interface residues (blue), and non-existing residues in the reference alignment (colourless/black) (Tab. S11). All residues notThe coloring of the residue positions on the left is according to the C domain (green), C-A linker (blue), $A_{Core}$ (red), and $A_{Sub}$ (orange).

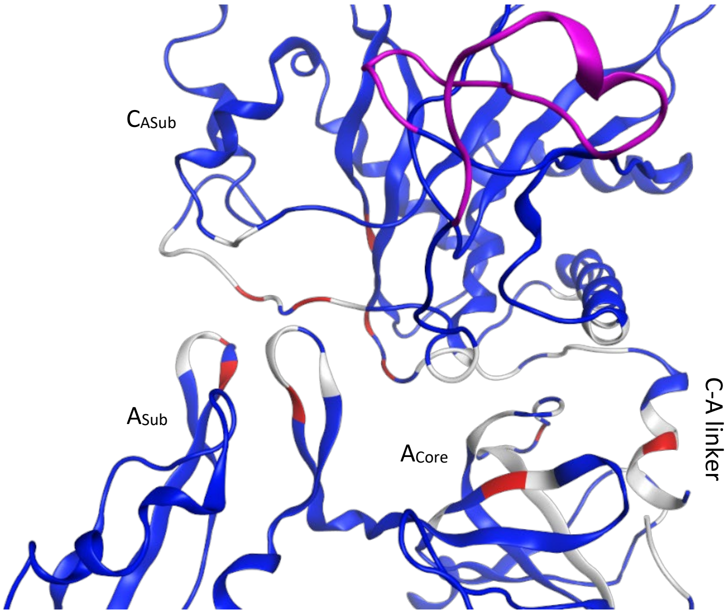


**Figure S5.** 2VSQ *Xsze* FclJ_C6-A6 homology model of HSPrad analysis with highlighted hotspot residues (red), non-hotspot residues (white), non-interface residues (blue), and non-existing residues in the reference alignment (colourless) (Tab. S11). The novel loop 'Chain II' marked in purple.

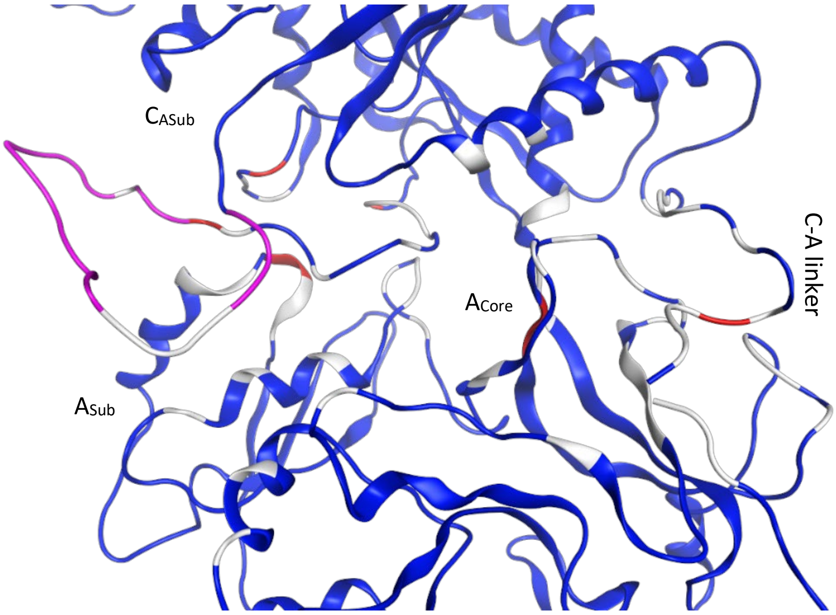


**Figure S6.** 4ZXH *Xsze* FclJ_C6-A6 homology model of HSPrad analysis with highlighted hotspot residues (red), non-hotspot residues (white), non-interface residues (blue), and non-existing residues in the reference alignment (colourless) (Tab. S11). The novel loop 'Chain II' marked in purple.

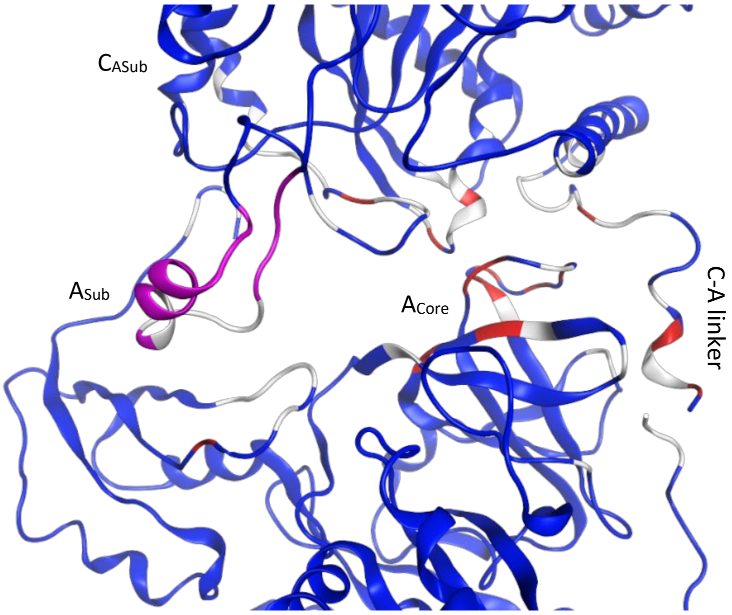


**Figure S7.** 5T3D *Xsze* FclJ_C6-A6 homology model of HSPrad analysis with highlighted hotspot residues (red), non-hotspot residues (white), non-interface residues (blue), and non-existing residues in the reference alignment (colourless) (Tab. S11). The novel loop 'Chain II' marked in purple.

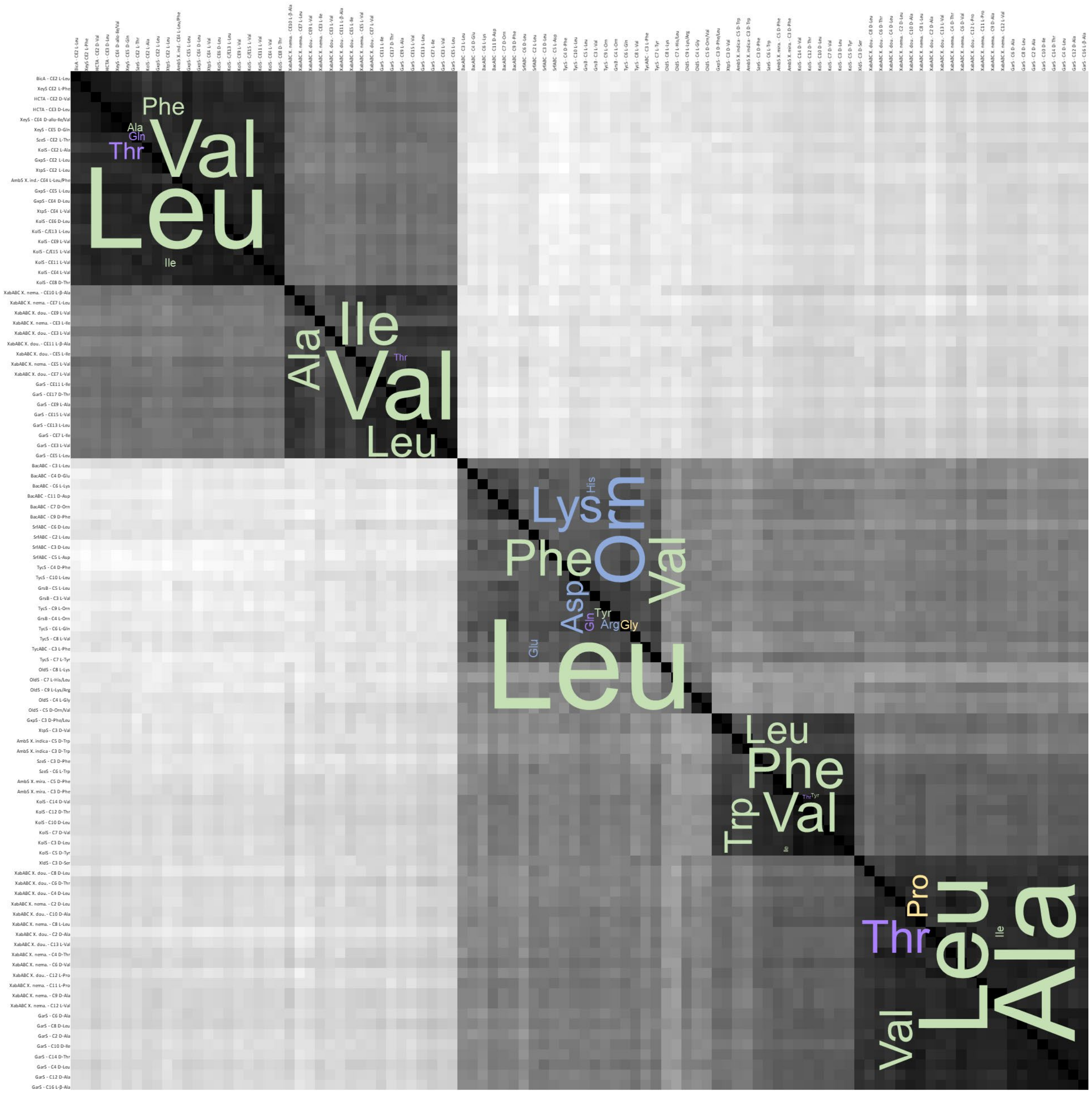


**Figure S8.** HeatMap of the Clustal Omega alignment of a selection of 100 $C_{ASub}$ domains. Higher similarity is represented by darker plot colors. Blocks of highest similarity (> 75 %) were further designated with amino acid WordMaps in three-letter abbreviation of the known corresponding A domain specificities. The color-code of the amino acids are referred to their electrically charged side chains (light blue), polar uncharged side chains (light purple), hydrophobic side chains (light green), and special cases (light yellow) at physiological pH (7.4).

### Supplementary Explanation

**Explanation S1.** *Crystal structure C-A template dynamics mark major contributions of the interacting residues*

The structural changes during the catalytic states are mainly manifested in the C-A interface by the conformational changes of the A domains’ larger N-terminal domain ($A_{Core}$) in respect to the smaller C-terminal subdomain ($A_{Sub}$). During the 30° rotation of the $A_{Sub}$ domain from the 2VSQ open conformation to the 4ZXH adenylate forming conformation in AOI1 Chain I and V are mainly interacting with the opposite $A_{Sub}$ highly flexible Chain XII-XVI. This goes along with a higher abundance of hot residues mainly in Chain XII & XV and the introduction of XIII in the interface paired with a stronger, shifted connection in the C domains' Chain I & V. Meanwhile, AOI2 loosen its connections in Chain III, and entirely in Chain IV, with their opposite $A_{Core}$ Chain IX. Likewise, in the $A_{Core}$/$A_{Sub}$ transition Chain VIII, but tighten a bit up in Chain XI.

The 140° body torsion from the 4ZXH adenylate forming conformation in the 5T3D thioester forming conformation makes Chain XVI the only connection of the $A_{Sub}$ to the C domain in AOI1 in unison with more hot residues in AOE2 and the $A_{Core}$ and $A_{Core}$/$A_{Sub}$ transition in Chain VIII-XI. Opposite the $A_{Core}$ on the C domain side, Chain III shows more hot residues, and Chain IV with parts of Chain V (while Chain V loses its stronger connected components) is reintroduced into the interface.

During these highly complex conformational changes, the linker interactions in AOI3 with the C domain only show slight alterations in the interface, with the weakest in the 5T3D.

Emphasis should be brought to the fact that the maintenance of the C-A platform in AOI2 is mainly determined by the interaction of the C domain Chain III, IV, and VI with the $A_{Core}$ Chain IX. Furthermore, the transition from $A_{Core}$ to $A_{Sub}$ occurs through Chain VIII&XI by interacting in AOI1&2.
